# Covalent activation of the C-type lectin DC-SIGN

**DOI:** 10.1101/2025.09.12.674704

**Authors:** Jonathan Lefèbre, Maurice Besch, Noémi Csorba, Kristóf Garami, Zoltán Orgován, Gitta Schlosser, Iris Bermejo, Péter Ábrányi-Balogh, György M. Keserű, Christoph Rademacher

## Abstract

Dendritic Cell-Specific Intercellular adhesion molecule-3-Grabbing Non-integrin (DC-SIGN) is a C-type lectin receptor expressed on antigen-presenting cells, crucial for pathogen recognition and immune modulation. The shallow and polar carbohydrate binding site of DC-SIGN presents challenges for ligand design. Here, we explored covalent modification targeting specific lysine residues as a novel strategy to modulate DC-SIGN function. Screening a lysine-targeted electrophilic fragment library using orthogonal functional assays identified two potent activators. Structural analyses via NMR spectroscopy, mass spectrometry and computational modeling confirmed structural perturbations of the carbohydrate recognition domain and revealed distinct mechanisms of activation. While both activators significantly enhanced DC-SIGN’s affinity for monosaccharide ligands, one compound induced oligomerization via covalent coupling and non-covalent secondary site interactions, whereas the other selectively modified lysine K373 directly within the primary carbohydrate-binding site. These findings demonstrate the potential of lysine-targeted covalent compounds as a novel therapeutic strategy for modulating DC-SIGN function and potentially C-type lectins in general.

**Table of contents:** 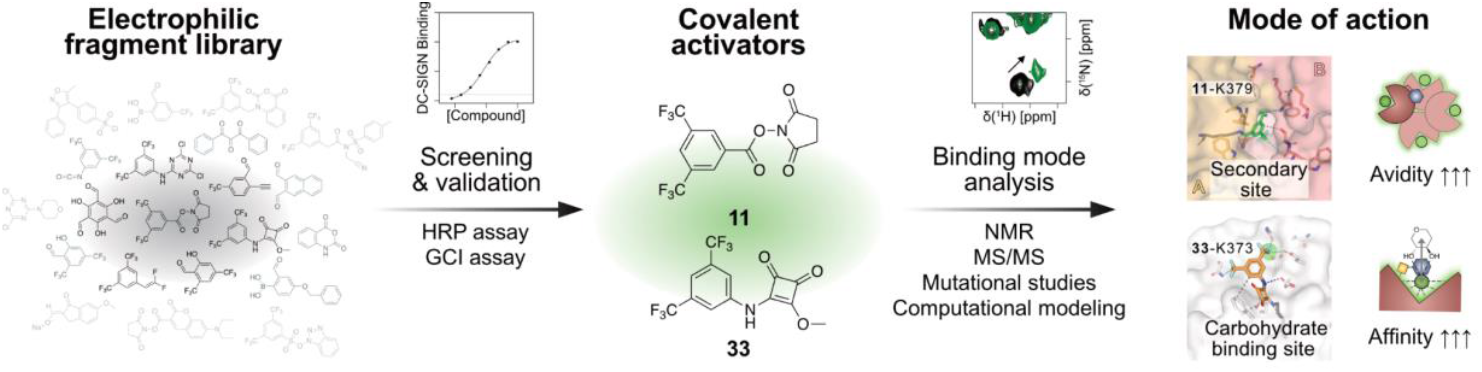

We introduce the first covalent activators of a C-type lectin. Using GCI, NMR, MS/MS and computational modeling, we delineate mechanisms from a functional electrophile-first screen on DC-SIGN that yields two modes: NHS-ester **11** modifies K379 to induce CRD oligomerization via a secondary site, and squarate **33** modifies K373 in the carbohydrate site to strengthen glycan binding.

## Introduction

DC-SIGN (Dendritic Cell-Specific Intercellular adhesion molecule-3-Grabbing Non-integrin) is a Ca^2+^-dependent lectin expressed on antigen presenting cells, playing a key role in pathogen recognition, immune modulation, and cell adhesion ^[1]^. It recognizes carbohydrate patterns on various pathogens and self-antigens, facilitating their capture and internalization, making it an attractive therapeutic target. However, targeting DC-SIGN’s carbohydrate-binding site with small molecules remains notoriously difficult due to its inherent low druggability and the typically weak affinity of monosaccharide-based ligands ^[2]^. On the one hand, efforts to develop ligands have predominantly focused on carbohydrate-based ligands, often depending on multivalent presentation on various supports to increase potency ^[2a, 3]^. On the other hand, identification of several secondary sites and moderate hit rates in fragment screening campaigns have suggested DC-SIGN to be amenable to fragment-based drug discovery (FBDD) approaches ^[4]^.

As an alternative strategy, covalent ligand approaches have emerged as a promising method to efficiently modulate challenging targets and to enhance both potency and selectivity ^[5]^. By forming irreversible or prolonged interactions with target proteins, covalent ligands can overcome the limitations of conventional affinity-driven ligand design. In addition, covalent fragment approaches combine the advantages and effectiveness of FBDD and target identification by covalent labeling ^[6]^. Covalent modulation can be achieved by ligand-first or electrophile-first approaches. In the former case, a high affinity ligand is equipped with a well-positioned electrophilic functional group, referred to as warhead. In the latter approach, a path commonly followed for challenging targets without ligands, small electrophilic fragments are screened to find starting points for FBDD approaches ^[7]^.

In this study, we pursued an electrophile-first approach by designing and screening a Lys-targeted covalent fragment library against DC-SIGN. We identified and characterized the first covalent activators of DC-SIGN, providing novel insights into mechanisms of glycan-binding enhancement through covalent modification. Detailed biophysical and structural investigations elucidate distinct modes of action, demonstrating how covalent targeting of lysine residues can effectively modulate glycan recognition by DC-SIGN through induction of oligomerization or by directly affecting monosaccharide recognition. These findings open new avenues for exploiting covalent strategies in the development of DC-SIGN-targeted but also C-type lectin-targeted small molecules in general.

## Results & Discussion

### Library design and chemistry

We have compiled a 34-membered library of reactive fragments based on known covalent warheads reported to target lysine residues in proteomic studies or in targeted covalent inhibitor (TCI) development (Table S1) ^[6, 8]^. Seventeen fragments were commercially available while the other half of the fragments were synthesized (for the details of the syntheses, see Supporting Information). First, we evaluated the reactivity of the library members with *N*-acetyl lysine in a HPLC-MS-based kinetic assay, measuring pseudo-first-order rate constants and the formation of the corresponding adducts formed at pH 10.2. At this pH, the ε-amine of lysine would be deprotonated which mimics the intrinsic reactivity of Lys residues at protein binding sites, where its pK_a_ can range from 5 to 11 depending on the microenvironment ^[6, 9]^ (Figure 1a, Table S2). The results confirmed that the library covers a wide range of reactivities (Figure 1b). We found 14 fragments reacting immediately (t_1/2_ < 5 min), while 5 and 4 showed reactivity under 20 min or 90 min. The half-life of 11 fragments was over 24 h containing 3 mildly reactive compounds (>48 h). The aqueous stability of the fragments was then evaluated under physiological conditions (pH 7.4). Among the highly reactive compounds six showed >80% stability after 1 h, while for five fragments the stability was <50%. Thirteen of the 20 less reactive fragments showed acceptable stability (>80%), and only three were found to be unstable (<50%). Based on the wide range of reactivity, we assumed that lysines in proteins showing diverse nucleophilicity can be targeted (Figure 1b, Table S2). Having many highly reactive warheads supports the diversity in targeting lysines with low reactivity, while low reactivity warheads might provide specific labeling.

**Figure 1.**
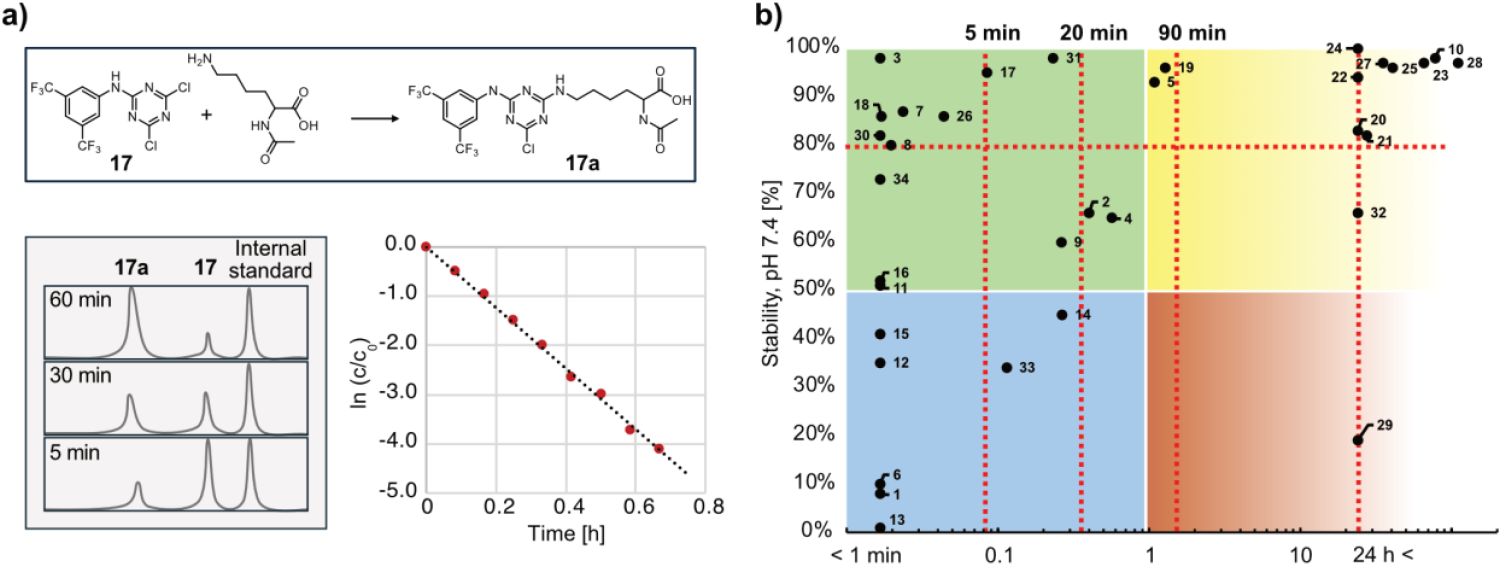
The electrophilic fragment library displays a wide range of reactivities. **a)** Schematic representation of the reaction of **17** with *N*-acetyl lysine. Time-resolved HPLC chromatograms for the reactivity assay and plotting the decrease of the fragment for the ln(k) and t_1/2_ determination by linear regression. **b)** Correlation between buffer stability and *N*-acetyl lysine reactivity. The y-axis represents the percentage of compound remaining after 1 h, indicating stability under physiological conditions. The x-axis shows the reactivity of each compound, expressed as the half-life (t_1/2_) derived from pseudo-first-order kinetic measurements of *N*-acetyl lysine conjugation. The plot illustrates the relationship between compound stability and reactivity, with shorter half-lives indicating faster reaction rates. Red dashed lines correspond with thresholds mentioned in the text (Table S2).

### Functional screening identifies DC-SIGN activating electrophiles

The first screening assay selected is based on the multivalent interaction of the glycoprotein horseradish peroxidase (HRP) with the carbohydrate binding site of the tetrameric DC-SIGN ectodomain (ECD). The assay has been previously utilized to screen non-covalent inhibitors as well as to evaluate the inhibitory or activating mutations on the DC-SIGN carbohydrate recognition domain ^[10]^. To test the covalent compounds, the purified protein was immobilized on plates and incubated with 0.5 mM of compound overnight at 4 °C or room temperature. Activity of the modified protein was then assessed by addition of HRP followed by detection of its peroxidase activity (Figure 2a).

**Figure 2.**
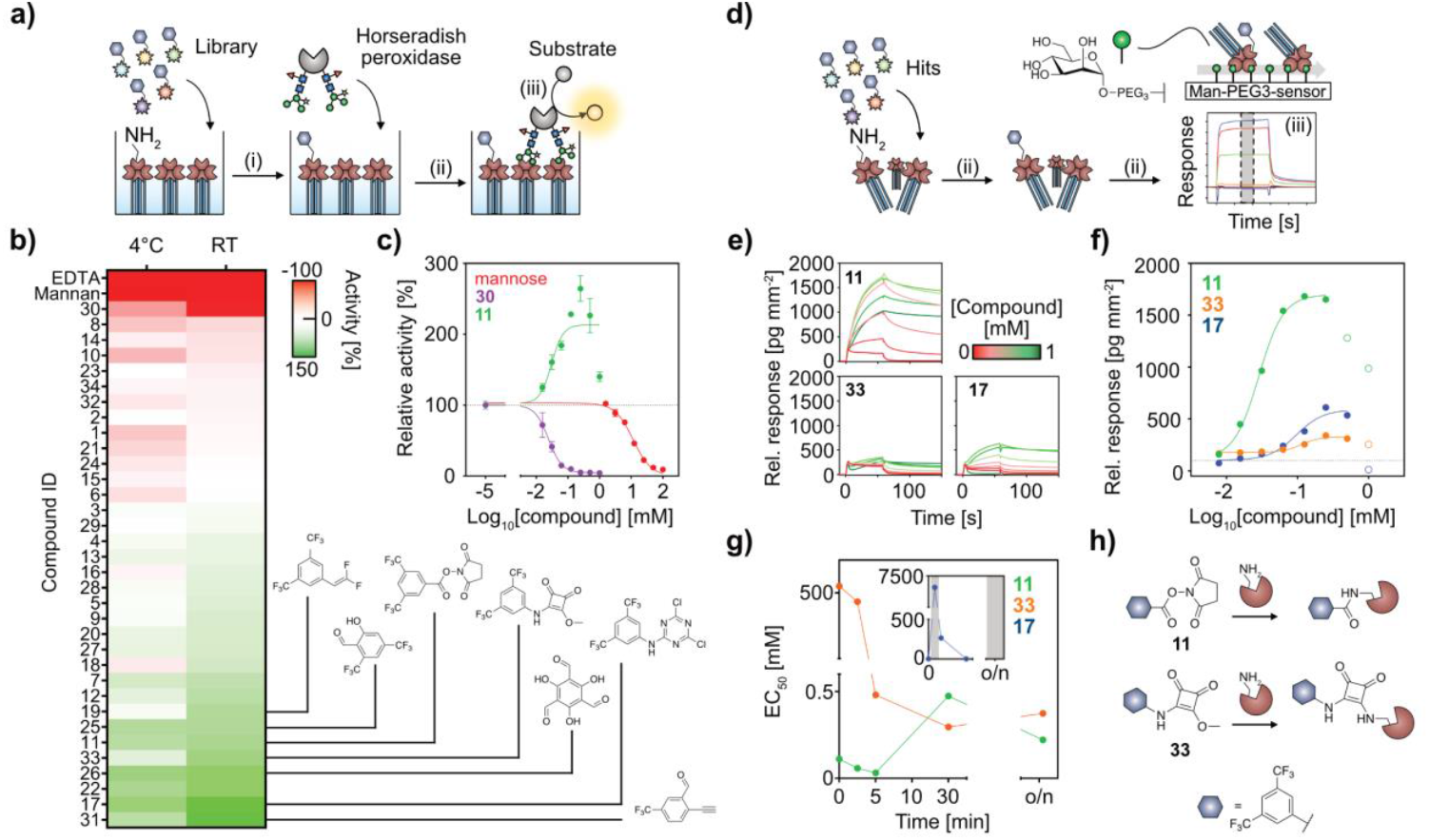
Functional screening and validation of the covalent warhead library against DC-SIGN ECD. **a)** The HRP assay was used to screen the electrophilic fragments at single concentrations in dose-response experiments. Plate-immobilized DC-SIGN ECD is incubated with 0.5 mM of each compound overnight (i). After washing, HRP is added to the wells (ii). HRP interacts with DC-SIGN ECD via its endogenous glycosylation. The amount of HRP bound is evaluated via its peroxidase activity (iii). **b)** Heatmap displaying screening results for incubation at 4°C or at RT. EDTA and mannan were used as controls for the DC-SIGN-glycan interaction. Overall, more electrophilic fragments activating DC-SIGN were found. Structures of activators displaying an increase in DC-SIGN binding activity of ≥ 50% relative to the DMSO control are shown. Results represent the mean of three biological replicates. **c)** Example dose-response curves after overnight incubation at RT, showing mannose as control inhibitor (IC_50_ = 11.4 mM), **30** as inhibitor (IC_50_ = 0.02 mM) and **11** as activator (EC_50_ = 0.03 mM). Dose-response curves of other hit compounds are shown in figures S1 and S2. **d)** A GCI assay was established to orthogonally validate hit compounds in dose-response experiments. DC-SIGN ECD in solution is incubated with each compound (i). Modified DC-SIGN is injected over a quasi-planar GCI sensor modified with mannose-PEG3-amine, allowing for multivalent protein-gylcan interactions (ii). Binding is evaluated from resulting sensorgrams using response at equilibrium (iii). **e)** Sensorgrams of DC-SIGN ECD modified with validated activators **11, 33** and **17** showing drastic change in relative response and reduced off rates of the interaction. **f)** EC_50_ fit of binding response from sensorgrams in **(e)** yielded micromolar activities for the hit compounds **11** (EC_50_ = 0.03 mM), **33** (EC_50_ = 0.1 mM), **17** (EC_50_ = 0.1 mM). Data points without filling were not used for fitting. **g)** Dose-response curves of the validated hit compounds after varying incubation times using the HRP assay, indicated clear time-dependency for **11** and **33**, but not for **17**. Grey areas indicate timepoints were the EC_50_ could not be fitted reliably. **h)** The two validated hit compounds **11** and **17** share the same non-covalent 3,5-bis(trifluoromethyl)phenyl moiety but yield different reaction products.

Using a cutoff of ∼50% activity change compared to the DMSO control, we identified only **30**, a naphthalene-2,3-dicarbaldehyde *ortho*-phthalaldehyde (OPA)-type fragment, as a potential inhibitor, whereas eight compounds (**11, 17, 19, 22, 25, 26, 31, 33**) showed ≥ 50% activation of DC-SIGN (Figure 2b) ^[11]^. While the almost exclusive identification of activators is surprising, we could exclude bias of the assays to identify only activators using mannan, mannose and EDTA as known inhibitors of the DC-SIGN carbohydrate binding site as controls. Moreover, temperature-dependent increase in activity of all selected compounds, suggested a covalent mechanism underlying the compound activities. We further confirmed **30** as an inhibitor and seven of the eight activators (salicylaldehydes **25** and **26**, squarate **33**, difluorostyrene **19**, dichlorotriazine **17**, formylacetylene **31**, *N*-hydroxysuccinimide (NHS)-ester **11**) in dose-response experiments, revealing micromolar activities (Figures 2c, S1 and S2). The fragments identified as hits react through various mechanisms, i.e. through acylation (**11**), conjugate addition-elimination (**33**), imine formation (**25, 26, 31**), substitution (**17, 19, 33**).

To orthogonally validate activity of these compounds in solution, we established a grating-coupled interferometry (GCI) assay, in which altered affinity of DC-SIGN for its monosaccharide ligand can be evaluated based on binding to mannose-PEG3-amine coupled to the sensor chip (Figure 2d). Upon injection, DC-SIGN ECD showed sufficient binding to the chip with affinities similar to previous surface plasmon resonance setups utilizing mannosylated BSA on the sensor surface (Figure S3a) ^[12]^. In control experiments with mannose, the assay yielded IC_50_s comparable to published data and results from titrations in the HRP assay, demonstrating dependency of the interaction on the carbohydrate binding site (Figure S4a) ^[13]^.

For the covalent hits, DC-SIGN ECD was preincubated with the compounds overnight and quenched by the addition of *tris*(hydroxymethyl)aminomethane (Tris) in 5-fold excess relative to the ligand prior to injection (Figure 2d). While the inhibitor **30**, as well as activators **26, 25, 19** and **31** precipitated the protein or showed no activity compared to the DMSO control, compounds **11, 33** and **17** activated binding to the chip surface (Figures 2e, f, S5 and S6). Notably, although no precipitation upon incubation with **11, 33** or **17** was observed, compound concentrations exceeding 0.25 mM led to reduction in binding response, indicating covalent modification at additional less reactive sites. Yet, saturation of the binding response by all three compounds allowed for fitting of binding parameters and yielded EC_50_s comparable to those obtained in the plate-based assay. Hill slopes > 1 of the binding isotherms indicated positive cooperative effect induced by the compounds. Together with the observation of a drastic change in off rates, indicating the compounds to modulate kinetics of the DC-SIGN ECD-carbohydrate interaction, we hypothesized the binding response to result from an increase in affinity or avidity or both (Figure S7).

As DC-SIGN carbohydrate recognition is not limited to simple monosaccharides and establishes additional contacts outside of the principal Ca^2+^ dependent carbohydrate binding site, as utilized in the case of larger oligosaccharides being present on HRP, we also tested DC-SIGN modified with covalent adducts (**11, 33, 17**) on a sensor chip coupled to Man(α1,2)Man-PEG3-amine (Figures S3b and S4b) ^[14]^. We obtained similar results as for the mannose-PEG3-amine sensor, suggesting that the activity of the covalent compounds does not originate from modulating additional contacts to larger glycans outside of the canonical carbohydrate-binding site (Figure S8 and S9).

Finally, to confirm the covalent activity of the three hits, we quenched the reaction at different time points in our plate-based assay and detected activity by binding to HRP. This revealed clear time-dependency of the EC_50_ of **33** and indicated irreversible modification of the protein. **11** showed fast reaction followed by reduced activity at incubation times exceeding ten minutes. For **17**, a time-dependent increase in response was observed that was however increasingly linear suggesting unspecific reaction at longer incubation times (Figure 2g). Consequently, we excluded **17** from further investigation.

Taken together, our functional screening and validation of the reactive fragment library against DC-SIGN ECD identified and validated two covalent activators and no true inhibitors. Considering the different warhead chemistries as well as products and selective reactivity of hits **11** and **33** for specific Lys sidechains, we speculated a general tendency of covalent modification of Lys residues to activate DC-SIGN (Figure 2h).

### Covalent activators modulate glycan binding kinetics of the carbohydrate recognition domain via different mode-of-actions

The neck domain of DC-SIGN ECD contains several Lys involved in coiled-coil interactions important for tetramerization of the receptor (Figure 3a) ^[15]^. To exclude that the activity of the hit compounds relies on unspecific reaction with the neck domain instead of the carbohydrate recognition domain (CRD), we transferred our GCI assay to the isolated DC-SIGN CRD.

**Figure 3.**
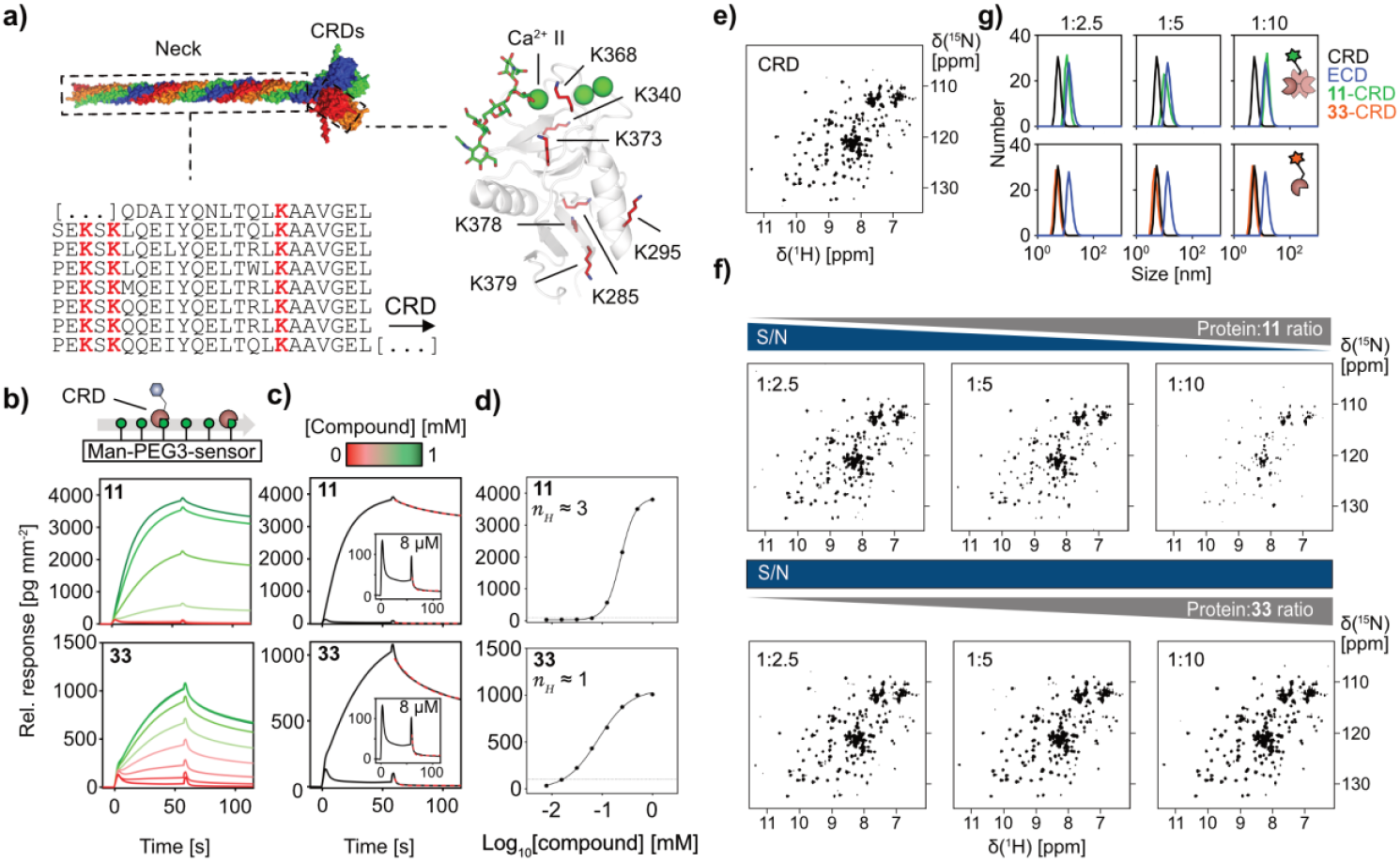
Electrophilic fragment hits activate DC-SIGN CRD via different mechanisms. **a)** Model of the DC-SIGN ECD tetramer, colored by monomer. The Sequence of the neck domain with lysines highlighted in red and the CRD in complex with an oligosaccharide (PDB: 1K9I) with lysines as red sticks are shown. **b)** The GCI assay implemented for DC-SIGN ECD was transferred to the CRD to test the effect of activators **11** and **33** on the monovalent DC-SIGN-glycan interactions. **c)** Off-rates (k_off_) of the CRD-mannose-PEG3 interaction are drastically reduced for both **11** and **33**-modified DC-SIGN CRD, when comparing low (0.008 mM; k_off_,_**11**_ ≈ 0.25 s^-1^, k_off_,_**33**_ ≈ 0.41 s^-1^) and high (1 mM; k_off_,_**11**_ ≈ 0.03 s^-1^, k_off_,_**33**_ ≈ 0.04 s^-1^ concentrations of the compounds. **d)** The activity of **11** was reduced compared to experiments with the ECD (EC_50_ = 0.2 mM) and the hill slope suggested cooperativity effects. **33** showed comparable activity (EC_50_ = 0.1 mM) on the CRD as on the ECD and a hill slope ∼1. **e)** ^1^H-^15^N HSQC NMR spectrum of ^15^N-labeled wildtype DC-SIGN CRD in the absence of compound. **f)** ^1^H-^15^N HSQC NMR spectra of DC-SIGN CRD modified with **11** at increasing ratios show reduced S/N, while **33** does not change spectral quality. **g)** Comparison of the protein hydrodynamic radius in DLS upon modification indicates and increase in size of **11**-modified DC-SIGN CRD. At a protein-**11** ration of 1:10, DC-SIGN CRD has a similar size as the unmodified DC-SIGN ECD, suggesting **11** to induce oligomerization. In accordance with ^1^H-^15^N HSQC NMR data, **33**-modified DC-SIGN CRD displays no increase in size.

Similar to experiments with the ECD, mannose titration revealed monovalent IC_50_s in the expected range, and an increase in affinity for the Man(α1,2)Man surface compared to the mannosylated one, both confirming the carbohydrate specificity of the interaction (Figures S10 and S11). Using the same setup as for the ECD, dose-response experiments with the covalent hits **11** and **33** confirmed the activating potential of the compounds (Figure 3b). No reduction in binding response was observed at higher concentrations, suggesting that either additional reaction sites at the neck domain or changes in the relative orientation of the CRDs in the tetramer upon reaction could lead to inhibition of carbohydrate binding. In line with observations from experiments with the ECD, we observed reduced off rates upon modification of the CRD with the compounds (Figure 3c). As the isolated CRD cannot oligomerize in the absence of the neck domain, we reasoned that reduced off-rates could point towards modulation of the CRD-monosaccharide affinity ^[16]^. This was further supported by a decreased dissociation constant (K_D_) for the sensor surface for both **11** and **33**-modified DC-SIGN CRD (Figures S12 and S13). When tested on the Man(α1,2)Man surface, activity of **11** and **33** was only slightly increased, further supporting a model in which changes in monosaccharide binding instead of oligosaccharide binding dominate the mode of action of the compounds (Figure S14).

Interestingly, **11** showed a significantly higher EC_50_ and a similar hill slope of > 1 when compared to experiments with the ECD (Figure 3d). Apart from indicating the activity of **11** to be partially dependent on the presence of the neck domain, this suggested cooperativity of the **11**-modified CRD interacting with the chip. In contrast, the EC_50_ of **33** was similar to experiments with the ECD and a hill slope ≈ 1 suggested **33** to specifically modulate the CRD in both the tetrameric ECD and the isolated CRD.

Next, we aimed to elucidate binding sites and structural perturbations induced by **11** and **33** interacting with DC-SIGN CRD. We incubated ^15^N-labeled DC-SIGN CRD with 0.25 mM of each compound, followed by dialysis to remove unreacted ligand. We recorded ^1^H-^15^N HSQC NMR spectra of ^15^N-labeled wildtype DC-SIGN CRD in the absence of ligand to establish a reference for subsequent comparison (Figure 3e). This baseline allowed us to attribute later spectral changes specifically to the non-covalent moieties of **11** or **33**, anchored through covalent attachment at the modified lysine residues ^[17]^.

Strikingly, ^1^H-^15^N HSQC NMR spectra **11** and **33**-modified DC-SIGN CRD wild type revealed drastic differences. Spectra of **11**-modified DC-SIGN CRD showed significantly reduced signal-to-noise (S/N) compared to the unmodified and **33**-modified protein (Figure 3f). As no precipitation was observed visually, we reasoned that an increase in size and therefore an increase in relaxation rates of the protein upon reaction with **11** is the origin of the reduction in S/N. This was confirmed by dynamic light scattering (DLS), showing a clear shift towards a population of larger particles (Figure 3g). Since the reduction in S/N and the population of larger particles increased with the ligand-to-protein ratio, we concluded that compound **11** induces oligomerization of DC-SIGN in a concentration-dependent manner. The size of the **11**-CRD particles was close to those of the DC-SIGN ECD tetramers, and no soluble aggregates were observed, suggesting a distinct oligomerization pathway.

Taken together, our GCI and HSQC NMR assays suggested distinct mechanisms underlying DC-SIGN activation by **11** and **33**. While **11** triggered formation of CRD oligomers, **33** showed selective covalent modification without altering the protein monomeric state.

### Activation by 33 relies on covalent coupling to K373 at the carbohydrate binding site

To study the covalent and non-covalent interaction of **33** with the DC-SIGN CRD we calculated chemical shift perturbations (CSPs) relative to the wild type protein from ^1^H-^15^N HSQC NMR spectra, informing about changes in the chemical environment of the ^15^N labeled backbone amide of each residue ^[18]^.

We observed only one new cross-peak upon modification with **33**, supporting reaction with a single Lys sidechain (Figure 4a). In line with this, CSPs in residues of the long loop (E349, N350, V351), β-strands 3 and 4 (E358, D366) and adjacent loops (F313, A372) suggested a localized effect of covalent modification by **33** (Figure 4b and 4c). Intriguingly, affected residues such as E349, N350, V351 and D366 are directly involved in coordinating Ca^2+^ and binding monosaccharides, or as in the case of F313 and E358, form the extended recognition surface for the interaction with oligosaccharides ^[14b, 19]^. Computational analysis of Lys solvent-accessible surface areas (SASA) and predicted pK_a_ values identified K368 as solvent-exposed and K373 as more buried, though both exhibited similar predicted nucleophilicity (Table S3). Accordingly, we hypothesized whether covalent modification of nearby K368 or K373 could sterically allow the non-covalent moiety to support either monosaccharide binding or affect Ca^2+^ coordination (Figure 4d).

**Figure 4.**
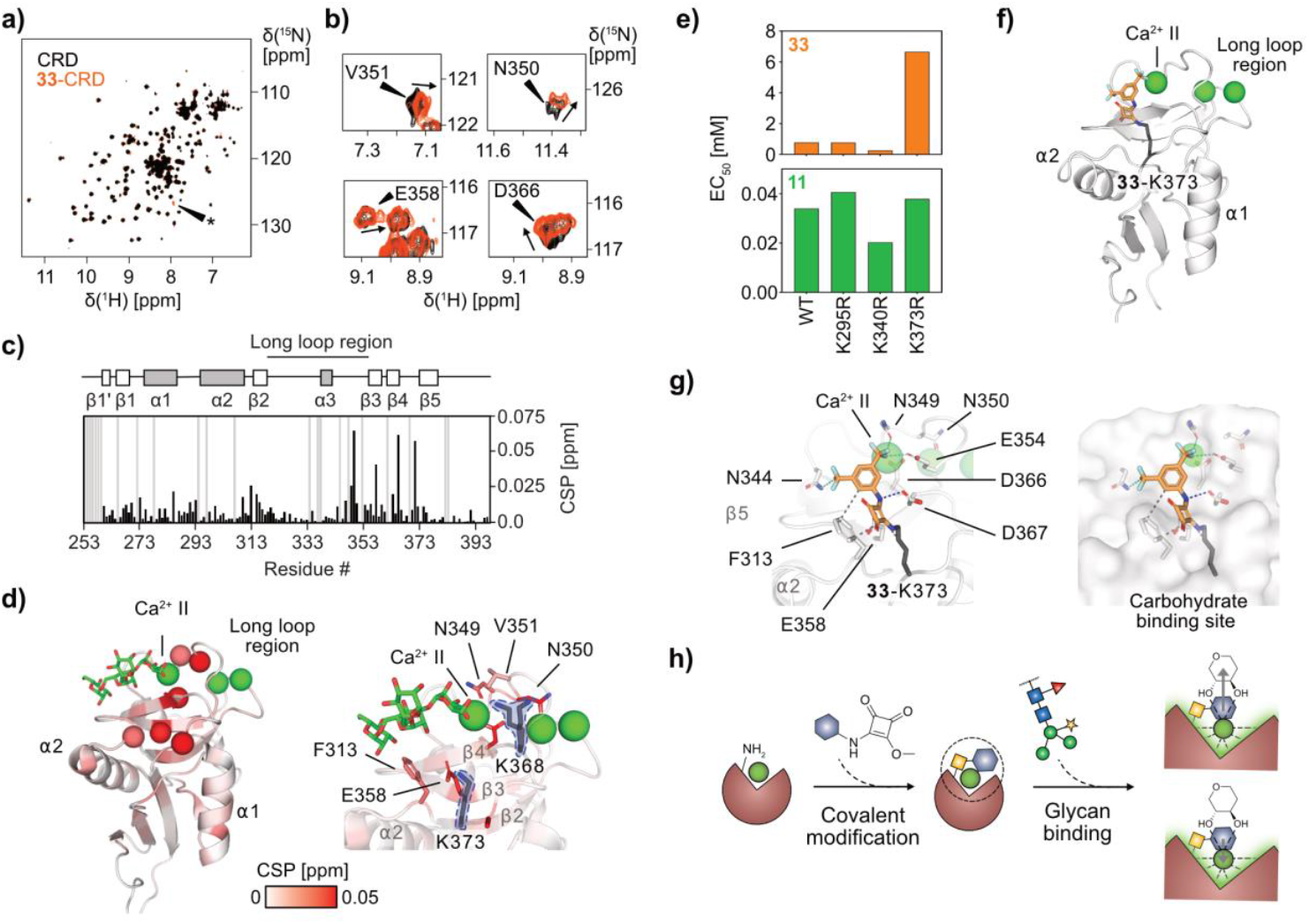
Compound **33** activates DC-SIGN by modifying residue K373 at the carbohydrate binding site. **a)** Superposition ^1^H-^15^N HSQC NMR spectra of unmodified ^15^N-labeled and **33**-modified DC-SIGN CRD wild type. **b)** Examples of residues showing CSPs compared to the unmodified protein. **c)** and **d)** CSP mapping onto the X-ray crystallographic structure of DC-SIGN CRD (PDB: 1SL4) indicate **33** to perturb the long loop region and β-strands 3 and 4 forming the Ca^2+^-dependent carbohydrate binding site of DC-SIGN. Cα atoms of residues showing CSP > 0.025 ppm are shown as spheres. Magnification of the region around suggests K368 or K373 as major modification site. **e)** Comparison of EC_**50**_ values obtained for the Lys mutants in HRP binding assays. The K373R mutant shows decreased activity upon reaction with **33**. For compound **11** no differences in EC_50_ values were observed. **f)** Covalent docking of **33** to K373 predicts binding mode of **33** at the carbohydrate binding site of DC-SIGN. **g)** Close-up of the predicted binding mode of **33** docked to DC-SIGN. Both, the 3,5-bis(trifluoromethyl)phenyl moiety and the squarate linker, interact with residues of the extended (F313, E358) and the canonical carbohydrate binding site (N349, N350, E354). **h)** Schematic representation of the mode-of-action of **33** activating DC-SIGN. Upon covalent modification of K373, **33** either directly supports interaction with a glycan (top) or indirectly by supporting Ca^2+^ binding (bottom). The 3,5-bis(trifluoromethyl)phenyl moiety and the squarate linker are colored blue and yellow, respectively.

To validate our hypothesis on **33** modifying K368 or K373 in the carbohydrate binding site we mapped labeling sites in chymotrypsin and ProAlanase digested peptides by LC-MS/MS. This proved modification of K373 on peptide NDDKCNLA**K**^373^F (Figures S15 and S16, Table S4).

Considering the non-covalent moiety of **33** to represent a low-complexity fragment, likely with low affinity, we reasoned that mutation of K373 to a non-nucleophilic amino acid should deplete activity of the compound. Since a K373N mutation has been previously shown to alter binding of DC-SIGN to glycans, we inserted a soft K373R mutation into the ECD construct, consequently removing nucleophilicity while maintaining positive charge in this site ^[14a, 20]^. We also generated control mutants K295R and K340R that we hypothesized would have no effect on the activity of **33**. Successful expression and purification of the mutants via mannan affinity chromatography suggested all mutants to be folded and in an active state. To evaluate a potential change in EC_50_ we incubated the mutants alongside the wildtype with **33** and measured activity based on binding to HRP. Strikingly, **33** only showed drastically reduced activity after incubation with K373R, while K295R and K340R displayed EC_50_s similar to that of the wild type (Figure 4e). Notably, compared to the wild type K373R showed reduced binding to HRP. To exclude that depletion of activity is resulting from this change in glycan binding, we also measured activity **11**-modified mutants. No change in EC_50_ was observed, further confirming selectivity of **33** for K373 (Figure 4e).

Finally, to rationalize the binding mode of **33** at the carbohydrate binding site of DC-SIGN we applied a induced fit docking protocol using CSPs from our ^1^H-^15^N HSQC NMR experiments and determined that the warhead of **33** is located close enough to K373 (3.5 Å N(H_2_)-C(OMe)) for the formation of a covalent bond, that was also confirmed by its independent covalent docking (Figure 4f). In the best-scored docking pose, the 3,5-bis(trifluoromethyl)phenyl moiety and the squarate linker, interact with residues of the extended and the canonical carbohydrate binding site (Figure 4g).

Taken together, the combination of HSQC NMR, LC MS/MS, mutational analysis and docking suggested **33** to covalently modify K373 acting either directly on the DC-SIGN-glycan interaction or indirectly via modulating Ca^2+^ coordination (Figure 4h).

### 11 induces oligomerization via non-covalent binding to a secondary site

As the NHS-ester warhead of **11** cannot covalently cross-link the protein, we speculated whether a non-covalent interaction at a neighboring CRD could support the formation of oligomers. We have previously identified a secondary site in DC-SIGN, which was specifically targeted by phenol and compounds resembling the non-covalent moiety of our hits (Figure 5a) ^[4a, 4d]^.

**Figure 5.**
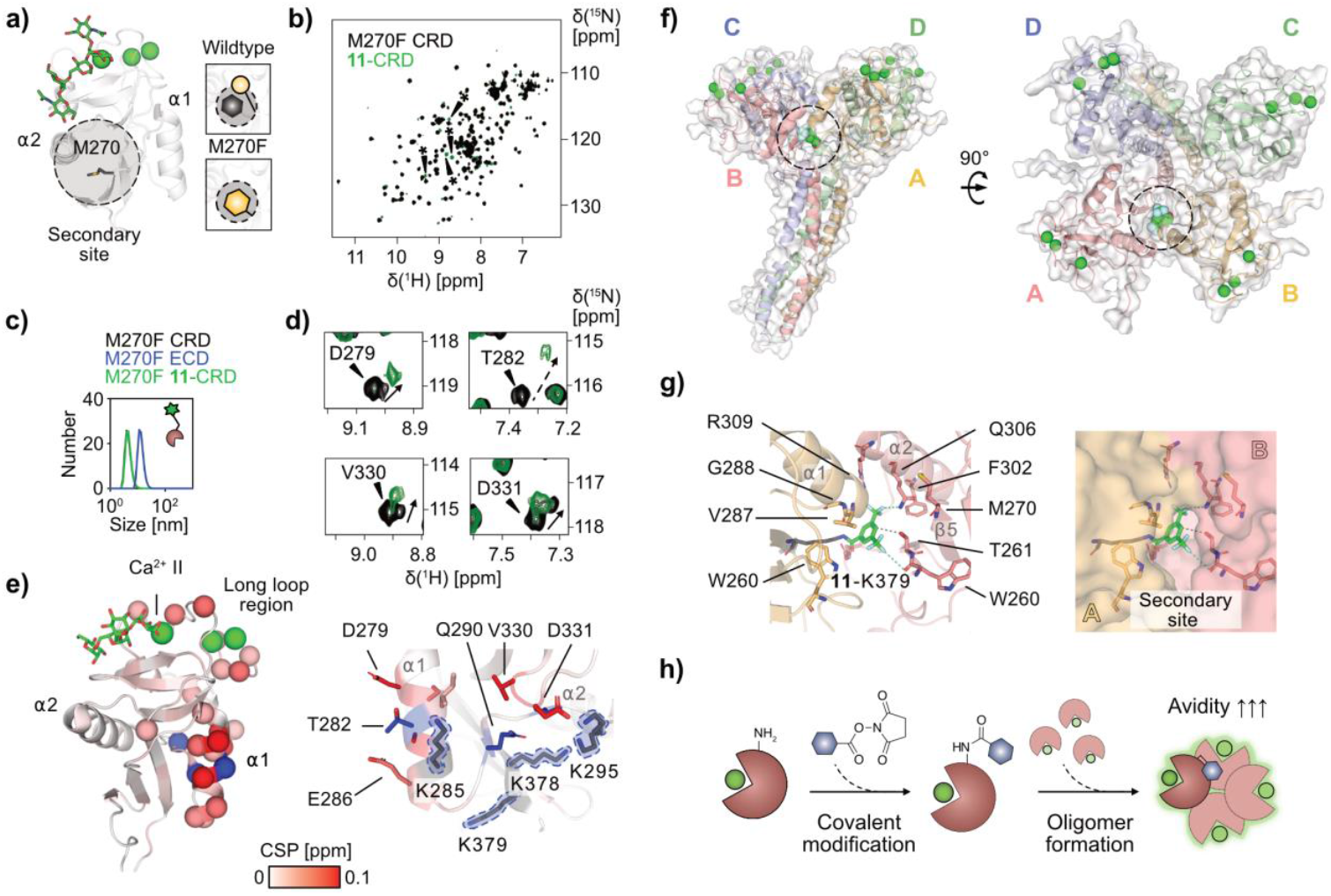
Compound **11** induces oligomerization by modifying K379 and interacting with a secondary site. **a)** Model of the secondary site below α-helix 2, centered at residue M270, previously found to interact with phenol and drug-like fragments. The secondary site can be blocked by inserting a M270F mutation ^[4d, 10b]^. **b)** Superposition ^1^H-^15^N HSQC NMR spectra of unmodified ^15^N-labeled and **11**-modified DC-SIGN CRD M270F. No reduction in S/N was observed upon reaction with **11** at protein-**11** ratios of 1:10. New cross-peaks marked by stars appear upon modification with **11**, indicative of the formation of several new amide bonds. **c)** Size comparison of unmodified and **11**-modified DC-SIGN CRD M270F and DC-SIGN ECD M270F confirms absence of oligomerization due to blocking of the secondary site. **d)** The ^1^H-^15^N HSQC NMR spectra of **11**-modified DC-SIGN CRD M270F enable analysis of CSPs due to improved S/N compared to wild type **11**-modified DC-SIGN CRD. Examples of residues showing large CSPs compared to the unmodified protein. **e)** CSPs mapped onto the X-ray crystallographic structure of DC-SIGN CRD (PDB: 1SL4), show clustering of larger CSPs at α-helix 1, located opposite to the secondary site. Cα atoms of residues showing CSP > 0.025 are shown as spheres. Blue spheres indicate vanishing or severely broadened resonances upon modification with **11**. Magnification of the region around α-helix 1 reveals several lysine residues in vicinity of the most perturbed residues. **f)** Modeling of the complex of tetrameric DC-SIGN including two C-terminal neck repeats with compound **11** ^[21]^. Covalent coupling at CRD A positions the ligand distal from the carbohydrate binding site at the interface to CRD B, supporting oligomerization. **11** is shown as spheres. The interaction site in CRD B is highlighted with a dotted circle. Ca^2+^ ions are shown as green spheres. **g)** Close-up of the interface between CRD A and B. Covalent modification of K379 directs the 3,5-bis(trifluoromethyl)phenyl moiety of **11** to interact with a previously described secondary site in CRD B ^[4d]^ .The ligand directly interacts with T261 and Q306 of the secondary site, facilitating protein-protein interactions supporting oligomer formation. **h)** Schematic representation of the mode-of-action of **11** activating DC-SIGN. Upon covalent modification of K379, the non-covalent moiety of **11** supports oligomerization allowing the protein to interact with glycans with higher avidity. The 3,5-bis(trifluoromethyl)phenyl moiety is colored in blue.

To test whether the oligomerization could be induced by non-covalent binding of **11** to this site, we repeated our ^1^H-^15^N HSQC NMR experiments with the ^15^N labeled CRD M270F mutant that was shown to block phenol binding to DC-SIGN (Figure 5a) ^[4d, 10b]^. In line with our hypothesis, no decrease in S/N was observed and DLS further confirmed lack of oligomerization of the M270F CRD (Figure 5b and 5c). In the absence of oligomer formation, we were able to calculate CSPs induced at the site of covalent modification by **11** (Figure 5d). Mapping of CSPs on the X-ray structure of DC-SIGN revealed clustering of larger perturbations in α-helix 1 and the loop connecting α-helix 1 and α-helix 2 (Figure 5e). As blocking of the secondary site restricts specific non-covalent interactions, we argued that these CSPs result from covalent modification of one or several of the four Lys are located in this region (K285, K295, K378 and K379) (Figure 5e). This was supported by additional peaks appearing in the spectra, resulting from the formation of new amide bonds at the side chains of modified Lys residues (Figure 5b).

To define the reaction site in more detail, we digested the labeled protein by chymotrypsin and ProAlanase, revealing **11** to label K379 on peptide KFWICK**K**^379^S (Figures S17 and S18, Table S5). This is in agreement with the prediction that K379 is one of the most nucleophilic lysine available for covalent targeting (Table S3). Notably, K379 lies opposite to a secondary site we previously identified as critical for the oligomerizing activity of **11** and as a secondary binding site. Because these two sites are located on opposing faces of the CRD, we hypothesized that covalent modification at K379 in one CRD and non-covalent binding at the secondary site in a neighboring CRD could promote inter-CRD interactions. To explore this, we analyzed molecular dynamics simulations of the DC-SIGN tetramer, which revealed that such a dual interaction is possible at the CRD-CRD interface (Figure 5f). In this complex, the 3,5-bis(trifluoromethyl)phenyl moiety is directly interacting with residues of the secondary site, consequently explaining the lack of oligomerization upon modification of the M270F mutant (Figure 5g).

Overall, considering the modified protein to strongly increase binding to the mannose chip with a hill slope > 1, we concluded the covalent coupling of **11** at CRD A and non-covalent interaction with the secondary site at another CRD B to fix the resulting oligomer in a geometric orientation that would favor multivalent interaction with glycans, therefore increasing avidity (Figure 5h). Most probably, the same mechanism is also present in the ECD, for which we have previously shown that the secondary site is accessible in the flexible relative arrangement of CRDs in the tetramer ^[10b]^.

## Conclusion

We identified and characterized the first covalent activators of the human C-type lectin DC-SIGN through systematic functional screening of an electrophilic fragment library, biophysical validation, and structural analyses. Our results reveal distinct mechanisms by which these activators modulate glycan recognition. These results emphasize the benefits considering a variety of warheads in covalent screening libraries. In addition to their different reactivity that enables effective mapping the tractable sites of the target, different pre-reaction, transition state and post reaction geometries due to their different labeling chemistries might help exploring different functional mechanisms. While the NHS ester-containing **11** induces oligomerization via covalent modification at multiple lysine residues combined with non-covalent interactions at a previously identified secondary site, **33** achieves selective, non-oligomerizing activation through targeted covalent modification of K373 through its squarate warhead directly within the carbohydrate binding site. Although **11** significantly enhances glycan binding, its oligomerization-dependent mechanism complicates structural characterization and mechanistic interpretation, consequently complicating further development of compounds using the same mechanism. Conversely, the well-defined, single-site modification by **33** allows precise structural insight into its activation mode. This is, to the best of our knowledge, the first report of site-specific covalent modulation of a C-type lectin, and highlights lysine-targeted covalent modification as a novel and effective strategy for enhancing DC-SIGN-ligand interactions.

## Acknowledgements

C. R., J.L. and G.M.K thank the funding from the European Union’s Horizon 2020 research and innovation programme under the Marie Skłodowska Curie grant agreement no. 956314 ALLODD. M.B. thanks the funding from National Research Fund, Luxembourg, AFR PhD Grant 17929849. This study was supported by the Development Laboratory (PharmaLab) project (RRF-2.3.1-21-2022-00015). K.G. was supported by the Doctoral Excellence Fellowship Programme (DCEP-25-1-BME-53) funded by the National Research Development and Innovation Fund of the Ministry of Culture and Innovation and the Budapest University of Technology and Economics. I.A.B. thanks the European Commission for a Marie-Skłodowska Curie Fellowship (No. 895202). P. Á.-B. is supported by the János Bolyai Research Scholarship of the Hungarian Academy of Sciences. We would like to thank the NMR facility at the Department of Pharmaceutical Sciences, Faculty of Life Sciences, University of Vienna, for providing critical support and access to instrumentation for this work.

## Author contributions

J.L. expressed proteins, conducted functional screening and validation, analyzed NMR data and wrote the manuscript. M.B. expressed proteins, conducted NMR experiments and participated in writing the manuscript. N. Cs. synthesized, collected and characterized the library members and participated in protein MS and MS/MS measurements. K. G. participated in NMR and biochemical measurements. Z. O. performed computational modelling. G. S. performed MS and MS/MS measurements. I.A.B. performed the glycan synthesis, characterized the compounds and participated in GCI experiments. P. Á.-B. supervised the research work, curated data and participated in writing the manuscript. G. M. K. conceived and supervised the research, acquired funding and participated in writing the manuscript. C.R. designed and directed the research, acquired the funding and supported writing of the manuscript.

