## Supplementary Information for "Covalent activation of the C-type lectin DC-SIGN"

### Experimental procedures

#### Protein expression and purification

##### General remarks

Unless stated otherwise, all chemicals, growth media and enzymes used for protein expression and purification were purchased from Sigma Aldrich or Carl Roth. Codon-optimized genes for the bacterial expression of wildtype and mutant DC-SIGN CRD and ECD were purchased from GenScript.

##### DC-SIGN extracellular domain

DC-SIGN ECD wildtype, M270F, K373R, K295R and K340R were produced as previously described [1]. In brief, the protein was expressed insolubly in BL21 (DE3) *E. coli* transformed with a pET30b vector encoding amino acids 64–404 of DC-SIGN. Bacteria were grown in Luria-Bertani (LB) medium with 35 mg/L kanamycin and expression was induced with IPTG at $OD_{600}$ of 0.9. Following lysis, inclusion bodies were harvested by centrifugation and solubilized. The protein was refolded via rapid dilution and dialyzed against 25 mM Tris-HCl, 150 mM NaCl, 25 mM $CaCl_2$, pH 7.8, for subsequent purification via mannan agarose affinity chromatography. Purified protein was dialyzed against 25 mM HEPES, 150 mM NaCl, 10 mM $CaCl_2$, pH 7.4, concentrated using a centrifugal spin filter, snap frozen and then stored at -80°C.

##### DC-SIGN carbohydrate recognition domain

Unlabeled and $^{15}N$-labelled DC-SIGN CRD wildtype and M270F were produced as previously described [1]. In brief, the protein was expressed insolubly in BL21 (DE3) *E. coli* transformed with a pET28a vector encoding amino acids 253–404 of DC-SIGN and a *N*-terminal His-tag. Bacteria grown in M9 minimal medium supplemented with 35 mg/L kanamycin and 0.5 g/L $^{15}NH_4Cl$ were induced at $OD_{600}$ of 0.9 using IPTG, lysed and inclusion bodies were harvested by centrifugation. Following solubilization, the protein was refolded overnight *via* rapid dilution. Next, the protein was dialyzed against 50 mM Tris-HCl, 150 mM NaCl, 10 mM $CaCl_2$ (pH 7.8) and purified *via* $Ni^{2+}$ NTA affinity chromatography. Purified protein was pooled and dialyzed against 20 mM MES, 40 mM NaCl, 10 mM $CaCl_2$ (pH 6.0), concentrated using a centrifugal spin filter, snap frozen and then stored at −80 °C.

### Plate-based horseradish peroxidase assay

#### General remarks

Unless otherwise stated, all reagents and buffers were obtained from Sigma Aldrich or Carl Roth. Compounds **1-34** were dissolved in $d_6$-DMSO and stored at −80 °C. GraphPad Prism was used for all data processing and analysis.

#### Screening

100 µg/mL DC-SIGN ECD were immobilized in immobilization buffer (25 mM HEPES, 150 mM NaCl, 25 mM $CaCl_2$, pH 7.4) overnight at 4 °C on transparent Nunc Maxisorp 96-well plates (Thermo Fisher). Protein solution was removed for subsequent testing of covalent compounds. For single point screening of the covalent library, compounds were diluted to 0.5 mM in compound buffer (25 mM HEPES, 150 mM NaCl, 25 mM $CaCl_2$, 5% DMSO) and added to the plate-immobilized DC-SIGN ECD. Control wells for the DMSO reference, and the mannan and EDTA controls included only 25 mM HEPES, 150 mM NaCl, 25 mM $CaCl_2$, 5% DMSO. Plates were incubated overnight at 4 °C or at RT. Plates were washed three times with immobilization buffer and then blocked for 2 hours at 4 °C with blocking buffer (25 mM HEPES, 150 mM NaCl, 25 mM $CaCl_2$, 2% BSA, 0.01% Tween-20). Plates were washed three times and then incubated for 2 hours at RT with 0.5 µg/mL HRP in immobilization buffer. Inhibitory controls additionally contained 10 µg/mL mannan or 25 mM EDTA. Following incubation, plates were washed three times with immobilization buffer and bound HRP was detected using a TMB substrate kit (Thermo Fisher) and 0.18 M $H_2SO_4$ according to manufacturer's instructions. Absorption was measured at 450 nm.

#### Dose-response experiments

Immobilization, blocking and HRP detection steps were done as described above. Compounds were serially diluted on the plates with immobilized DC-SIGN ECD in compound buffer starting from 1 mM and incubated overnight at RT. For time-dependency experiments the reaction was stopped at defined timepoints by washing three times with immobilization buffer supplemented with 25 mM Tris (pH 7.4). Serial dilution of mannose was used as control instead of EDTA and mannan.

### Grating-coupled interferometry

#### General remarks

Unless stated otherwise, all reagents and buffers were obtained from Sigma Aldrich or Carl Roth. The assay was performed in a GCI-based biosensor (WAVEdelta, Creoptix AG/Malvern Panalytical) and a PCP WAVEchip sensor chip (Creoptix AG/Malvern Panalytical). GCI data referencing, analysis and data fitting were performed in Scrubber 2.0 (BioLogic Software) and Graphpad Prism.

**Sensor preparation**

The chip surface was conditioned in 10 mM sodium tetraborate, 1 M NaCl, pH 8.5. Mannose-PEG3- $NH_2$ (Sussex Research Laboratories) and Man(α1,2)Man-PEG3-$NH_2$ were dissolved in ultrapure water to 100 mM and then diluted to 5 mM in 25 mM HEPES, 150 mM NaCl, pH 7.4. The sensor chip was functionalized at room temperature by EDC/NHS activation and chip channels 2 and 3 were coupled to mannose-PEG3-$NH_2$ and Man(α1,2)Man-PEG3- $NH_2$ by injection for 1200 s, respectively. After immobilization, the chip surface was deactivated with 1 M ethanolamine for 420 s. Channel 1 served as a reference channel and was only passivated with 1 M ethanolamine without prior ligand injection.

**Dose-response experiments and affinity measurements**

For dose-response measurements with DC-SIGN ECD, compounds were serial diluted to in 25 mM HEPES, 150 mM NaCl, 10 mM $CaCl_2$, 5% DMSO containing 25 µM protein. Samples were incubated with serial dilutions of the compounds starting from 1 mM overnight at RT. Tris (pH 7.4) was added to a final concentration of 5 mM to quench the reaction. The sample was further diluted in running buffer (25 mM HEPES, 150 mM NaCl, 10 mM $CaCl_2$, 5% DMSO, 0.005% Tween-20, 5 mM Tris, pH 7.4) to minimize buffer mismatch and to reach a final protein concentration of 10 µM. Samples were centrifuged to remove precipitates prior to injection. Samples containing only DMSO and dilution series of mannose served as controls. Binding of modified DC-SIGN ECD was tested in multi-cycle runs using a baseline injection of 45 s followed by 60 s association and 60 s dissociation cycle at a flow rate of 5 µL/min. Regeneration buffer containing 25 mM HEPES, 150 mM NaCl, 50 mM EDTA (pH 7.4) was injected for 120 s after each cycle, followed by 60 s injection of running buffer at a flow rate of 50 µL/min. Blank injections of running buffer after every 5th cycle were used for double referencing.

For dose-response measurements with DC-SIGN CRD, protein samples were dialyzed against 25 mM HEPES, 150 mM NaCl, 10 mM $CaCl_2$, pH 7.4 prior to incubation with serial dilutions of the compounds starting from 1 mM overnight at RT. The procedure was as described for the ECD, but effective protein concentration used for injection was 20 µM.

For determination of the affinity of ECD and CRD (unmodified or modified with 1 mM **11** or **33** overnight at RT) for the glycosylated chip, DC-SIGN ECD and CRD were titrated over the immobilized ligand in running buffer (25 mM HEPES, 150 mM NaCl, 10 mM $CaCl_2$, 0.005% Tween-20, pH 7.4). ECD samples were serially diluted starting from 25 µM, and CRD samples starting from 100 µM. Binding measurements were performed in multi-cycle mode with a 45 s baseline, 60 s association, and 60 s dissociation phase at a flow rate of 5 µL/min. Regeneration was carried out after each cycle by injecting regeneration buffer (25 mM HEPES, 150 mM NaCl, 50 mM EDTA, pH 7.4) for 120 s, followed by 60 s of running buffer at 50 µL/min. Blank buffer injections were included every fifth cycle for double referencing.

#### $^{1}$H-$^{15}$N HSQC NMR

##### General remarks

Unless otherwise stated, all reagents and buffers were obtained from Sigma Aldrich or Carl Roth. NMR tubes were purchased from Bruker. $^{15}$N HSQC NMR experiments were conducted on an Ultra Shield 500 MHz spectrometer (Bruker) equipped with a TCI H/F-C-N Prodigy probe. All $^{1}$H-$^{15}$N HSQC spectra were collected at a sample temperature of 298 K with 256 increments in nitrogen, 24 scans per increment and 2048 points in the direct dimension. The relaxation delay d1 was set to 1.0 s. The W5 Watergate pulse sequence was used for solvent suppression [2]. Spectra were processed in TopSpin 4.2.0. Data analysis was performed using CCPN Analysis 3.2.0 [3].

##### **Titration and binding site identification**.

For binding site identification via $^{1}$H-$^{15}$N HSQC NMR, protein samples were dialysed against 25 mM HEPES, 150 mM NaCl, 10 mM $CaCl_2$, pH 7.4 prior to incubation with compounds. Samples were prepared in 1 mL Eppendorf tubes using 5% $d_6$-DMSO, 10% $D_2O$, and 200 µM uniformly $^{15}$N-labeled DC-SIGN CRD in 25 mM HEPES, 150 mM NaCl, and 10 mM $CaCl_2$ at pH 7.4. Samples were adjusted to a final volume of 160 µL using Milli-Q water. All samples were incubated overnight at RT after adding the respective ligands in ligand-to-protein molar ratios of 2.5:1, 5:1, and 10:1. Post-incubation, the samples were subjected to centrifugal filtration using 0.5 mL 3 kDa MWCO Amicon Ultra centrifugal filters (Thermo Fisher Scientific). Centrifugation runs were performed at 5000 × g for 15 min at 4 °C, after every run, protein buffer (20 mM MES, 40 mM NaCl, 10 mM $CaCl_2$, pH 7.4) was replenished to the original sample volume. The buffer exchange was repeated four times to ensure complete removal of unreacted ligand. Final samples were transferred into 3 mm NMR tubes (Bruker) and analyzed using standard acquisition parameters as described in the general remarks section.

To analyze chemical shift perturbations (CSPs), a previously published resonance assignment of DC-SIGN CRD was transferred to the nearest neighbor in a reference spectrum recorded without compound [4]. Unassigned, overlapping or disappearing peaks were not assigned.

CSPs were calculated as previously described according to **Equation 1** and then mapped to the X-ray crystallographic structure of DC-SIGN CRD (PDB ID: 1SL4) [5].

$$\mathrm{CSP} = \sqrt{\frac{\left(\delta(^{1}H)\right)^2 + \left(\alpha \cdot \delta(^{15}N)\right)^2}{2}} \tag{1}$$

The empirical weighing factor α was set to 0.15 for all calculations.

#### Dynamic light scattering

For protein size assessment via DLS measurements, samples were immediately recovered from the NMR tubes, and the final volume was adjusted to 500 µL using Milli-Q water. The samples were transferred to disposable UV-transparent cuvettes (Sarstedt). The measurements were performed using a Zetasizer Pro DLS (Malvern Panalytical) at 25 °C and 120 sec equilibration time. All datasets were processed using the ZS Xplorer software (Malvern Panalytical).

#### Mass spectrometry

For binding site identification via mass spectrometry, the protein samples were diluted to a final concentration of 1 mg/mL in 25 mM HEPES, 150 mM NaCl, 10 mM $CaCl_2$ (pH 7.4) prior to incubation with compounds for 16 h at room temperature. The protein:compound ratio was 1:20, and the final DMSO content was 2%.

##### LC-MS analysis

Mass spectrometric experiments were performed on a high-resolution hybrid quadrupole-time-of-flight mass spectrometer (Waters Select Series Cyclic IMS, Waters Corp., Wilmslow, U.K.). The mass spectrometer operated in positive V mode. Leucine enkephalin was used as Lock

Mass standard. Chromatographic separations were performed on a Waters Acquity I-Class UPLC system, coupled directly to the mass spectrometer.

**Peptide mapping**

Modification sites were determined by RPLC-MS/MS peptide mapping after proteolysis using various enzymes. Briefly, proteins were enzymatically digested after buffer exchange using Amicon Ultra-0.5 mL Centrifugal Filter units (10 kDa, Merck Millipore). Protein samples were reduced by dithiothreitol at 37 °C for 30 min. Chymotryptic cleavage was performed in 50 mM Tris-HCl solution (pH 8.2) with sequencing-grade chymotrypsin (Promega Corporation, Madison, USA) using 1:20 enzyme:protein ratio at 37°C for 2 hours. Digestion was stopped by adding formic acid in a final concentration of 0.2% (V/V). ProAlanase (Promega Corporation, Madison, USA) digestion was performed in 50 mM HCl solution using 1:50 enzyme:protein ratio at 37 °C.

Gradient elution was performed on a Waters Acquity Peptide BEH C18 UPLC column (2.1x150 mm, 1.7 µm) under the following parameters: mobile phase “A”: 0.1% formic acid in water, mobile phase “B”: 0.1% formic acid in acetonitrile; flow rate: 300 µL/min; column temperature: 60 °C; gradient: 2 min: 2% B, 20 min: 55% B, 20.5 min: 90% B. $MS^E$ experiments were performed using collision voltage ramping under the following parameters: *m/z* 50-2000, scan time: 0.3 sec, single Lock Mass: leucine enkephalin; low energy: 6 V, high energy: ramping 19-45 V. BiopharmaLynx 1.3.5 software (Waters Corp., Wilmslow, U.K.) was used to for data analysis.

#### Molecular docking simulation

**Induced fit docking**

The X-ray structure of the carbohydrate binding site of DC-SIGN (PDB ID: 1SL5) was used to perform the docking simulations for the monomeric structure. MD time frames from reference [6] were used for modelling the tetrameric structure. Proteins were prepared with the Protein Preparation Wizard (Schrödinger Release 2024-3) using default methods. Ligands were prepared with Ligprep of the Schrödinger Suite. Docking was performed using the Induced Fit Docking protocol using the OPLS 2005 force field generating 20 possible binding conformations [7]. Redocking was carried out into structures within 125 kJ $mol^{-1}$ of the best structure. and within the top 20 structures overall using the single precision method.

**Covalent Docking**

Protein structures were selected from the 20 possible binding conformations generated by Induced Fit docking based on visual inspection. In case of the monomer structure covalent docking simulations were performed using K373 as reactive residue. The enclosing box was defined as the center of the most relevant CSPs from our HSQC NMR experiments with 20 Å box size. A custom reaction type was used with pose prediction docking mode. 5 Å was set as minimization radius around the reactive residue. 20 possible output poses were selected per ligand reaction site. In case of the tetrameric structure covalent docking simulations were performed using K379 as reactive residue. The enclosing box was defined as the center of the most relevant CSPs from our HSQC NMR experiments with 30 Å box size. A custom reaction type was used with pose prediction docking mode. 5Å was set as minimization radius around the reactive residue. 20 possible output poses were selected per ligand reaction site.

##### Calculation of $pK_a$ and solvent accessible surface area (SASA)

$pK_a$ values were computed with the online version of PropKa using the tetrameric structure of DC-SIGN CRD [8]. The monomer $pK_a$ values were calculated as the average of the four monomers. The $pK_a$ values of lysines in the monomers are shown in table S3.

Solvent accessible surface areas were calculated using the Getarea webserver using a 1.4 Å radius for the water probe [9].

#### Library synthesis

##### General remarks

All chemicals and solvents with >95% purity were purchased from commercial vendors (Sigma-Aldrich, Fluorochem, Ambeed, and Enamine, respectively) and used without further purification. NMR measurements were performed on a Varian Unity Inova 500 NMR spectrometer or a Varian System 300 spectrometer (Varian, Palo Alto, CA, USA). $^{1}H$ NMR, $^{13}C$ NMR and $^{19}F$ NMR spectra were recorded in DMSO-*d6*, $CD_3CN$ or $CDCl_3$ solution at room temperature with the deuterium signal of the solvent as the lock. Chemical shifts (δ) and coupling constants (*J*) are given in parts per million (ppm) and Hz, respectively. Splitting patterns are abbreviated as follows: singlet (s), doublet (d), triplet (t), quartet (q), heptet (hept), multiplet (m), broad singlet (bs) and combinations thereof. All $^{13}C$ NMR spectra were proton decoupled. Reactions were monitored with HPLC-MS, GC-MS or Merck silica gel 60 $F^{254}$ TLC plates (Darmstadt, Germany). HPLC-MS measurements were performed using a Shimadzu LC-MS-2020 device (Shimadzu, Kyoto, Japan) equipped with a Reprospher-100 C18 (5 µm; 100 x 3 mm) column and a positive–negative double ion source (DUIS±) with a quadrupole

MS analyzer in a range of *m/z* 50–1000. Samples were eluted with gradient elution using eluent A (0.1% formic acid in water) and eluent B (0.1% formic acid in acetonitrile). The flow rate was set to 1.5 mL/min. The initial condition was 0% B eluent, followed by a linear gradient to 100% B eluent by 2 min, from 2 to 3.75 min 100% B eluent was retained, and from 3.75 to 4.5 min back to the initial condition and retained to 5 min. The column temperature was kept at 30 °C and the injection volume was 1-10 µL. GC-MS was performed on a Shimadzu GC-2010 instrument equipped with a GCMS-QP2010 Ultra detector, using EI ionization. The analytical method used a Zebron ZB-5MSi column (30 m, ID: 0.25 mm, df: 0.25 µm), helium as a carrier gas, and an oven temperature profile as follows: 0 min.: 80°C; 11 min.: 250°C; 16 min.: 250°C. Further parameters: injector temperature: 250°C; gas flow rate: 27.7 mL/min.; split ratio: 1/20, detector temperature: 280°C. HRMS measurements were performed on a Triple TOF 5600+ QTOF mass spectrometer (Sciex, Framingham, MA, USA) equipped with DuoSpray ion source. The resolution of the instrument was above 30 000 over the entire mass range. Samples were introduced into an acetonitrile flow (0.2 mL/min) in flow injection mode. Analyst TF 1.8.1 software (Sciex, Framingham, MA, USA) was used for controlling the instrument and for data processing. Flash column chromatography was performed using a CombiFlash Rf 200 instrument (Teledyne ISCO, Lincoln, NE). Normal-phase flash column chromatography was performed on Merck Silica Gel 60 (particle size 0.040 − 0.063 mm; Merck, Darmstadt, Germany). In the case of reversed-phase chromatography, RediSep Rf reversed-phase C18 columns (4.3 g, 26 g, 43 g, and 86 g) were used. Preparative HPLC was performed with a Combi Flash EZ PREP (Teledyne ISCO, Lincoln, NE) instrument equipped with a Gemini 5 µm NX-C18 110 LC Column (150 x 21.2 mm).

**Synthesis of 3,5-bis(trifluoromethyl)benzene-1-sulfonyl fluoride (3)**

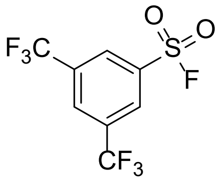

3,5-Bis(trifluoromethyl)benzenesulfonyl chloride (120 mg, 0.38 mmol, 1 eq.) was added to a 1:1 (V/V) mixture of acetonitrile/saturated potassium bifluoride ($HF_2K$) (10 ml) solution. The resulting mixture was stirred for 90 min then extracted twice with diethyl ether (10-10 mL). The combined organic layers were washed with brine (5 mL), dried over $MgSO_4$, filtered and concentrated in vacuo at room temperature. The crude product was eluted on a silica pad in 100% hexane, resulting in a colorless oil (67 mg, 59% yield).

$^{1}H$ NMR (300 MHz, $CDCl_3$) δ 8.47 (s, 2H), 8.29 (s, 1H).

$^{13}C$ NMR (75 MHz, $CDCl_3$) δ 135.42 (d, $J$ = 28.8 Hz), 133.71 (q, $J$ = 35.2 Hz), 129.07 – 128.76 (m), 128.63 – 128.40 (m), 121.66 (q, $J$ = 273.5 Hz).

$^{19}F$ NMR (282 MHz, $CDCl_3$) δ -63.14 (s), -193.40 (s).

NMR spectra are in agreement with the literature [10].

GC-MS $m/z$: calculated for $C_8H_3F_7O_2S$: 296; measured: 296.

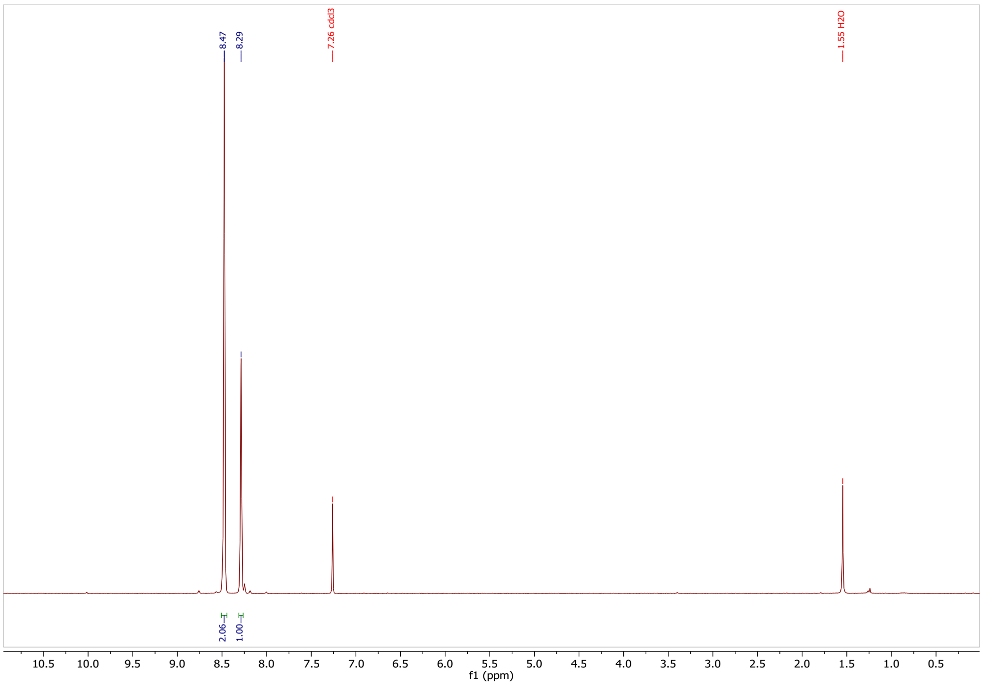

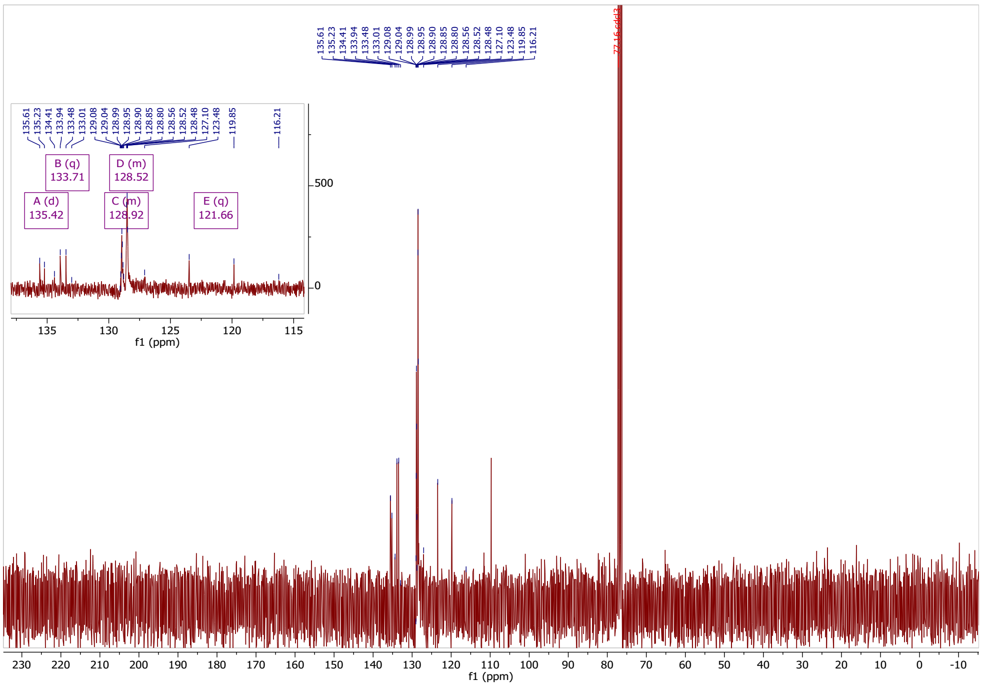

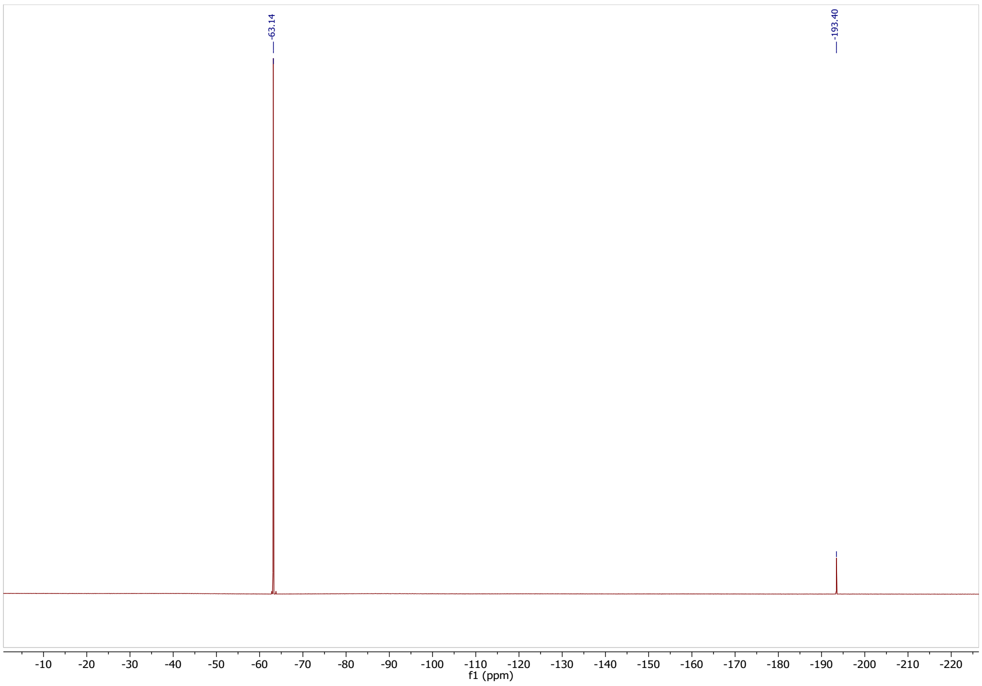

**Synthesis of 3,5-bis(trifluoromethyl)phenyl sulfurofluoridate (4)**

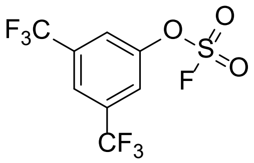

3,5-Bis(trifluoromethyl)phenol (46 mg, 0.20 mmol, 1 eq.) was dissolved in DCM (3 mL) and TEA (45 µL, 0.32 mmol, 1.6 eq.) was added. To this mixture 1-(fluorosulfonyl)-2,3-dimethyl-1*H*-imidazol-3-iumtrifluoromethane sulfonate (85 mg, 0.26 mmol, 1.3 eq.) was added and stirred 1 h. The reaction was monitored by TLC (hexane/ethyl acetate, 10:1). Upon completion, the reaction mixture was filtered through a silica pad, washed with DCM (5 x 2 mL) and the solvent was evaporated at room temperature resulting in a colorless liquid (46 mg, 74 % yield).

$^1$H NMR (300 MHz, $CD_3CN$) δ 8.17 (s, 1H), 8.11 (s, 2H).
$^{13}$C NMR (126 MHz, $CDCl_3$) δ 149.99, 134.61 (q, J = 35.0 Hz), 122.98 (hept, J = 3.7 Hz), 122.24 (q, J = 273.7 Hz), 122.20-122.13 (m).
$^{19}$F NMR (282 MHz, $CD_3CN$) δ -63.58, -221.63.
GC-MS *m/z*: calculated for $C_8H_3F_7O_3S$: 311; measured: 311.

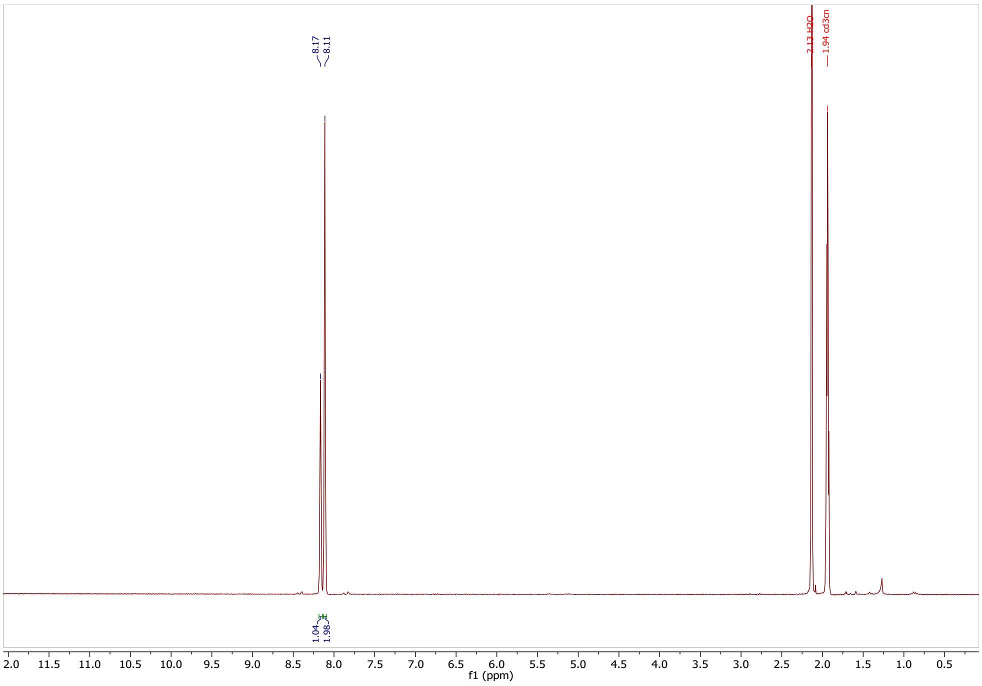

8.17
8.11
2.13 H2O
1.94 cd3cn
1.04
1.98
f1 (ppm)

1

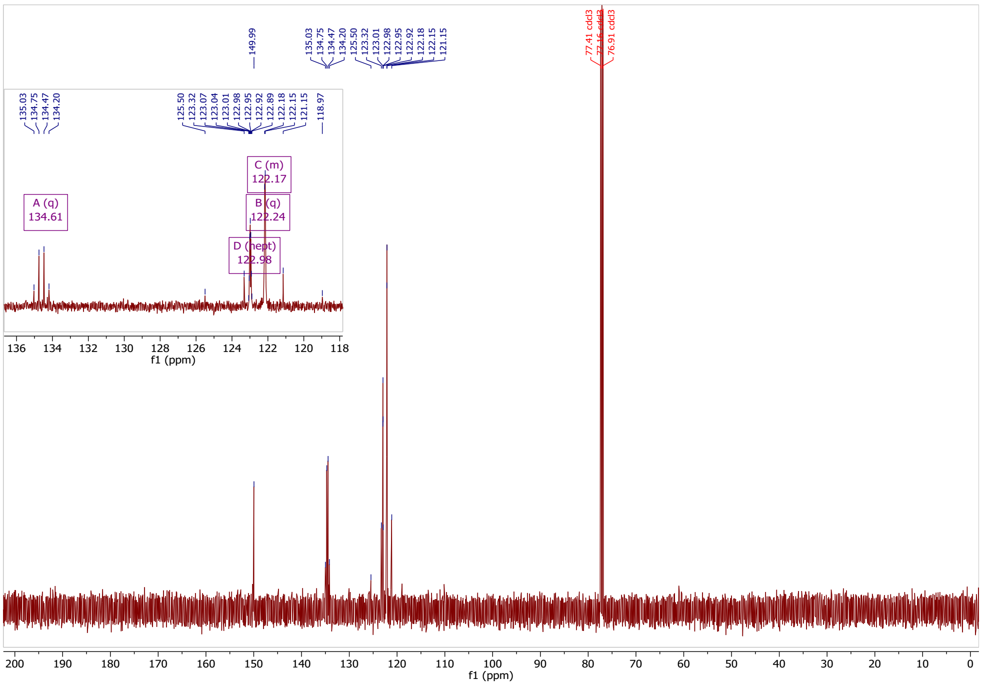

149.99
135.03
134.75
134.47
134.20
125.50
123.32
123.01
122.98
122.95
122.92
122.18
122.15
121.15
77.41 cdcl3
76.91 cdcl3
A (q)
134.61
B (q)
122.24
C (m)
122.17
D (hept)
122.98
118.97
f1 (ppm)

2

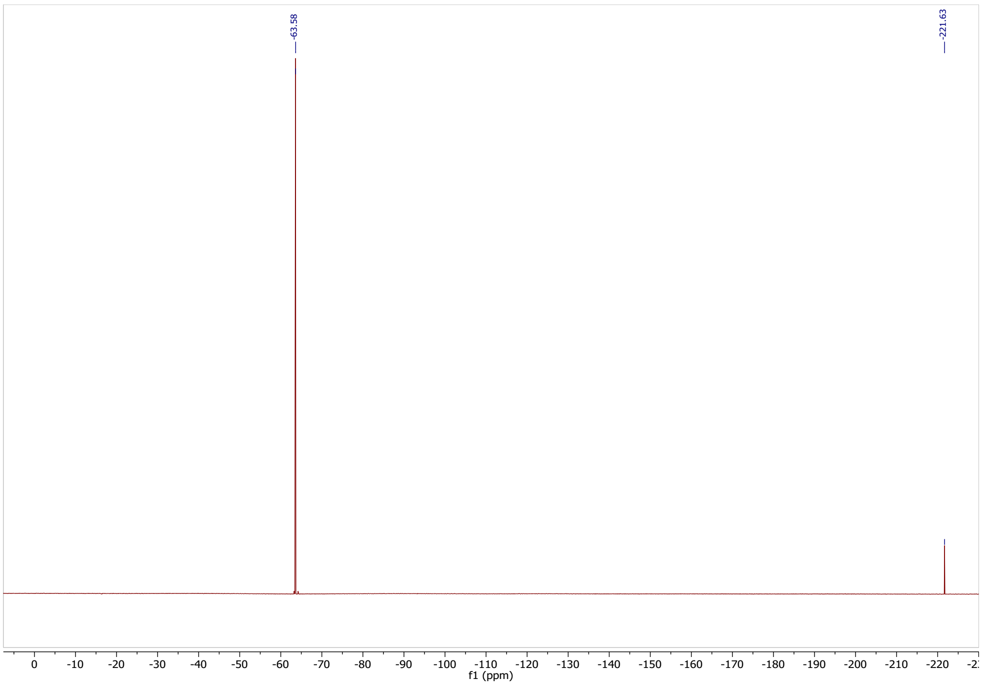

**Synthesis of 1-((3,5-bis(trifluoromethyl)phenyl)sulfonyl)-3-(4-methoxyphenyl)-1*H*-1,2,4-triazole (5)**

***N-((dimethylamino)methylene)-4-methoxybenzamide***

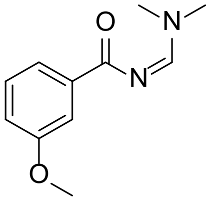

4-methoxybenzamide (1 g, 6.6 mmol, 1 eq.) was dissolved in DMF-DMA (2.55 mL, 19 mmol, 2.9 eq.) and stirred for 30 min at 120 °C. Next, the reaction mixture was allowed to reach room temperature then DEE (10 mL) was added and cooled to 0 °C. The resulting precipitate was filtered and washed with DEE/Hexane 1:1 affording the product, *N*-((dimethylamino)methylene)-4-methoxybenzamide as a white crystal (1.36 g, 74%) which was used in the next step without further purification.

LC-MS *m/z* calculated for $C_{11}H_{14}N_2O_2$: 206; measured: $[M+H]^+$, 207.

***3-(4-methoxyphenyl)-1H-1,2,4-triazole***

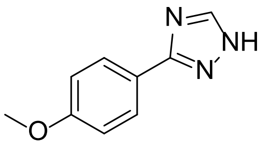

In a two-neck round bottom flask *N*-((dimethylamino)methylene)-4-methoxybenzamide (1.0 g, 4.8 mmol, 1 eq.) was dissolved in acetic acid (3.7 mL) and warmed up to 90 °C. Hydrazine (0.28 mL, 5.3 mmol, 1.1 eq.) was added dropwise to the mixture in 10 minutes and stirred for 90 minutes. After cooling down to room temperature, saturated aq. $NaHCO_3$ solution (15 mL) was added dropwise to the mixture until the solution was neutralized. Next, DEE (20 mL) was added and cooled to 0 °C resulting a white precipitate which was filtered and washed with cold DEE (2 x 2 mL) to afford the product as a white crystal (0.695 g, 82 %). The product was used in the next step without further purification.

$^1$H NMR (300 MHz, DMSO-*d6*) δ 8,26 (s, 1H), 7,95 (d, *J* = 8,8 Hz, 2H), 7,04 (d, *J* = 8,8 Hz, 2H), 3,80 (s, 3H).
LC-MS *m/z* calculated for $C_9H_9N_3O$: 175; measured $[M+H]^+$: 176.

NMR spectra is in agreement with the literature [11].

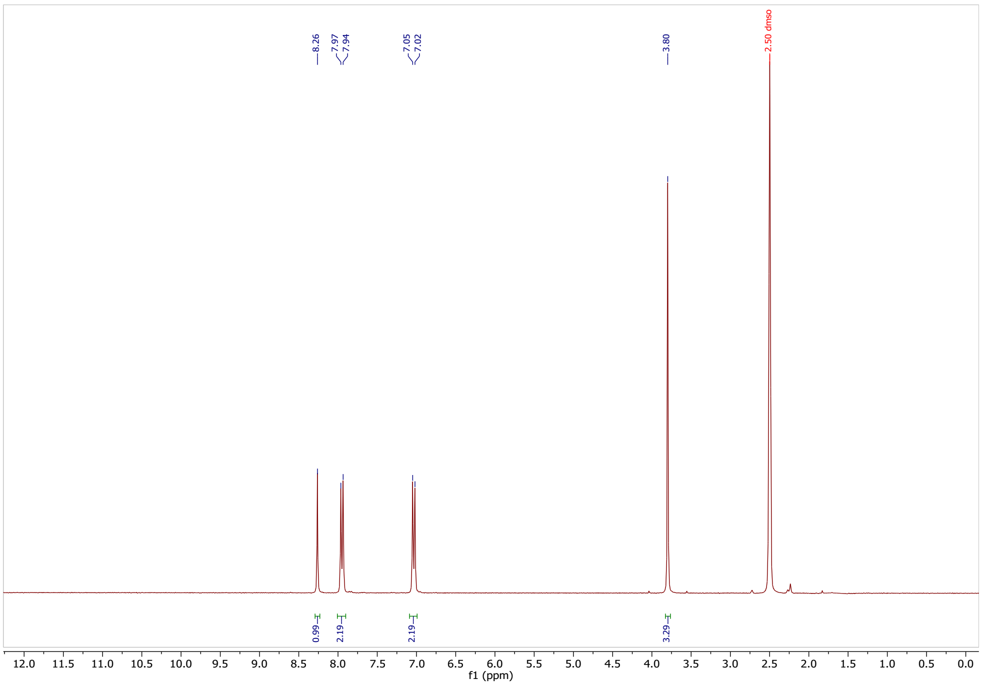

***1-((3,5-bis(trifluoromethyl)phenyl)sulfonyl)-3-(4-methoxyphenyl)-1H-1,2,4-triazole (5)***

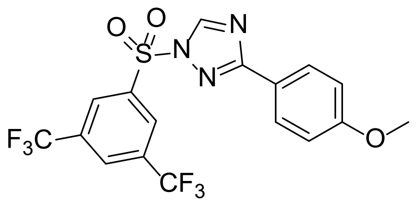

3-(4-Methoxyphenyl)-1*H*-1,2,4-triazole (0.4 g, 2.28 mmol, 1 eq.) was dissolved in MeCN (15 mL) and TEA (0.38 mL, 2.74 mmol, 1.2 eq.) was added. The mixture was cooled down to 0 °C. Next, 3,5-bis(trifluoromethyl)benzenesulfonyl chloride (0.715 g, 2.28 mmol, 1 eq.) in MeCN (10 mL) was added dropwise and stirred at RT for overnight. The solvent was evaporated, then the crude product was dissolved in DCM (25 mL) and extracted with water (10 mL) followed by brine (15 mL), dried over $Na_2SO_4$, filtered and concentrated in vacuo. The crude product was purified by normal-phase flash column chromatography (Hex/EtOAc 0-25%) affording the product as a white powder (0.73 g, 71% yield).

$^1$H NMR (300 MHz, $CDCl_3$) δ 8.75 (s, 1H), 8.59 (s, 2H), 8.20 (s, 1H), 8.03 (d, *J* = 8.5 Hz, 2H), 6.95 (d, *J* = 8.6 Hz, 2H), 3.85 (s, 3H).

$^{13}$C NMR (75 MHz, $CDCl_3$) δ 166.15, 162.01, 145.71, 139.10, 133.86 (q, *J* = 35.0 Hz), 129.35 – 128.66 (m), 128.95, 122.16 (q, *J* = 273.4 Hz), 121.23, 114.34, 55.53.

$^{19}$F NMR (282 MHz, $CDCl_3$) δ -63.01.

HRMS (ESI/Q-TOF) *m/z*: $[M+H]^+$ Calcd. for $C_{17}H_{12}N_3O_3F_6S$, 452.0503, measured: 452.0506

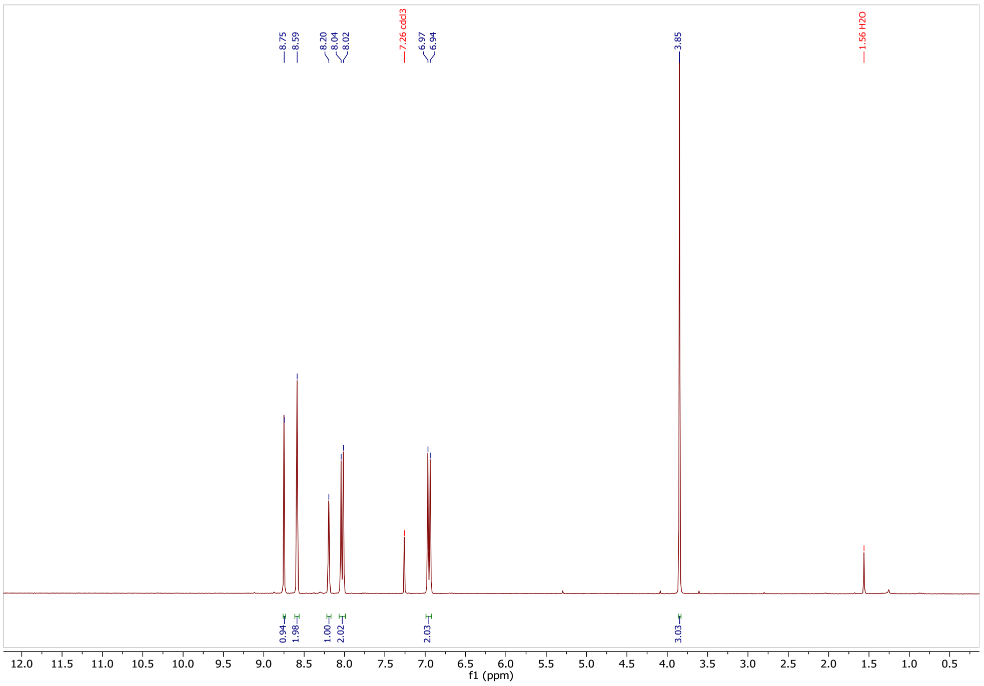

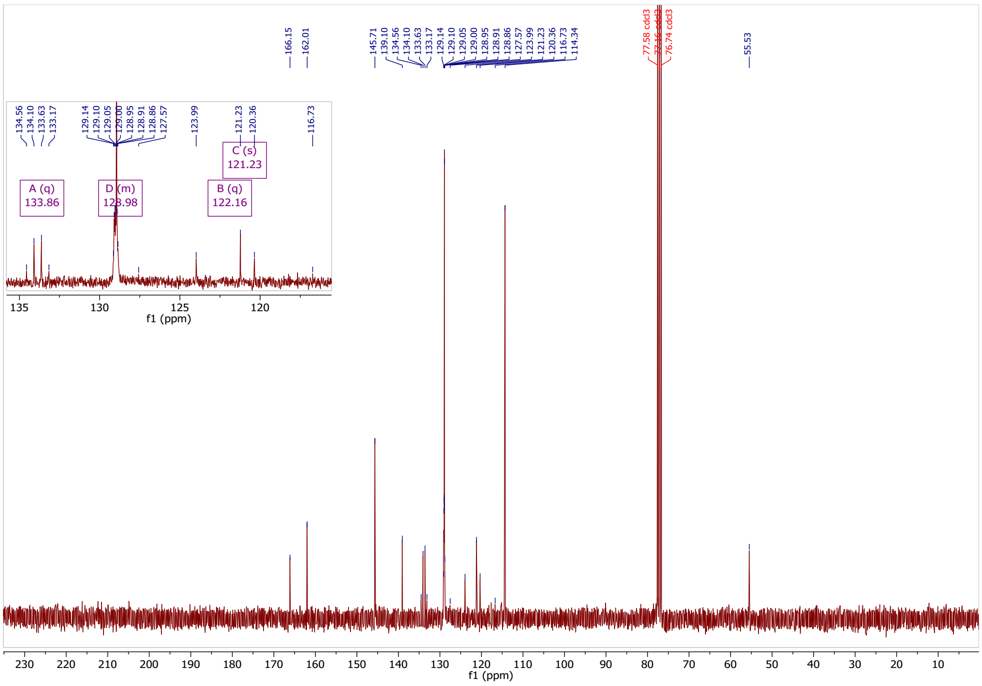

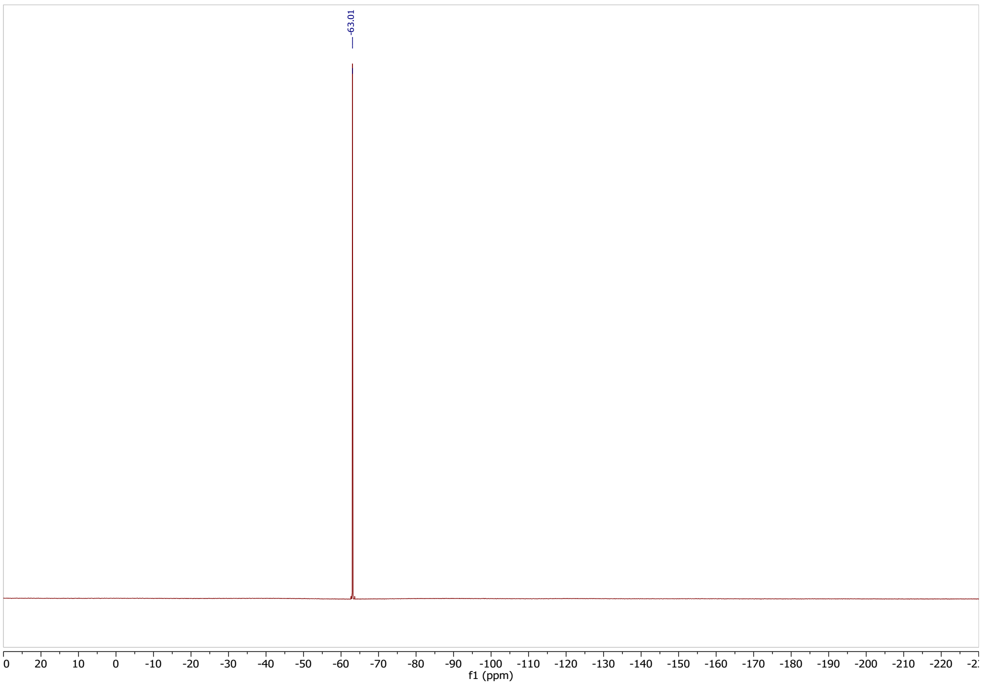

**1*H*-benzo[*d*][1,2,3]triazol-1-yl 3,5-bis(trifluoromethyl)benzenesulfonate (6)**

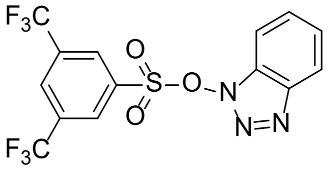

Under argon 3,5-bis(trifluoromethyl)benzenesulfonyl chloride (100 mg, 0.32 mmol, 1 eq.) was added dropwise to a suspension of HOBt (43 mg, 0.32 mmol, 1 eq.), anhydrous $CH_2Cl_2$ (5 mL) and triethylamine (0.045 mL, 0.32 mmol, 1 eq.) at 0 °C. The reaction mixture was stirred at 0 °C for 30 min then at room temperature for 2 h. After dilution with 20 mL of $CH_2Cl_2$, the organic phase was washed with water (10 mL) and brine (10 mL) and dried over $MgSO_4$. The organic phase was concentrated on a rotary evaporator affording a white solid (108 mg, 82 % yield).

$^1$H NMR (300 MHz, $CDCl_3$) δ 8.38 (s, 3H), 8.31 (s, 1H), 8.06 (dd, J = 8.5, 1.0 Hz, 1H), 7.75 – 7.62 (m, 2H), 7.53 – 7.46 (m, 1H).

$^{13}$C NMR (75 MHz, $CDCl_3$) δ 135.74, 134.04 (q, *J* = 35.2 Hz), 130.11 (q, *J* = 4.0 Hz), 129.80 (hept, *J* = 3.5 Hz), 126.30 (q, *J* = 273.1 Hz), 125.96, 120.83, 109.24.

$^{19}$F NMR (282 MHz, $CDCl_3$) δ -63.03.

HRMS (ESI/Q-TOF) *m/z*: $[M+H]^+$ Calcd. for $C_{14}H_8N_3O_3F_6S$ 412.0190; found 412.0185.

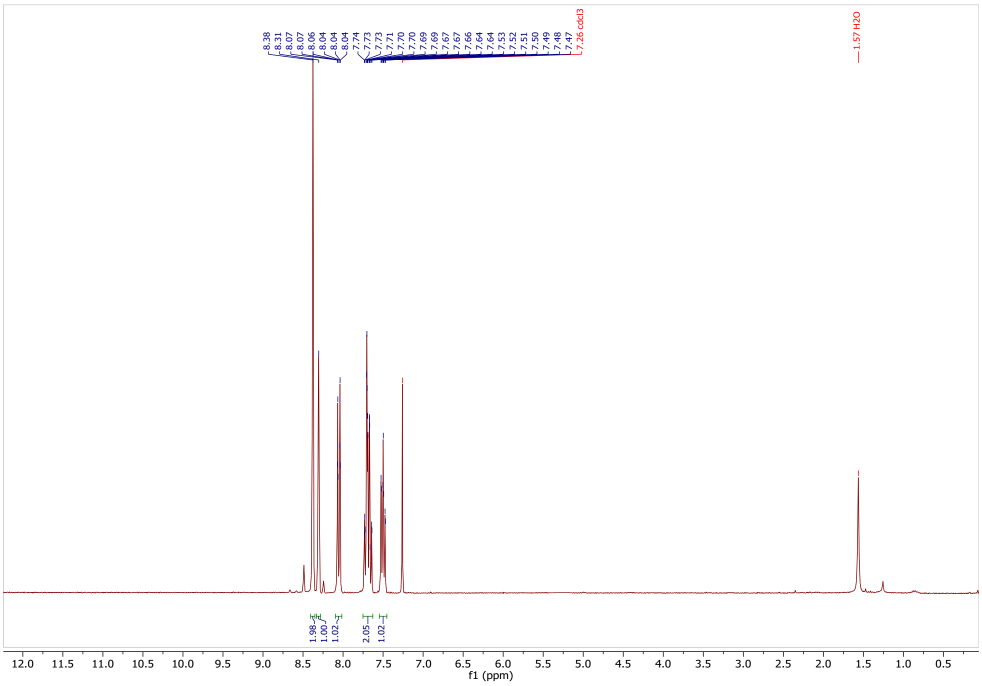

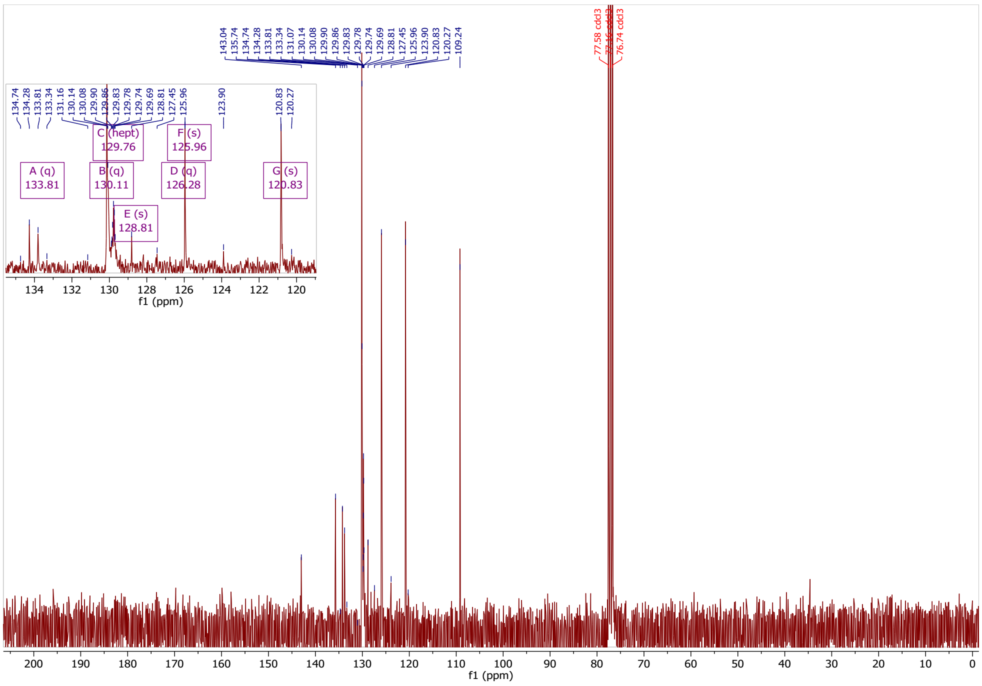

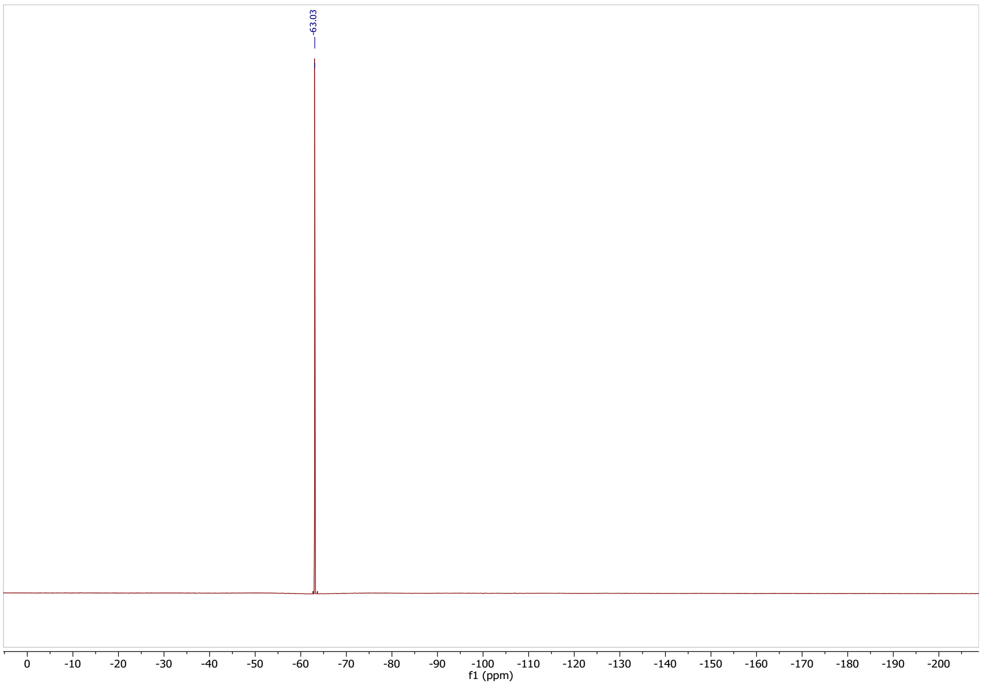

##### Ethyl 2-(((3,5-bis(trifluoromethyl)phenyl)sulfonyl)methyl)acrylate (7)

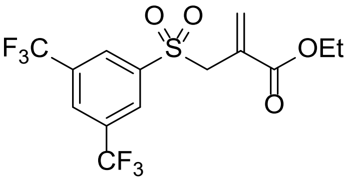

3,5-Bis(trifluoromethyl)benzenesulfonyl chloride (300 mg, 0.96 mmol, 1 eq.), $Na_2SO_3$ (242 mg, 1.92 mmol, 2 eq.) and $NaHCO_3$ (161 mg, 1.92 mmol, 2 eq.) were dissolved in distilled water (10 mL) and stirred for 5 h at 80 °C. After cooling down to room temperature the water was evaporated at reduced pressure. The crude product was dissolved in 10 mL of ethanol and the resulting suspension was filtered. The filtrate was concentrated in vacuo to obtain the sodium aryl sulfinate as a white powder (286 mg, 99% yield) which was used in the next step without further purification [12]. Sodium 3,5-bis(trifluoromethyl)benzenesulfinate (284 mg, 0.95 mmol, 1.5 eq.) and ethyl 2-(bromomethyl)prop-2-enoate (122 mg, 0.63 mmol, 1 eq.) was dissolved in dry methanol (8 mL). The reaction mixture was stirred at 65 °C for 5 h. Upon completion, the methanol was evaporated in vacuo and the crude was dissolved in EtOAc (15 mL). The organic phase was washed with water (10 mL) and brine (10 mL), dried on $Na_2SO_4$. The filtered organic phase was concentrated at reduced pressure and purified by flash chromatography in hexane/ethyl-acetate (0-50%) to give the desired product as a white powder (66 mg, 27%).

$^1$H NMR (300 MHz, $CDCl_3$) δ 8.31 (s, 2H), 8.14 (s, 1H), 6.61 (s, 1H), 6.10 (s, 1H), 4.24 (s, 2H), 4.00 (q, $J$ = 7.1 Hz, 2H), 1.17 (t, $J$ = 7.1 Hz, 3H).
$^{13}$C NMR (75 MHz, $CDCl_3$) δ 164.59, 141.42, 134.67, 133.13 (q, $J$ = 34.7 Hz), 129.53 – 129.10 (m), 128.47, 127.70 – 127.28 (m), 122.47 (q, $J$ = 273.5 Hz), 61.96, 57.85, 14.00.
$^{19}$F NMR (282 MHz, $CDCl_3$) δ -63.01.
HRMS (ESI/Q-TOF) *m/z*: $[M+H]^+$ Calcd. for $C_{14}H_{13}O_4F_6S$, 391.0439; found 391.0431.

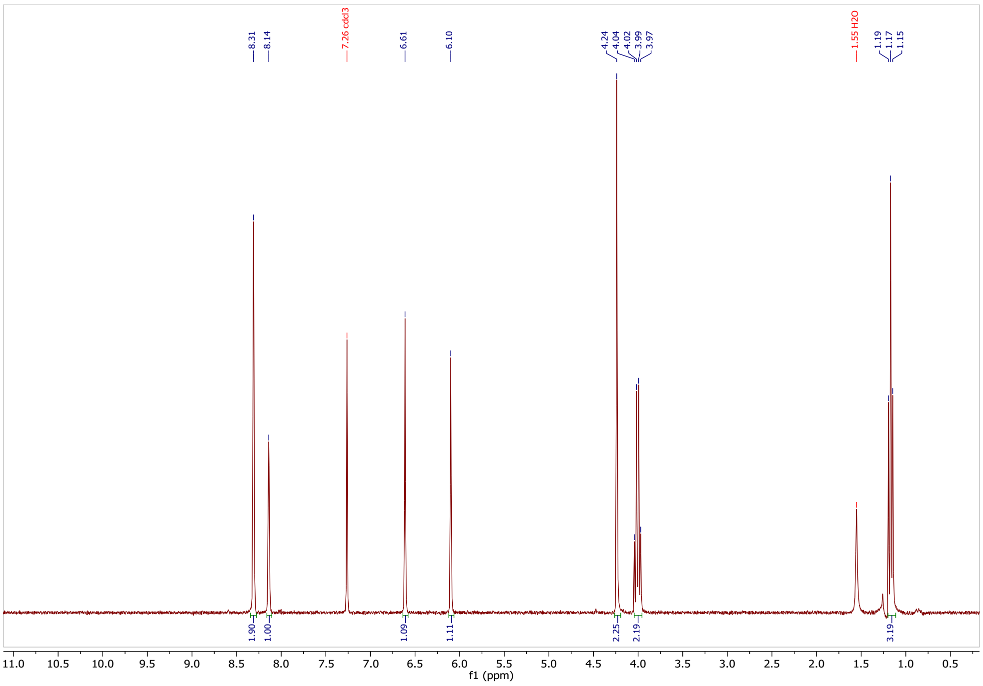

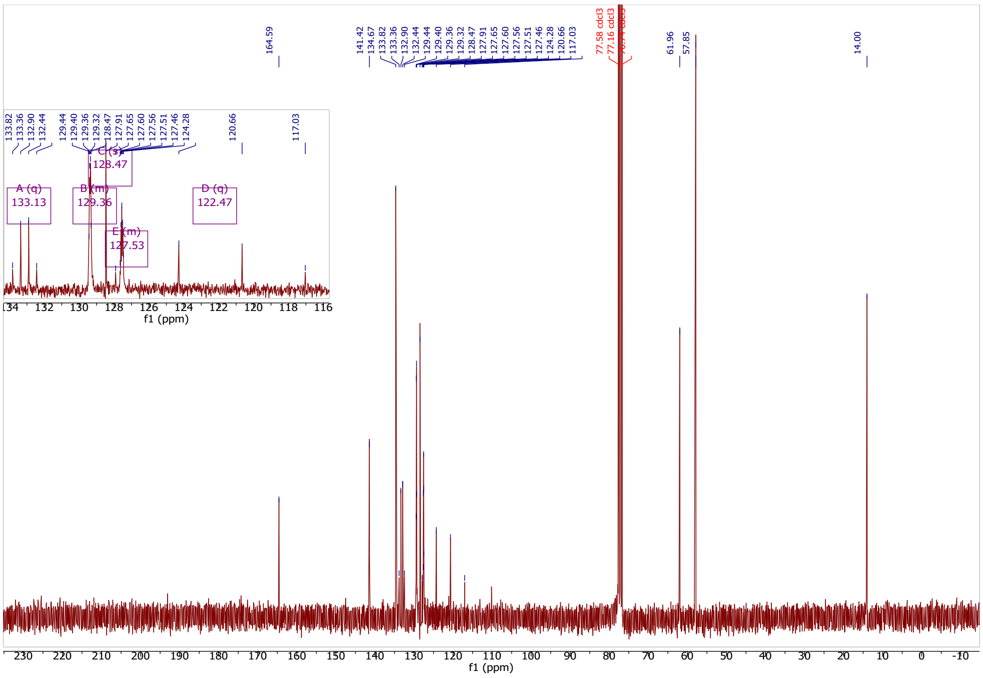

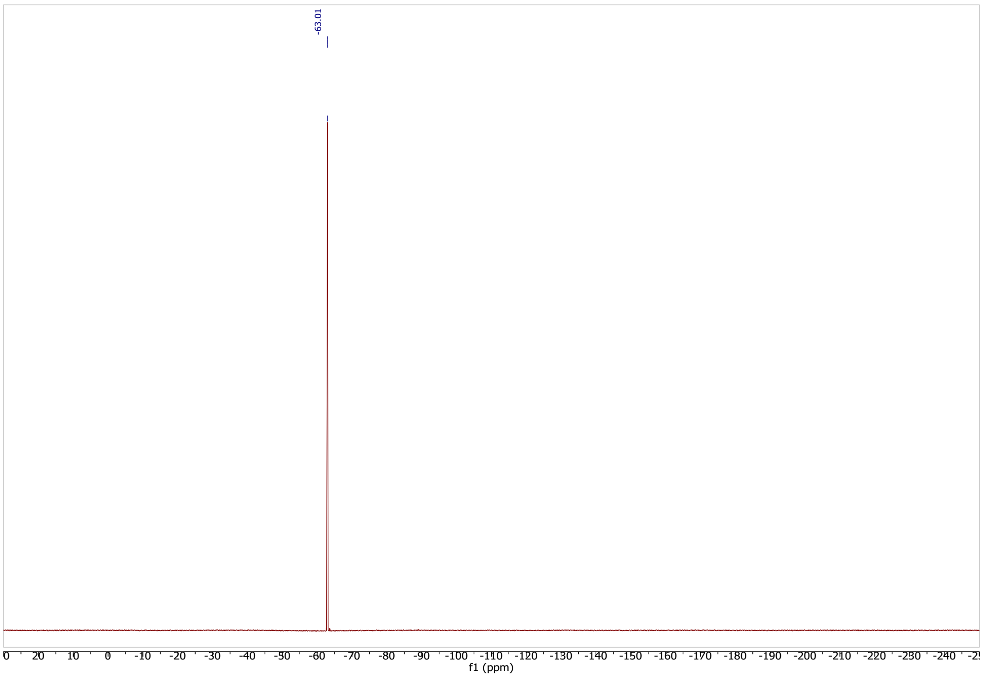

**Synthesis of 2-(3,5-bis(trifluoromethyl)phenyl)-*N*-(cyanomethyl)-*N*-tosylacetamide (9)**

***2-(3,5-bis(trifluoromethyl)phenyl)-N-tosylacetamide***

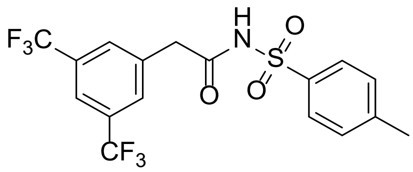

Under argon 2-(3,5-bis(trifluoromethyl)phenyl)acetic acid (600 mg, 2.20 mmol, 1 eq.), 4-methylbenzenesulfonamide (453 mg, 2.65 mmol, 1.2 eq.), 1-ethyl-3-(3-dimethylaminopropyl) carbodiimide hydrochloride (EDC·HCl, 507 mg, 2.65 mmol 1.2 eq.), 4-(dimethylamino)pyridine (DMAP, 135 mg, 1.10 mmol, 0.5 eq.) was dissolved in DCM (15 mL) and stirred for 16 h at room temperature. Next, the reaction was quenched by 1.0 N aq. HCl (25 mL). The mixture was diluted with EtOAc (20 mL) then separated. The aqueous layer was extracted with additional portions of EtOAc (3 x 20 mL). The combined organic layers were washed with brine (50 mL), dried over $MgSO_4$, filtered and concentrated in vacuo. The crude product was purified by normal-phase flash column chromatography (Hex/EtOAc 0-35%) affording the product as a white solid (471 mg, 50%).

$^1$H NMR (300 MHz, $CD_3CN$) δ 9.66 (br.s, 1H), 7.89 (s, 1H), 7.84 (d, $J$ = 8.1 Hz, 2H), 7.74 (s, 2H), 7.37 (d, $J$ = 8.1 Hz, 2H), 3.75 (s, 2H), 2.41 (s, 3H).
$^{13}$C NMR (75 MHz, $CD_3CN$) δ 169.03, 146.29, 137.54, 137.01, 131.86 (q, $J$ = 33.2 Hz), 131.49 – 131.28 (m), 130.51, 129.04, 124.48 (q, $J$ = 271.5 Hz), 122.26 – 121.97 (m), 42.19, 21.62.
LC-MS (EI) $m/z$: calculated for $C_{17}H_{13}F_6NO_3S$: 425; measured: $[M+H]^+$ 426.

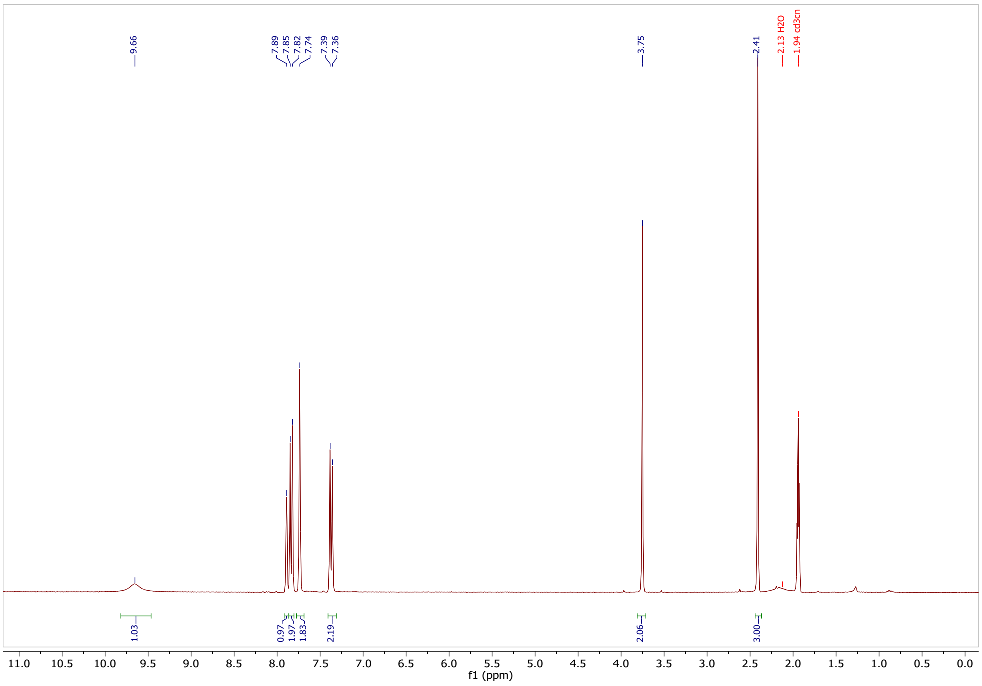

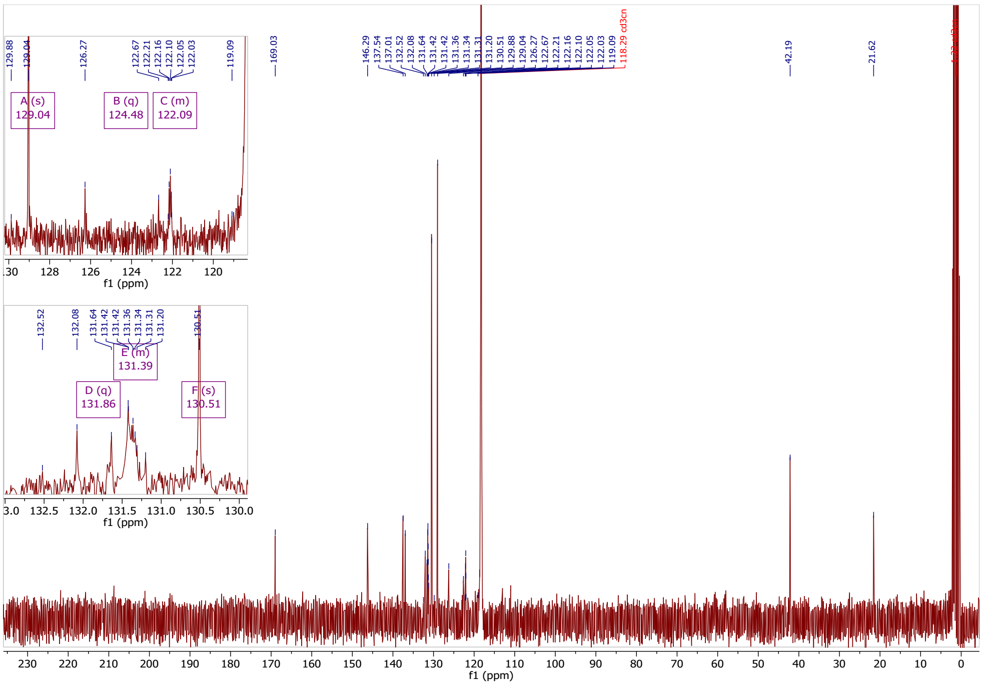

***2-(3,5-bis(trifluoromethyl)phenyl)-N-(cyanomethyl)-N-tosylacetamide***

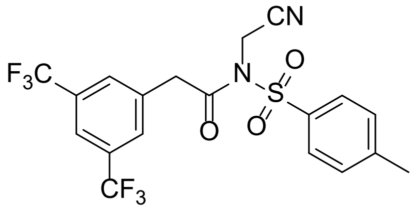

Under argon 2-(3,5-bis(trifluoromethyl)phenyl)-*N*-tosylacetamide (500 mg, 1.18 mmol, 1 eq.) was dissolved in dry DMF (4 mL) and DIPEA (0.61 mL, 3.53 mmol, 3 eq.) was added. Next, 2-iodoacetonitrile (0.83 mL, 11.80 mmol, 10 eq.) was added dropwise via a syringe. The reaction mixture was stirred at room temperature for 1 hour, after which an additional 5 equivalents of 2-iodoacetonitrile (0.42 mL, 5.9 mmol, 5 eq.) were added. Stirring was continued for further at room temperature. After 4 h the reaction was quenched with water (15 mL) and extracted with EtOAc (3 x 20 mL). The combined organic layers were washed with brine (10 mL), dried on $Na_2SO_4$ then filtered and concentrated in vacuo. The reaction mixture was purified by preparative RP-HPLC resulting in the product as a light-brown amorphous solid (84 mg, 15%).

$^1$H NMR (300 MHz, $CD_3CN$) δ 7.94 (s, 1H), 7.92 (s, 2H), 7.74 (s, 2H), 7.49 (d, *J* = 8.1 Hz, 2H), 4.71 (s, 2H), 4.29 (s, 2H), 2.46 (s, 3H).

$^{13}C$ NMR (126 MHz, $CD_3CN$) δ 170.73, 147.72, 137.60, 135.82, 131.79 (q, *J* = 33.3 Hz), 131.74 (q, *J* = 4.2 Hz), 131.38, 129.01, 124.48 (q, *J* = 272.0 Hz), 122.21 – 122.03 (m), 116.36, 42.41, 34.93, 21.70.

$^{19}F$ NMR (282 MHz, $CD_3CN$) δ -63.41.

HRMS (ESI/Q-TOF) *m/z*: $[M+H]^+$ Calcd. for $C_{19}H_{15}N_2O_3F_6S$, 465.0707, measured: 465.0700

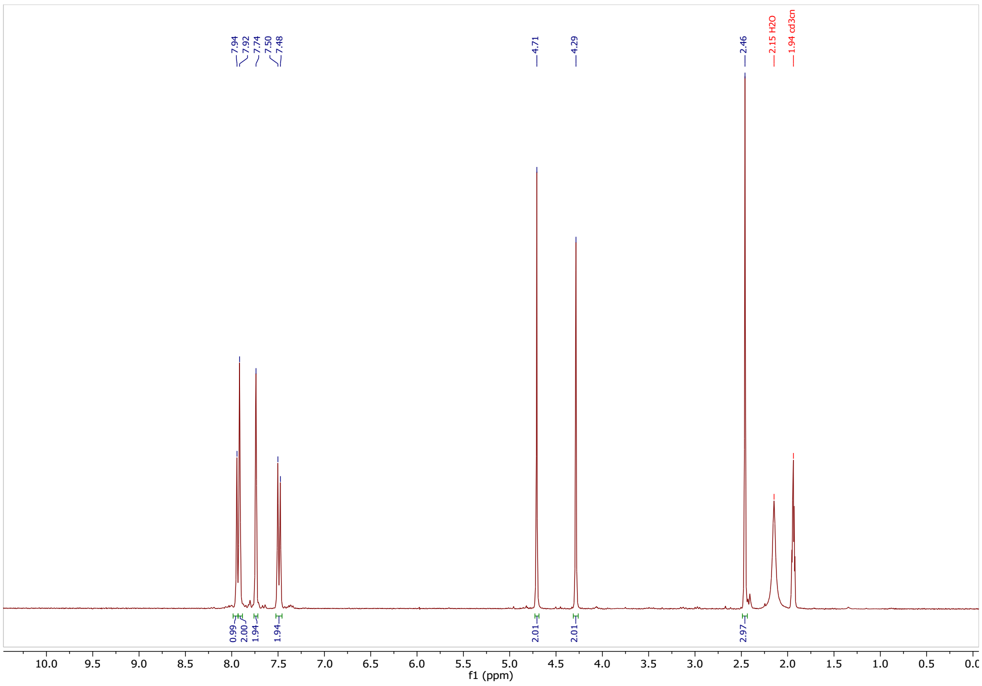

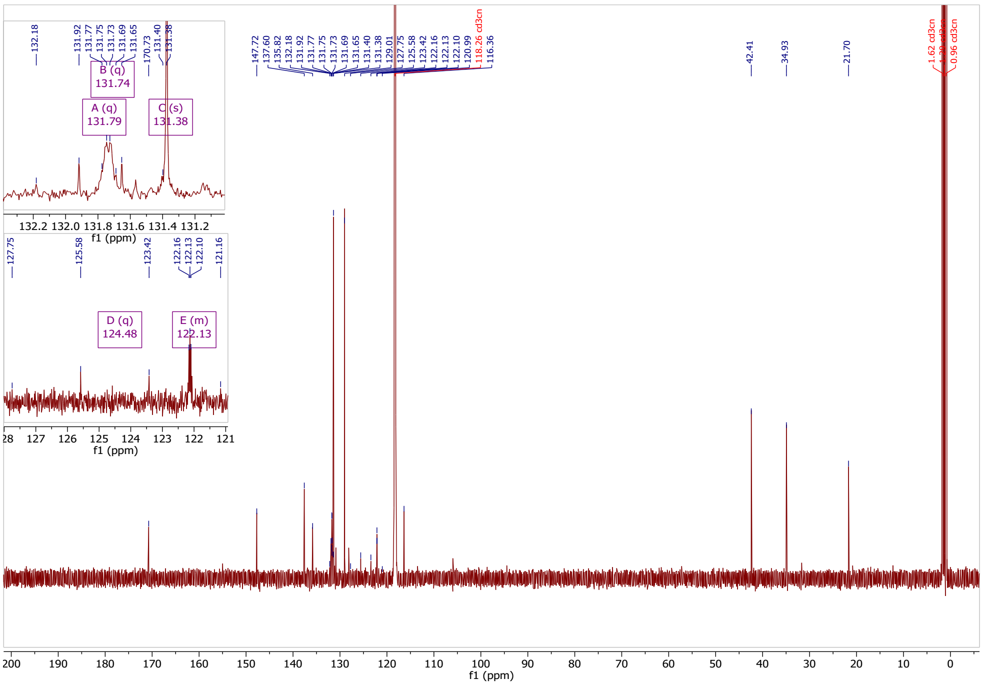

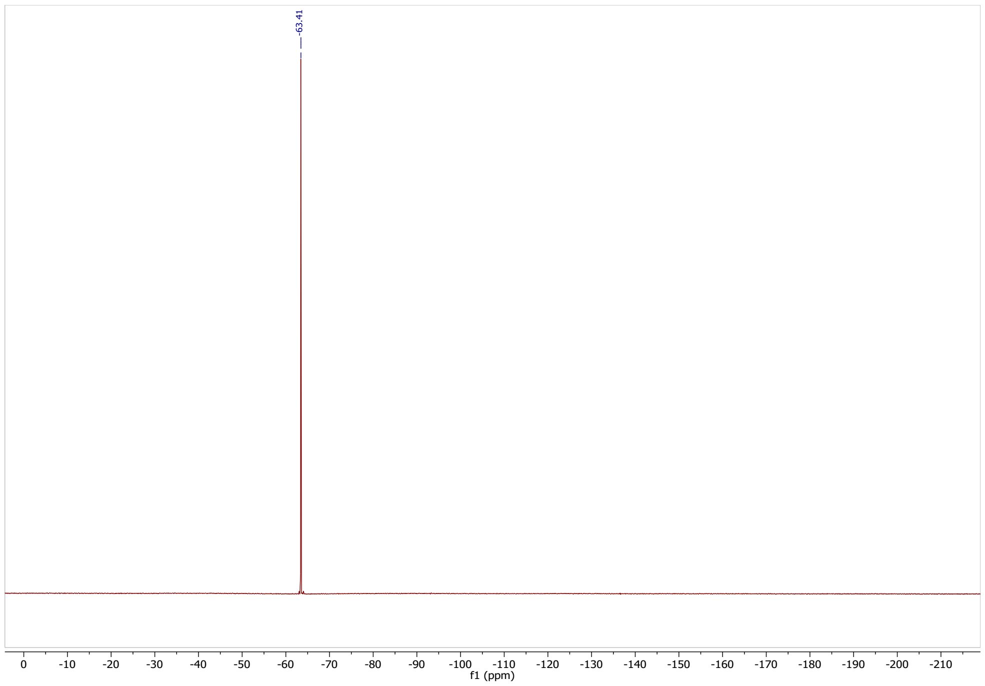

#### Synthesis of 2-((3,5-bis(trifluoromethyl)benzyl)(methyl)amino)-4*H*-benzo[d][1,3]thiazin-4-one (10)

##### *1-(3,5-bis(trifluoromethyl)phenyl)-N-methylmethanamine*

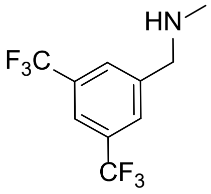

To a stirred solution of aq. methyl amine (40%) (1.93 ml, 14.3 mmol, 5 eq.) and methanol (30 mL) at 50 °C 1-(bromomethyl)-3,5-bis(trifluoromethyl)benzene (0.75 g, 2.86 mmol, 1 eq.) in methanol (5 mL) was added dropwise in 10 min. The reaction mixture was stirred at 50 °C for 24 h. Next, the solvents were concentrated in vacuo and the crude was dissolved in DCM and the organic phase was extracted with 20% NaOH solution (3 x 20 mL) and water (2 x 20 mL) then brine (1 x 20 mL), dried on $Na_2SO_4$, filtered and concentrated. The residue was purified by RP-flash chromatography (MeCN in water, 0.1% HCOOH, 5-100%) resulting in an off-white amorphous solid (0.16 g, 22% yield).

$^1$H NMR (300 MHz, DMSO-*d6*) δ 8.33 (s, 2H), 8.17 (s, 1H), 4.32 (s, 2H), 2.75 (s, 3H).
LC-MS *m/z* calculated for $C_{10}H_9F_6N$: 257; measured $[M+H]^+$: 258.

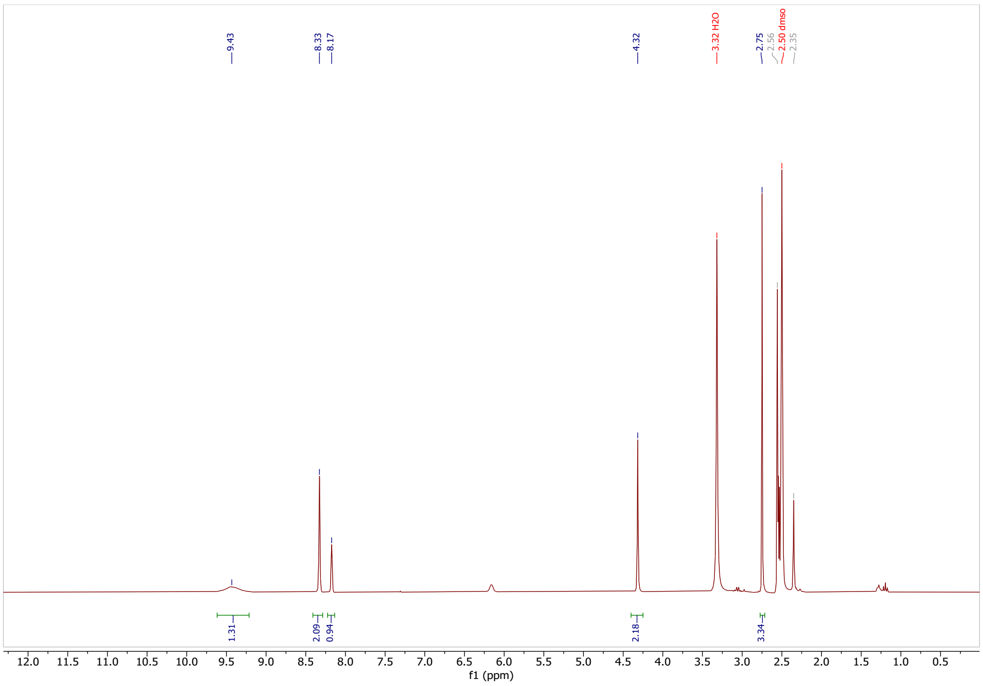

***Methyl 2-(3-(3,5-bis(trifluoromethyl)benzyl)-3-methylthioureido)benzoate***

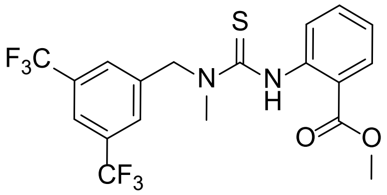

1-(3,5-Bis(trifluoromethyl)phenyl)-*N*-methylmethanamine (0.08 g, 0.32 mmol, 1 eq.) was dissolved in DCM (8 mL) then methyl 2-isothiocyanatobenzoate (0.05 mL, 0.32 mmol, 1 eq.) was added dropwise via a syringe. The mixture was allowed to stir for 3 h at room temperature. The reaction was quenched with 10% HCl (10 mL) and was separated. The aqueous phase was washed with DCM (3 x 10 mL). The combined organic phases were washed with brine (1 x 15 mL) and dried on $Na_2SO_4$, filtered and concentrated in vacuo. The crude product was purified by NP-flash chromatography with hexane/ethyl acetate (0-50%) affording the product as an off-white solid (0.07 g, 50%).

$^1$H NMR (300 MHz, $CDCl_3$) δ 11.13 (s, 1H), 9.01 – 8.81 (m, 1H), 8.01 (dd, J = 8.0, 1.7 Hz, 1H), 7.82 (s, 3H), 7.58 (td, J = 8.7, 7.3, 1.7 Hz, 1H), 7.14 (td, J = 8.3, 7.3, 1.2 Hz, 1H), 5.41 (s, 2H), 3.91 (s, 3H), 3.38 (s, 3H).

LC-MS *m/z* calculated for $C_{19}H_{16}F_6N_2O_2S$: 450; measured $[M+H]^+$: 451.

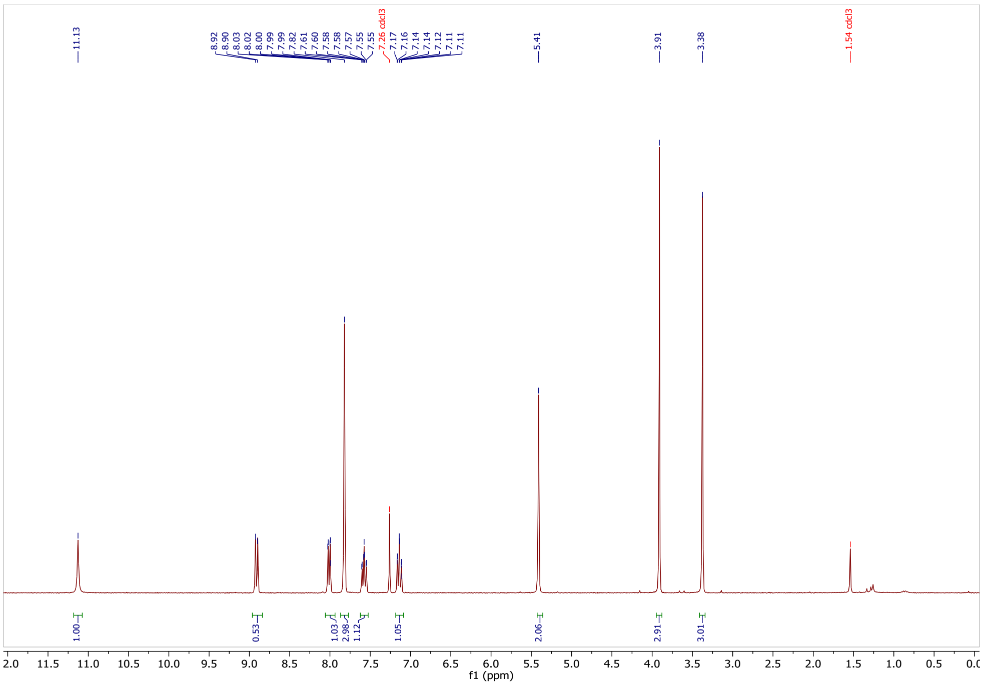

***2-((3,5-bis(trifluoromethyl)benzyl)(methyl)amino)-4H-benzo[d][1,3]thiazin-4-one***

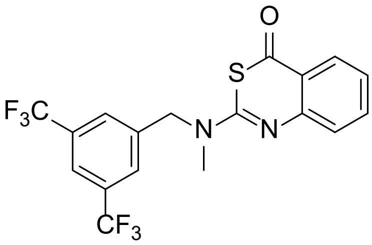

Methyl 2-(3-(3,5-bis(trifluoromethyl)benzyl)-3-methylthioureido)benzoate (0.035 g, 0.077 mmol, 1 eq.) was stirred in concentrated $H_2SO_4$ (1 mL) for 24 h at room temperature. The reaction mixture was poured over ice and neutralized with 10% aq. NaOH solution. Next, ethyl acetate was added, and the organic phase was separated. The aqueous phase was extracted with EtOAc (3 x 10 mL). The combined organic phases were washed with brine (1 x 10 mL) and dried on $Na_2SO_4$, filtered and concentrated in vacuo to obtain the final product as a white amorphous solid (30 mg, 92 % yield).

$^1$H NMR (300 MHz, $CDCl_3$) δ 8.07 (dd, $J$ = 8.0, 1.5 Hz, 1H), 7.83 (s, 1H), 7.80 (s, 2H), 7.64 (td, $J$ = 7.1, 1.7 Hz, 1H), 7.44 (d, $J$ = 7.4 Hz, 1H), 7.23 (td, $J$ = 7.1, 1.2 Hz, 1H), 5.04 (s, 2H), 3.19 (s, 3H).

$^{13}$C NMR (75 MHz, $CDCl_3$) δ 183.44, 157.08, 150.56, 139.76, 136.12, 132.30 (q, $J$ = 33.3 Hz), 128.45, 128.22 – 127.96 (m), 125.12, 124.25, 123.30 (q, $J$ = 272.6 Hz), 122.11 – 121.75 (m), 116.61, 53.34, 36.37.

$^{19}F$ NMR (282 MHz, $CDCl_3$) δ -62.86.

HRMS (ESI/Q-TOF) *m/z*: $[M+H]^+$ Calcd. for $C_{18}H_{13}N_2OF_6S$, 419.0652, measured: 419.0658.

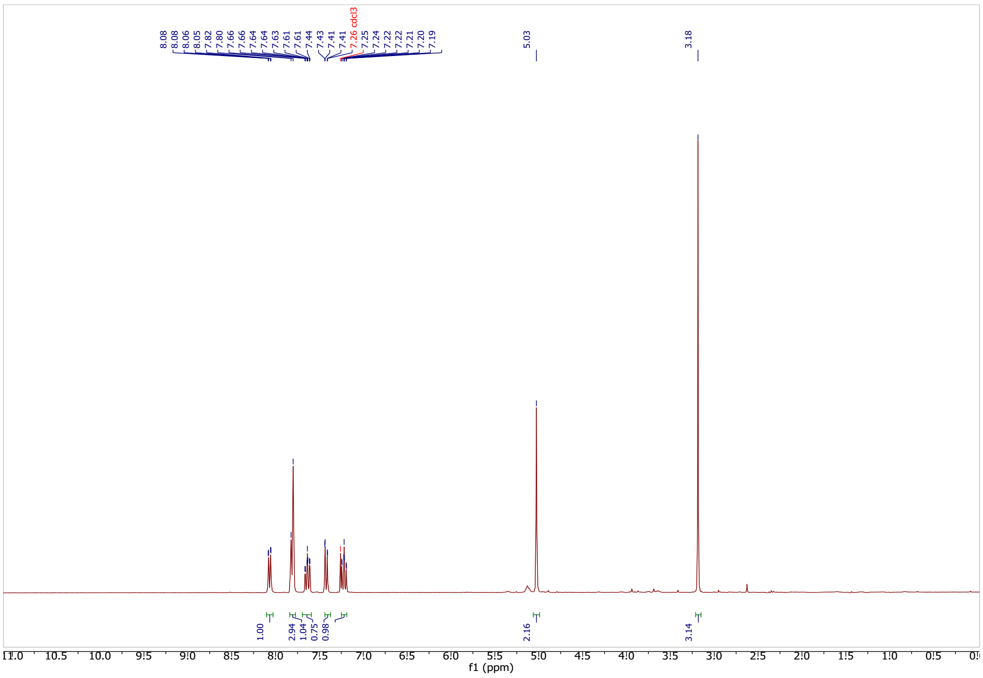

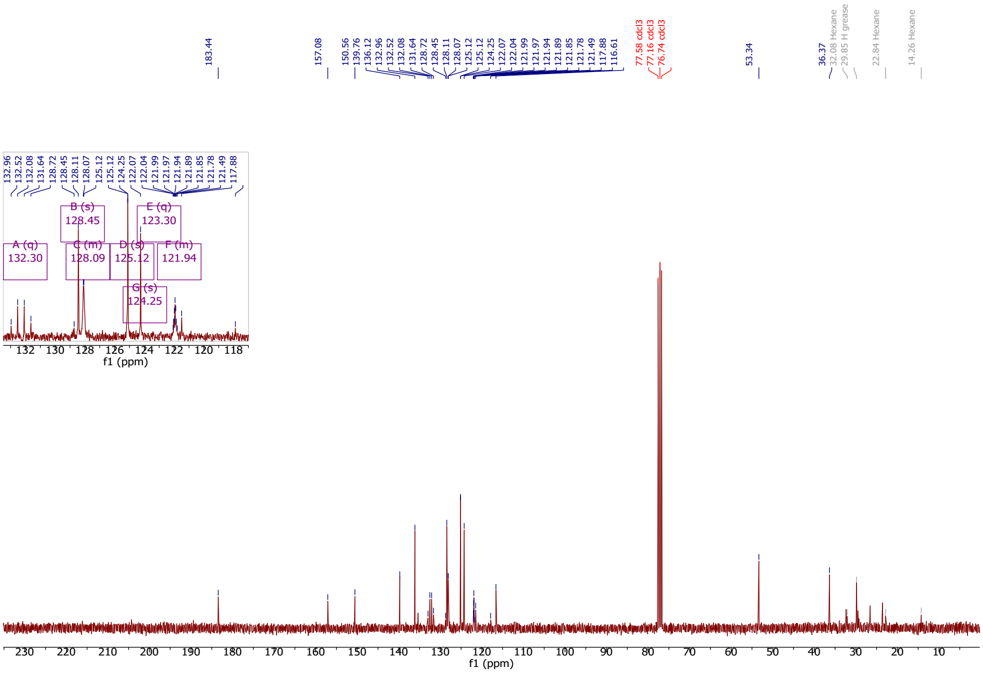

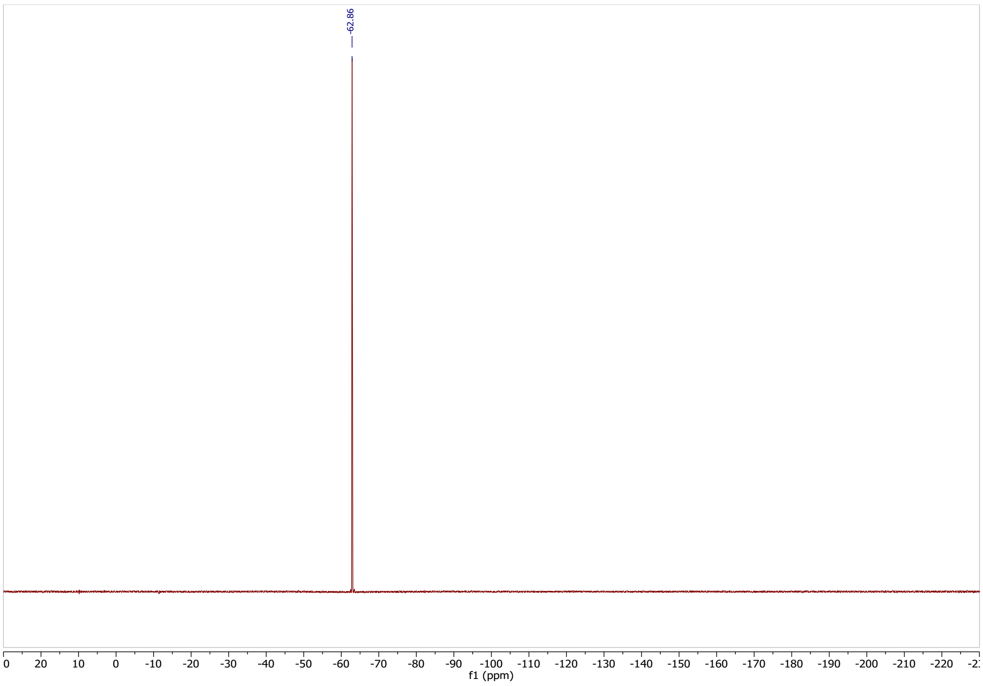

**Synthesis of 2,5-dioxopyrrolidin-1-yl 3,5-bis(trifluoromethyl)benzoate (11)**

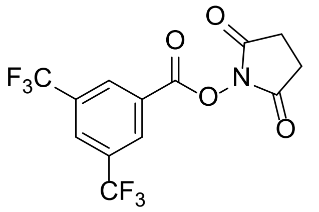

In a flame-dried 20 mL round bottom flask 1-hydroxypyrrolidine-2,5-dione (134 mg, 1.16 mmol, 1 eq.) and 3,5-bis(trifluoromethyl) benzoic acid (300 mg, 1.16 mmol, 1 eq.) was dissolved in dry 1,4-dioxane (5 mL) under argon. To this solution *N, N'*-dicyclohexylcarbodiimide (251 mg, 1.22 mmol, 1.05 eq.) was added. The mixture was warmed up to 50 °C and stirred for 3 h. The resulting suspension was filtered, and the precipitate was resuspended in fresh 1,4-dioxane (5 mL) and stirred for an additional 15 min. The suspension was filtered once more, and the combined supernatants were concentrated in vacuo then purified by normal-phase flash chromatography (hexane/ethyl-acetate 0-40%) to afford a white solid (224 mg, 54 % yield).

$^{1}$H NMR (300 MHz, DMSO-*d6*) δ 8.67 (s, 1H), 8.62 (s, 2H), 2.93 (s, 4H).
$^{13}$C NMR (75 MHz, DMSO-*d6*) δ 169.91, 159.71, 131.58 (q, J = 34.3 Hz), 130.44 (q, J = 3.2 Hz), 129.15 (hept, J = 3.5 Hz), 127.13, 122.55 (q, J = 273.8 Hz), 25.56.
$^{19}$F NMR (282 MHz, DMSO-*d6*) δ -61.55.
NMR spectra are in agreement with the literature [13].

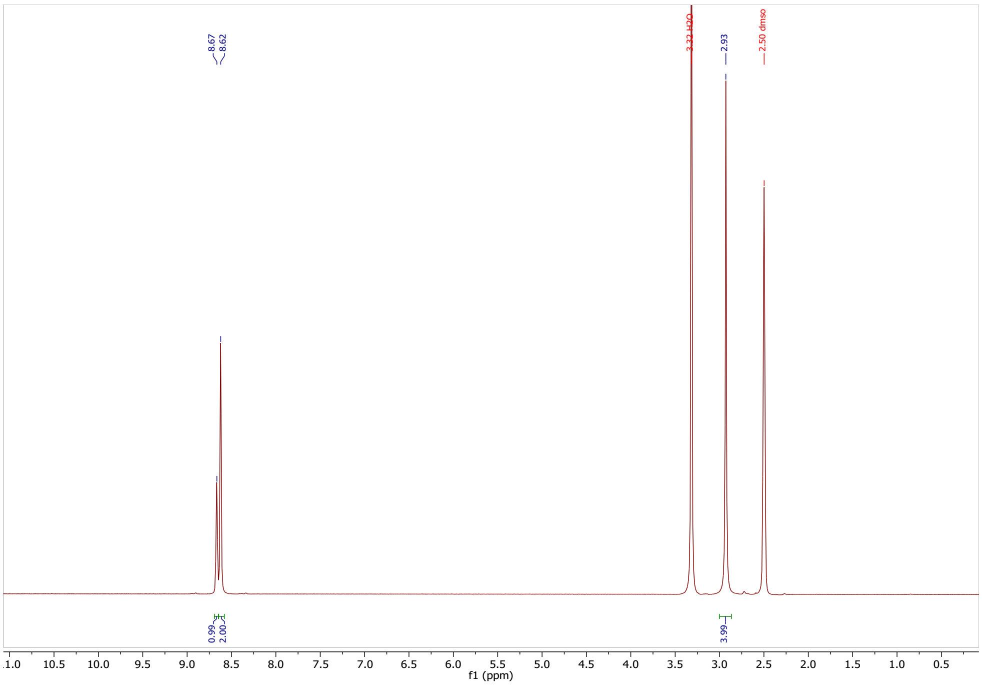
8.67
8.62
3.32 H2O
2.93
2.50 dmso
0.99
2.00
3.99
f1 (ppm)

1

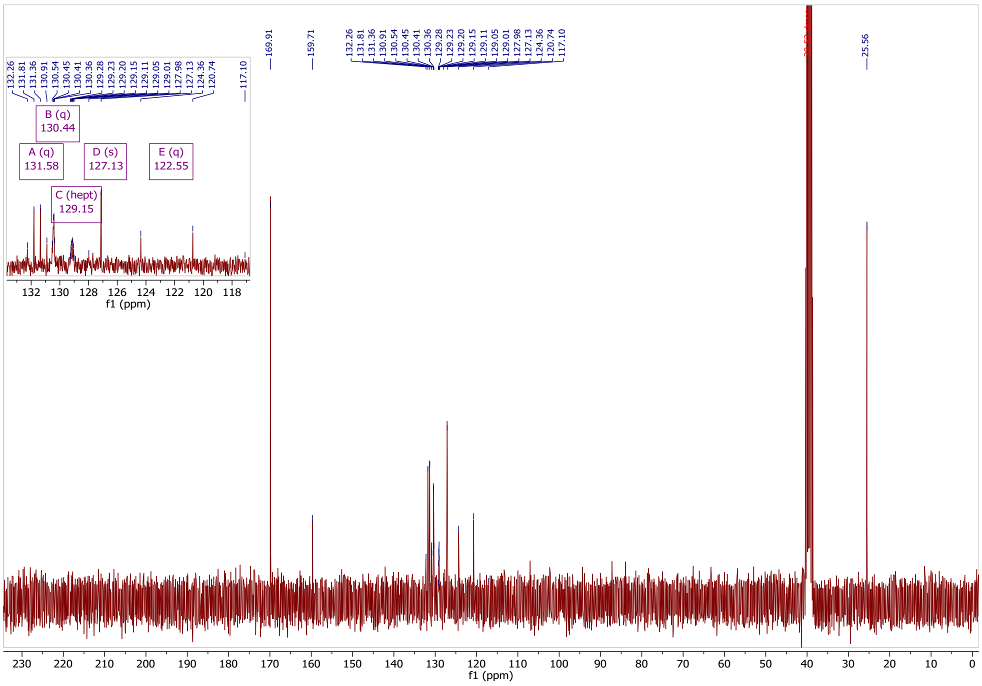
169.91
159.71
132.26
131.81
131.36
130.91
130.54
130.45
130.41
130.36
129.28
129.23
129.20
129.15
129.11
129.05
129.01
127.98
127.13
124.36
120.74
117.10
25.56
A (q)
131.58
B (q)
130.44
C (hept)
129.15
D (s)
127.13
E (q)
122.55
f1 (ppm)

2

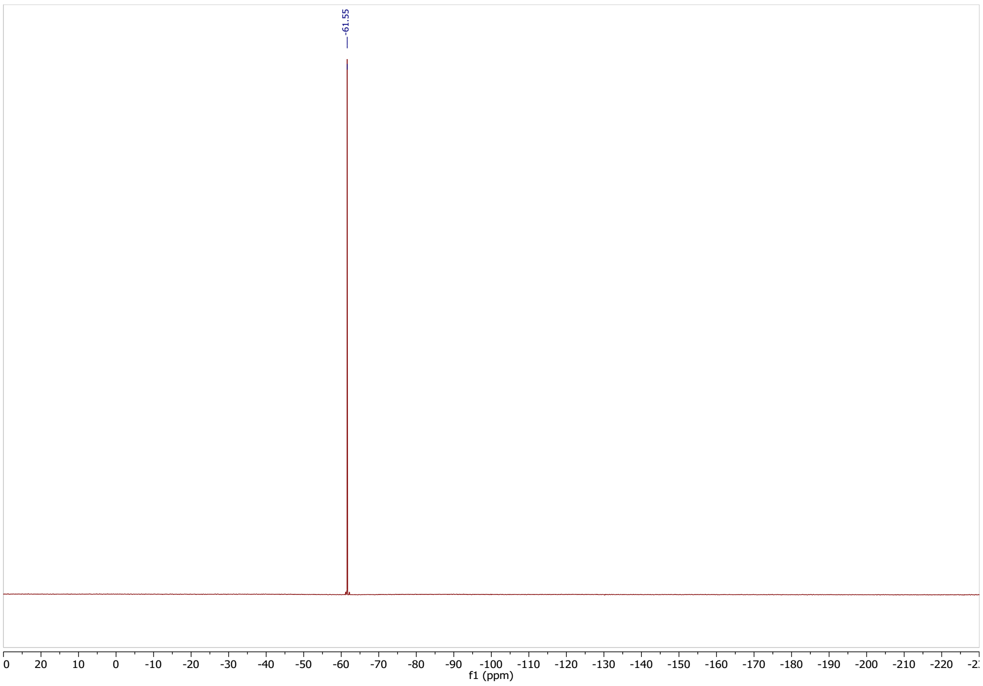

**Synthesis of 2,5-dioxopyrrolidin-1-yl 4-(3,5-bis(trifluoromethyl)phenyl)-4-oxobutanoate (12)**

***4-(3,5-bis(trifluoromethyl)phenyl)-4-oxobutanoic acid***

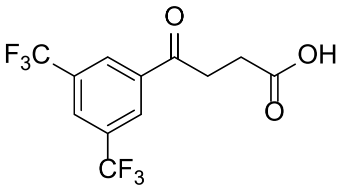

Under argon 1-bromo-3,5-bis(trifluoromethyl)benzene (1.18 mL, 6.8 mmol, 1 eq.) was dissolved in dry THF (10 mL) and cooled to -78 °C. To this mixture n-butyllithium (3 mL, 7.5 mmol, 1.1 eq.; 2.5 M hexane solution) was added dropwise and stirred for 15 minutes. Then furan-2,5-dione (0.82 g, 8.2 mmol, 1.2 eq.) was added to give an orange solution and stirred for 1 h. Next, 1.0 N aq. HCl (25 mL) was added, and the solution was stirred vigorously for 30 minutes at room temperature. The mixture was diluted with EtOAc (20 mL) then separated. The aqueous layer was extracted with additional portions of EtOAc (3 x 20 mL). The combined organic layers were washed with water (50 mL), brine (50 mL), dried over $MgSO_4$, filtered and concentrated in vacuo. The resulting orange liquid was purified by reverse phase chromatography (MeCN in water with 0.1% HCOOH; 5-100%) to afford the product as a white crystal (0.69 g; 32%).

$^1H$ NMR (300 MHz, $CD_3CN$) δ 8.48 (s, 2H), 8.23 (s, 1H), 3.35 (t, *J* = 6.3 Hz, 2H), 2.71 (t, *J* = 6.3 Hz, 2H).

$^{19}F$ NMR (282 MHz, $CD_3CN$) δ -63.45.

LC-MS *m/z*: calculated for $C_{12}H_8F_6O_3$: 314; measured: $[M+H]^+$ 315.

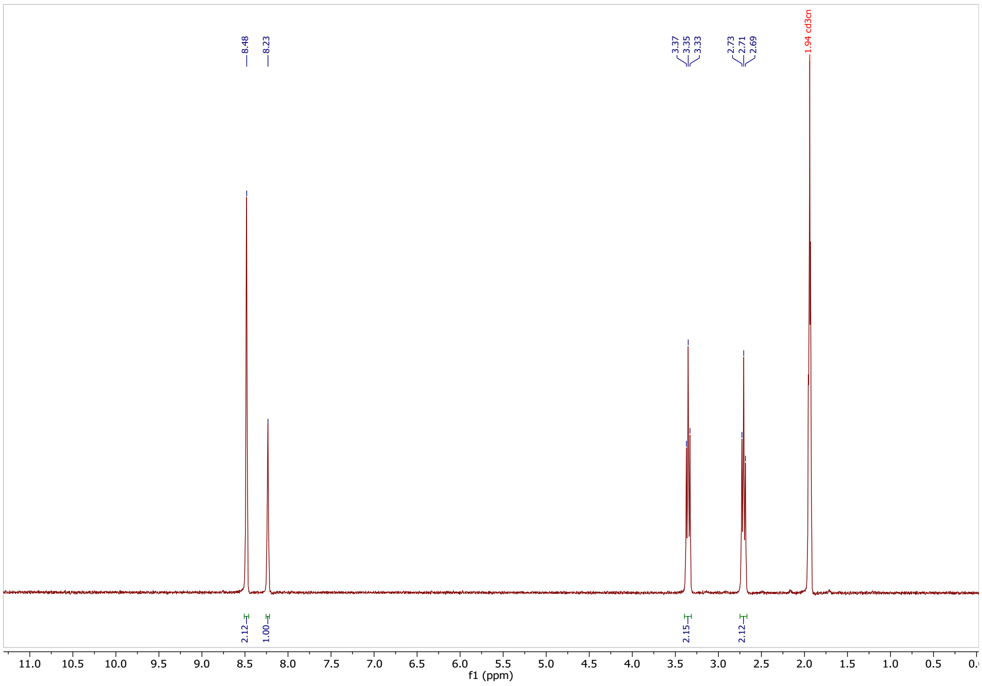

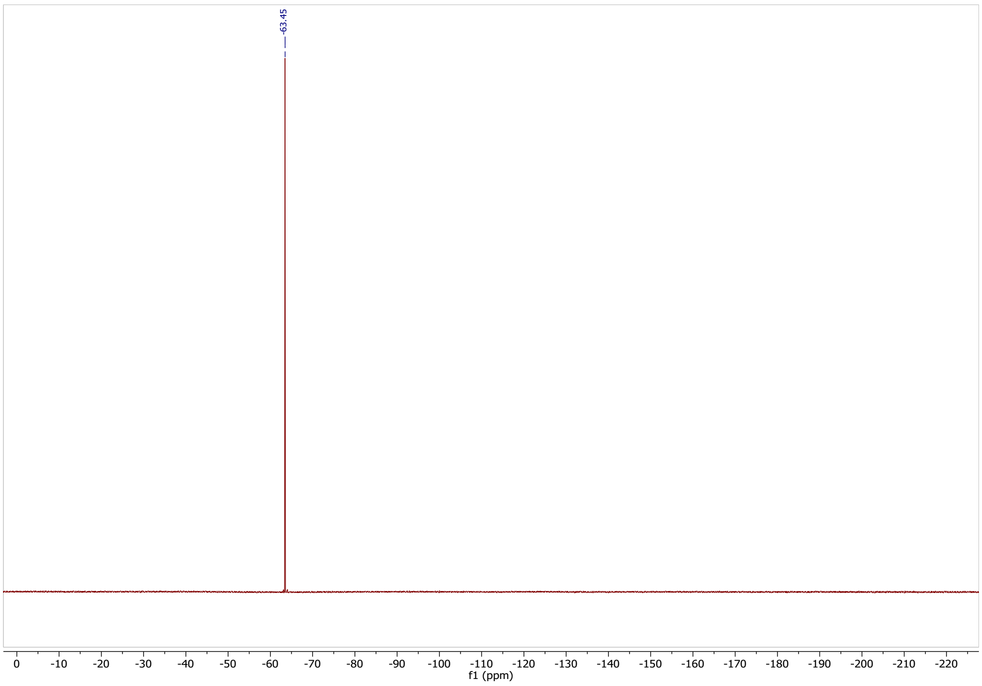

***2,5-dioxopyrrolidin-1-yl 4-(3,5-bis(trifluoromethyl)phenyl)-4-oxobutanoate***

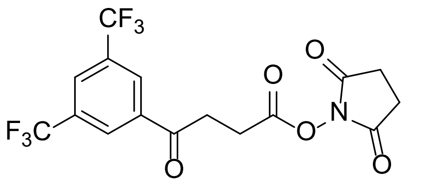

In a flame-dried round bottom flask 4-(3,5-bis(trifluoromethyl)phenyl)-4-oxobutanoic acid (0.13 g, 0.42 mmol, 1 eq.) and 1-hydroxypyrrolidine-2,5-dione (0.048 g, 0.42 mmol, 1 eq.) was dissolved in dry 1,4-dioxane (5 mL). To this solution *N,N'*-dicyclohexylcarbodiimide (0.095 g, 0.46 mmol, 1.1 eq.) was added. The mixture was warmed up to 50 °C and stirred overnight. The resulting suspension was filtered, and the precipitate was resuspended in fresh 1,4-dioxane (5 mL) and stirred for an additional 15 min. The suspension was filtered once more, and the combined supernatants were concentrated via rotary evaporation then purified by normal-phase flash chromatography (hexane/ethyl-acetate 0-40%) to afford a white solid (0.085 g, 50 %).

$^{1}$H NMR (300 MHz, $CD_3CN$) δ 8.49 (s, 2H), 8.25 (s, 1H), 3.52 (t, J = 6.1 Hz, 2H), 3.06 (t, J = 6.4 Hz, 2H), 2.76 (s, 4H).
$^{13}$C NMR (75 MHz, $CD_3CN$) δ 196.43, 170.99, 169.65, 139.04, 132.67 (q, J = 33.8 Hz), 129.84 – 129.21 (m), 127.89 – 127.39 (m), 122.77 (q, J = 272.9 Hz), 34.26, 26.39, 25.83.
$^{19}$F NMR (282 MHz, $CD_3CN$) δ -63.45.
HRMS (ESI/Q-TOF) *m/z*: [M+H]$^{+}$ Calcd. for $C_{16}H_{12}NO_5F_6$, 412.0619, measured: 412.0609

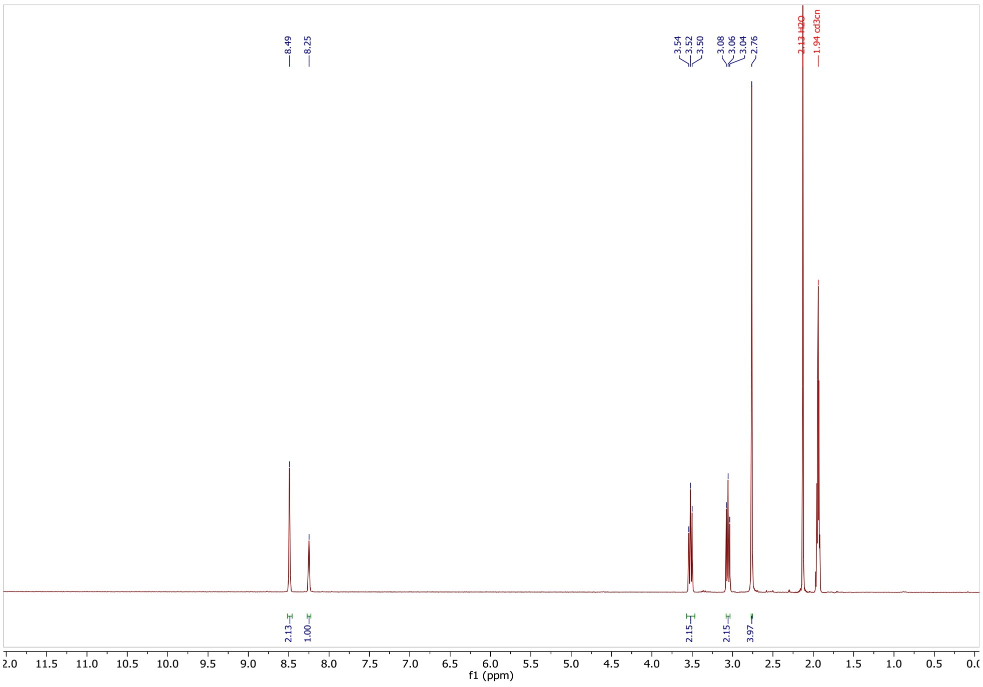

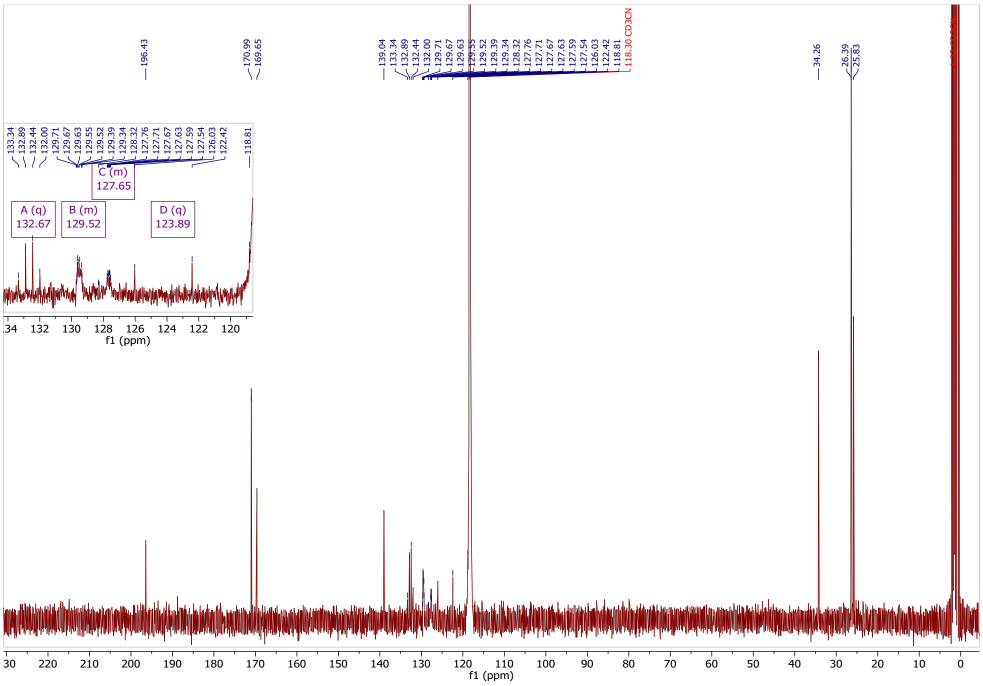

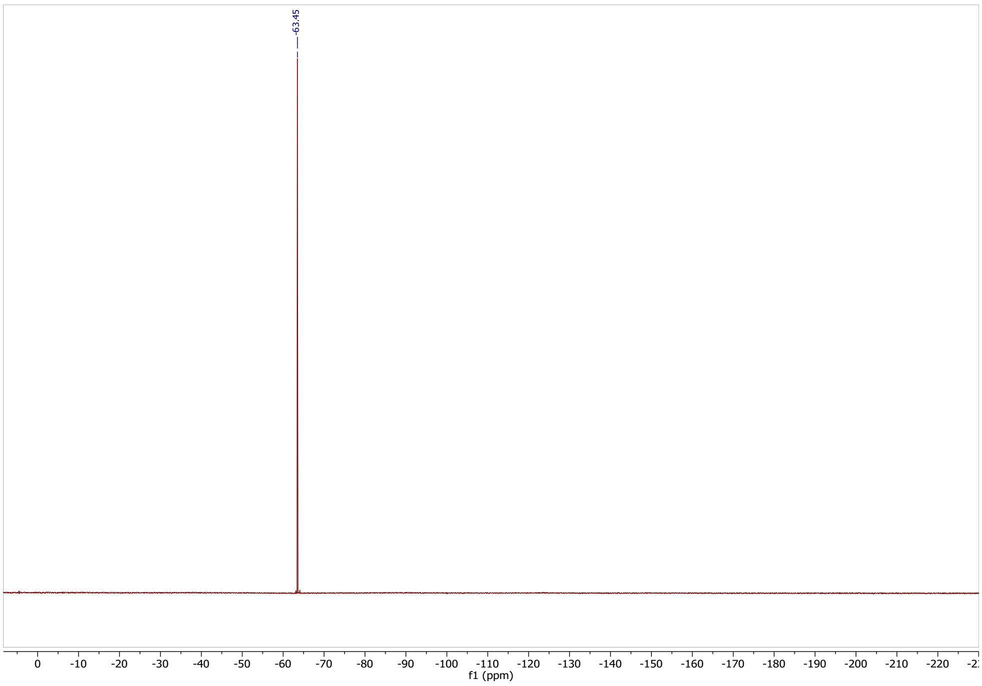

#### Synthesis of *N*-acetyl-*N*-phenyl-3,5-bis(trifluoromethyl)benzamide (14)

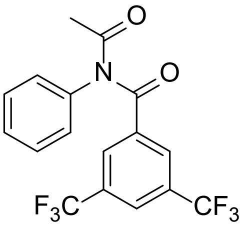

Under argon *N*-phenylacetamide (405 mg, 3.0 mmol, 1 eq.) was dissolved in pyridine (6 mL) and the mixture was cooled down to 0 °C. Next, 3,5-bis(trifluoromethyl)benzoyl chloride (1.1 mL, 6.0 mmol, 2 eqv.) was added dropwise via a syringe. The reaction mixture was allowed to reach room temperature and stirred for 4 h, then cooled to 0 °C and quenched with water (20 mL). The mixture was diluted with EtOAc (10 mL) then separated. The aqueous layer was extracted with additional portions of EtOAc (3 x 10 mL). The combined organic layers were washed with sat. aq. $NaHCO_3$ solution (25 mL) and brine (25 mL), dried over $Na_2SO_4$, filtered and concentrated in vacuo. The crude product was purified by normal-phase flash column chromatography (Hex/EtOAc 0-50%) affording the product as a white crystal (0.46 g, 41% yield).

$^1H$ NMR (300 MHz, $CDCl_3$) δ 7.98 (d, *J* = 1.7 Hz, 2H), 7.90 (s, 1H), 7.46 – 7.35 (m, 3H), 7.22 – 7.16 (m, 2H), 2.40 (s, 3H).

$^{13}C$ NMR (75 MHz, $CDCl_3$) δ 173.10, 169.71, 138.17, 137.31, 131.76 (q, *J* = 34.1 Hz), 129.88, 129.07, 128.68, 125.12 – 124.60 (m), 122.69 (q, *J* = 273.8 Hz), 25.68.

$^{19}F$ NMR (282 MHz, $CDCl_3$) δ -63.11.

HRMS (ESI/Q-TOF) *m/z*: $[M+H]^+$ Calcd. for $C_{17}H_{12}F_6NO_2$, 376.0772, measured: 376.0772.

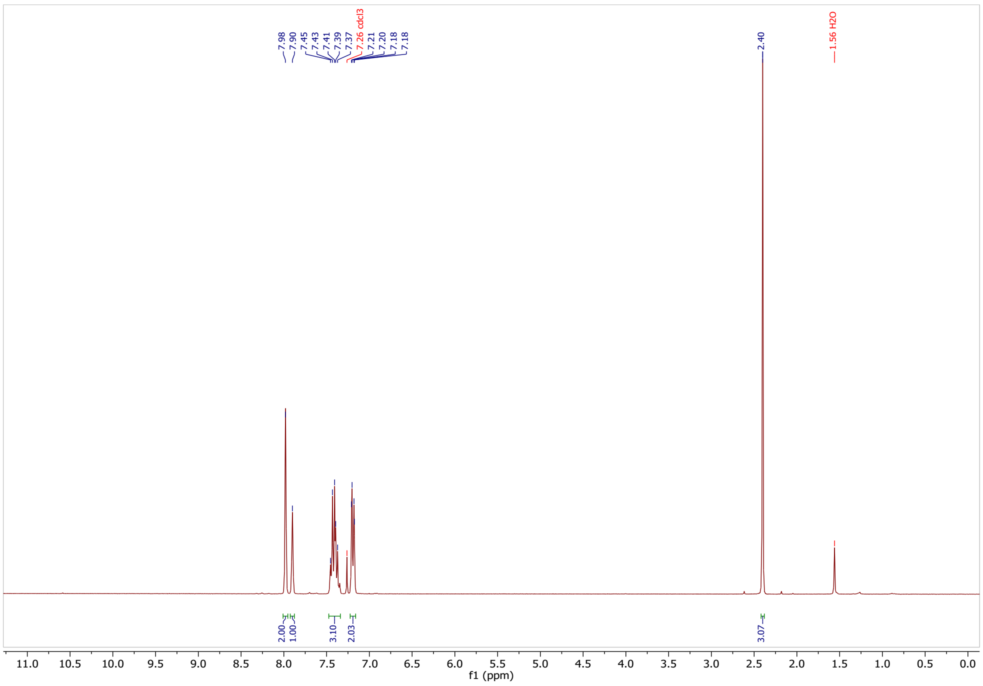

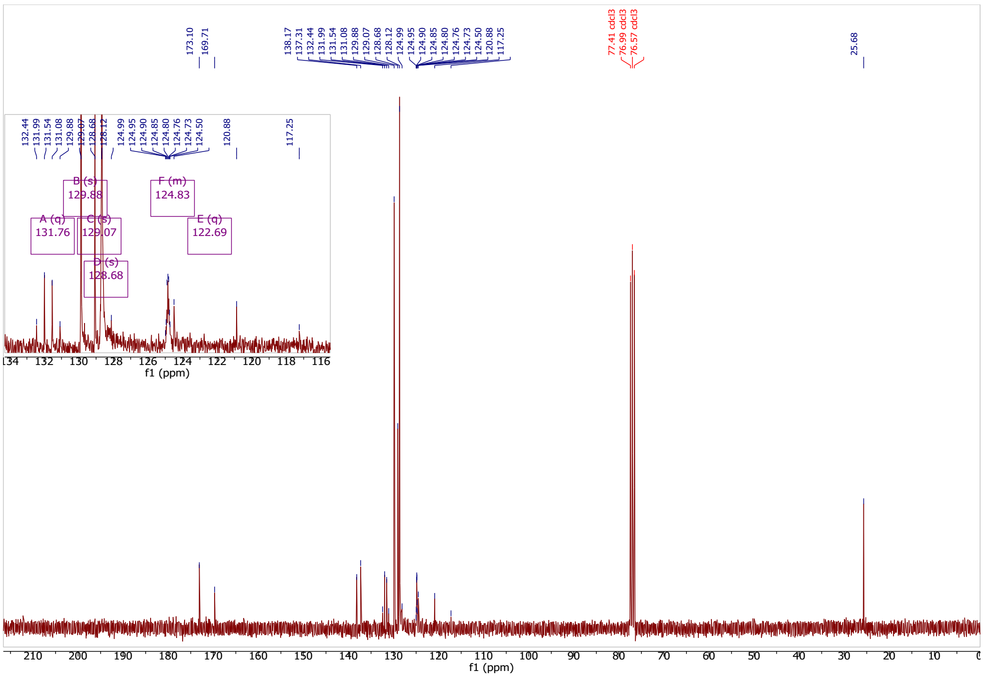

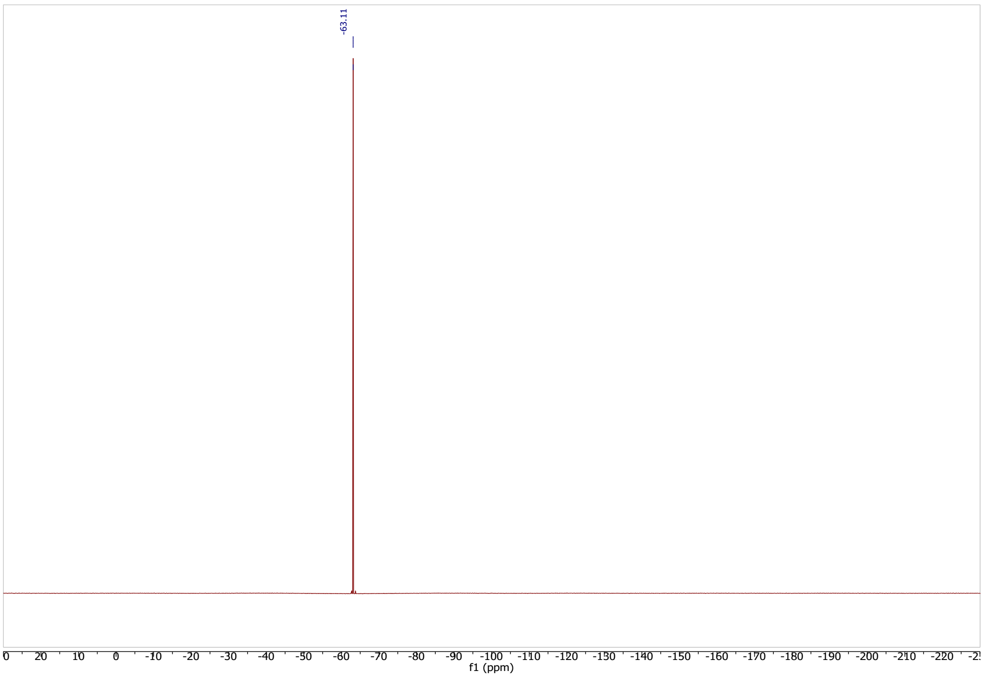

**Synthesis of 1-(3,5-bis(trifluoromethyl)benzyl)-1*H*-benzo[*d*][1,3]oxazine-2,4-dione (16)**

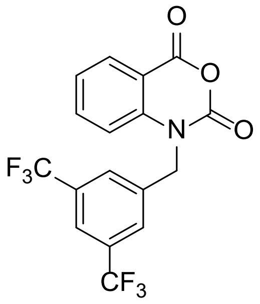

Under argon a solution of 2*H*-benzo[*d*][1,3]oxazine 2,4(1*H*)-dione (80 mg, 0.49 mmol, 1 eq.) in dry DMF (6 ml) was cooled down to 0 °C. Afterwards, NaH (14 mg, 0.59 mmol, 1.2 eq.) and 3,5-bis(trifluoromethyl)benzyl bromide (180.5 mg, 0.59 mmol, 1.2 eq.) was added. The resulting mixture was allowed to warm to RT and stirred overnight. The reaction was poured over crushed ice (10 mL). A precipitate began to form, and it was collected by filtration. The filter cake was washed with cold water (5 ml) and then hexanes (5 ml). The crude product was purified via preparative HPLC resulting in an amorphous off-white solid (30 mg, 16 % yield).

$^{1}H$ NMR (300 MHz, $CDCl_3$) δ 8.22 (dd, J = 7.9, 1.6 Hz, 1H), 7.85 (s, 1H), 7.77 (s, 2H), 7.78 – 7.66 (m, 1H), 7.35 (t, J = 7.6 Hz, 1H), 7.04 (d, J = 8.4 Hz, 1H), 5.41 (s, 2H).

$^{13}C$ NMR (75 MHz, $CDCl_3$) δ 157.84, 148.55, 140.92, 137.71, 137.50, 132.84 (q, J = 33.7 Hz), 131.58, 127.04 (q, J = 3.0 Hz), 124.94, 123.04 (q, J = 272.3 Hz), 122.60 (hept, J = 4.5 Hz), 113.95, 112.07, 47.98.

$^{19}F$ NMR (282 MHz, $CDCl_3$) δ -62.90.

HRMS (ESI/Q-TOF) *m/z*: $[M+H]^+$ Calcd. for $C_{17}H_{10}NO_3F_6$, 390.0564; found 390.0558.

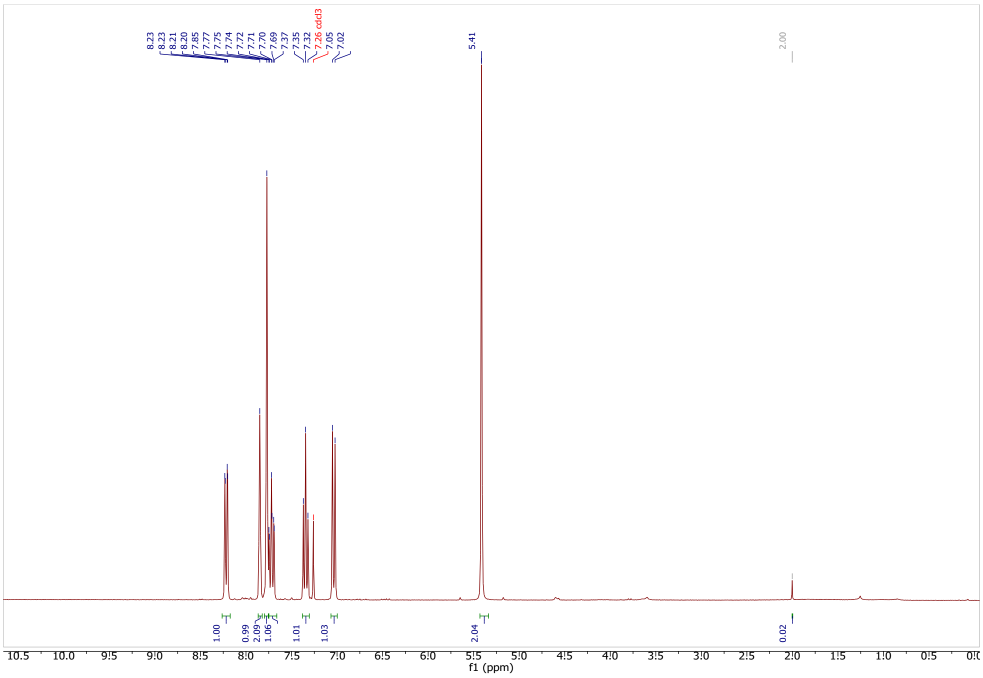

1

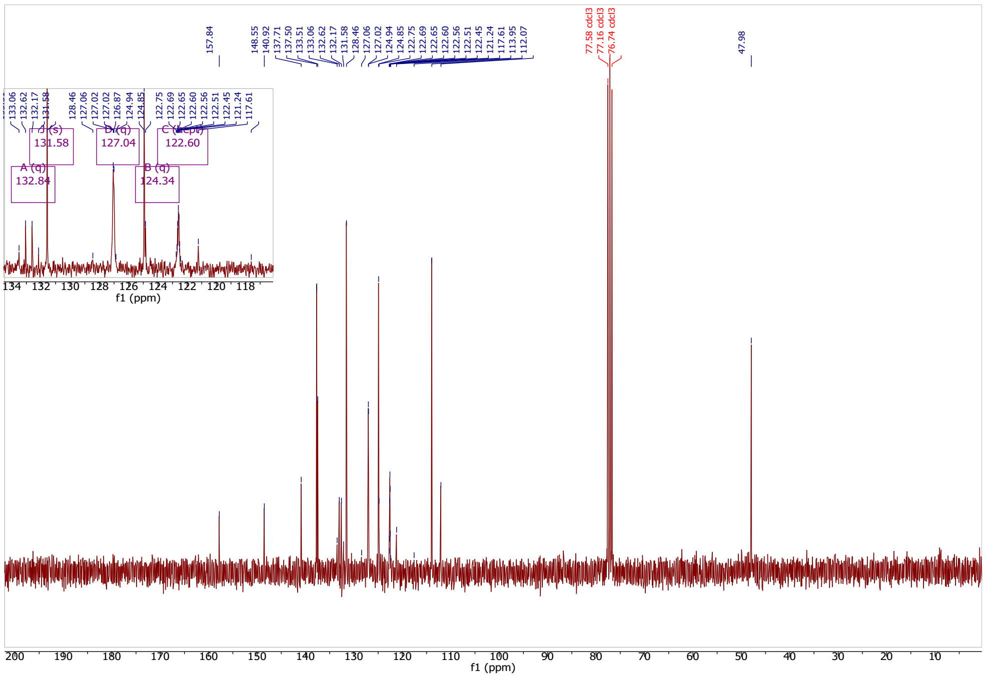

2

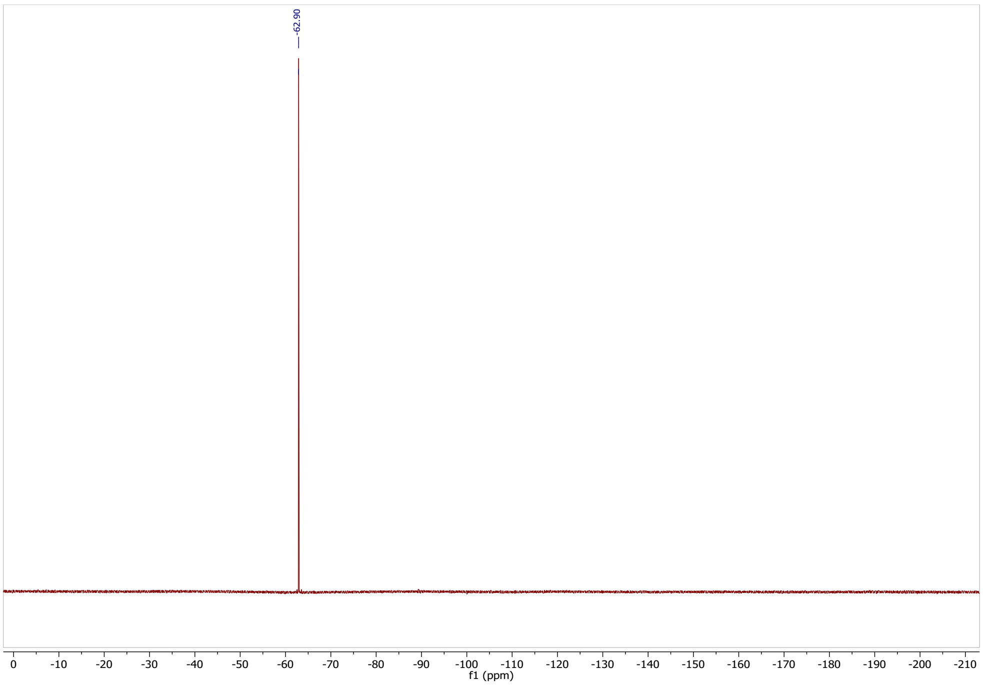

**Synthesis of *N*-(3,5-bis(trifluoromethyl)phenyl)-4,6-dichloro-1,3,5-triazin-2-amine (17)**

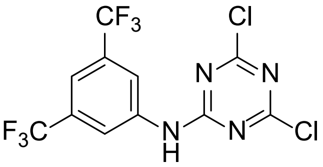

The suspension of 2,4,6-trichloro-1,3,5-triazine (177 mg, 0.97 mmol; 1.1 eq.) and $K_2CO_3$ (121 mg, 0.88 mmol, 1 eq.) in anhydrous THF (5 mL) was cooled to -10 °C. Next, the solution of 3,5-bis(trifluoromethyl)aniline (200 mg, 0.88 mmol, 1 eq.) in anhydrous THF (5 mL) was added dropwise to the mixture in 10 min via a syringe. After the completion of the reaction, the THF was evaporated then the crude product was taken up with water (15 mL) and extracted with ethyl-acetate (2 x 10 mL). The combined organic phases were washed with brine (10 mL) then dried over $Na_2SO_4$, filtered and evaporated in vacuo. The crude product was purified by flash column chromatography in hexane/ethyl-acetate (0-5%) to afford the compound as a white solid (80 mg, 24 %).

$^{1}$H NMR (300 MHz, $CD_3CN$) δ 9.10 (s, 1H), 8.20 (s, 2H), 7.82 (s, 1H).
$^{13}$C NMR (75 MHz, $CD_3CN$) δ 165.67, 139.64, 132.73 (q, *J* = 33.4 Hz), 124.88 (q, *J* = 271.1 Hz), 122.65 – 122.44 (m), 119.66 – 119.05 (m).

$^{19}$F NMR (282 MHz, $CD_3CN$) δ -63.69.

HRMS (ESI/Q-TOF) *m/z*: $[M+H]^+$ Calcd. for $C_{11}H_5N_4F_6Cl_2$, 376.9795; found 376.9788

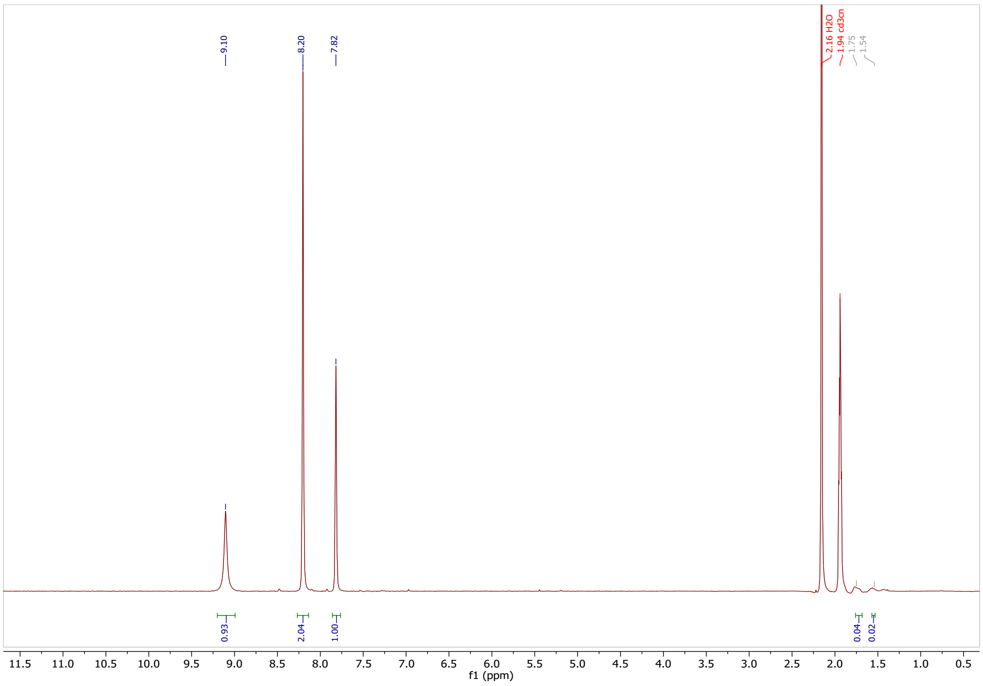

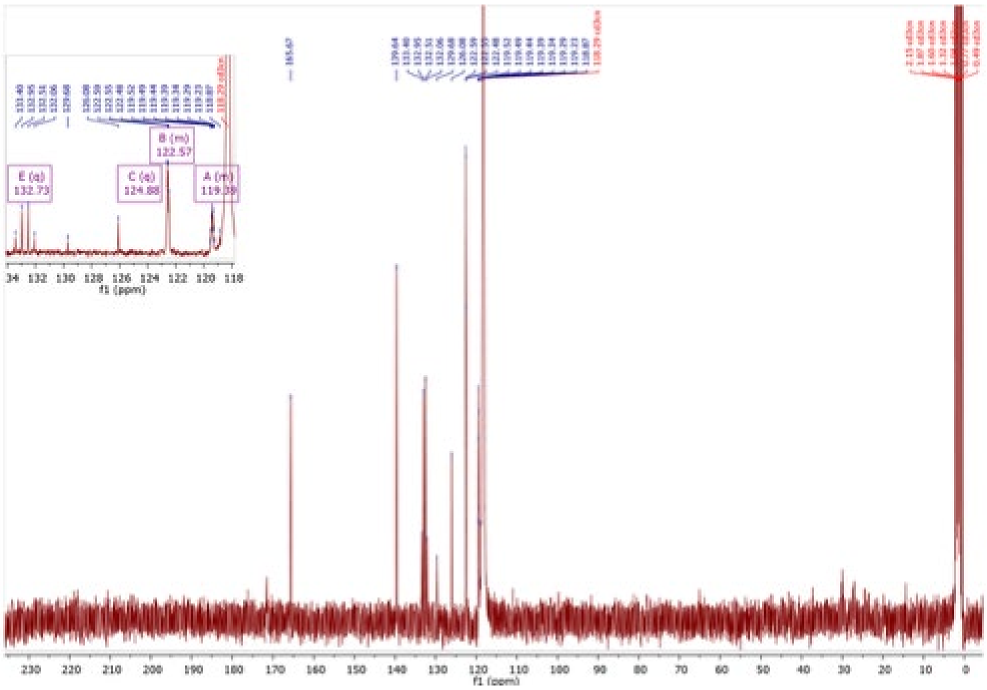

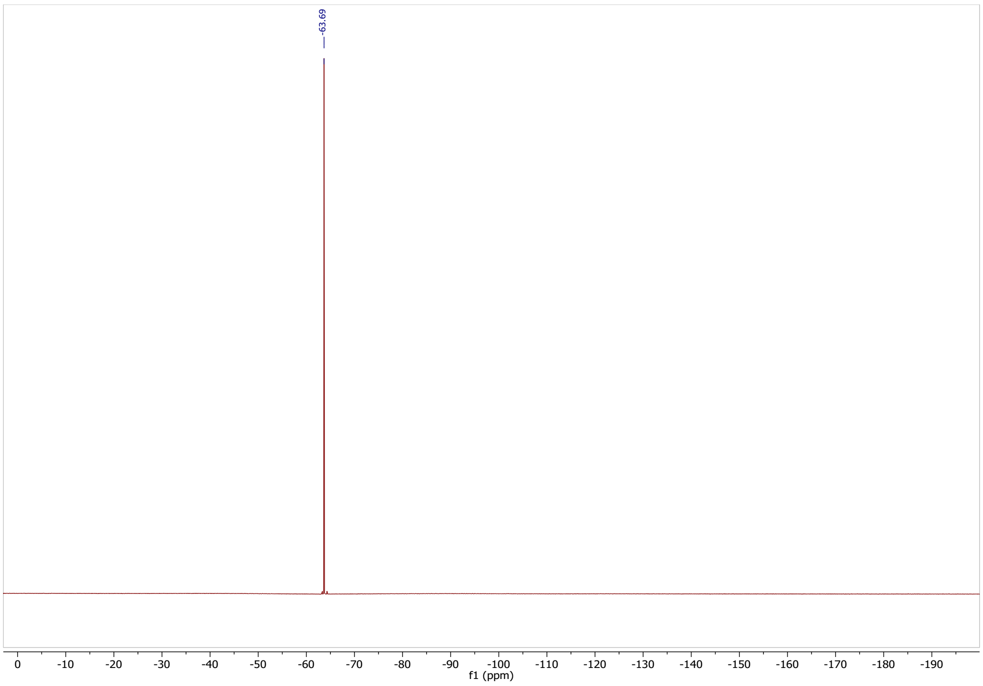

**Synthesis of 1-(2,2-difluoroethenyl)-3,5-bis(trifluoromethyl)benzene (19)**

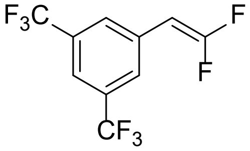

Under argon 3,5-bis(trifluoromethyl)benzaldehyde (400 mg, 1.65 mmol, 1 eq.) and $PPh_3$ (650 mg, 2.48 mmol, 1.5 eq.) was dissolved in dry DMF (7 mL) and the mixture was heated up to 90 °C then a solution of potassium 2-bromo-2,2-difluoroacetate (633 mg, 2.97 mmol, 1.8 eq.) in 3 mL of dry DMF was added dropwise to the mixture via a syringe with the rate of addition controlling the evolution of the $CO_2$ gas. Afterwards, the rection was stirred for 1 h at 90 °C. Upon completion, the reaction was cooled to 0 °C and then quenched with $H_2O$ (10 mL). Subsequently, $Et_2O$ (10 mL) was added to the mixture, and the organic layer was washed with $H_2O$ (3 x 10 mL) followed by aq. LiCl (10% in $H_2O$; 10 mL). Subsequently, MeI (351 mg, 2.48 mmol, 1.5 eq.) was added to the organic layer, and the mixture was stirred at room temperature for 30 min to methylate the residual $PPh_3$. The organic layer was washed with $H_2O$ (3 x 10 mL) and brine (10 mL) then dried over $Na_2SO_4$. After filtration and evaporation of the solvent, the crude product was purified by normal-phase flash chromatography (hexane/ethyl-acetate 0-8 %) to furnish the desired product as a colorless liquid (44 mg, 10 % yield).

$^1$H NMR (300 MHz, $CDCl_3$) δ 7.76 (s, 3H), 5.40 (dd, *J* = 25.0, 3.1 Hz, 1H).

$^{13}$C NMR (75 MHz, CDCl3) δ 157.30 (dd, $J$ = 300.5, 292.5 Hz), 132.85 (dd, $J$ = 7.8, 6.3 Hz), 132.38 (q, $J$ = 33.6 Hz), 127.64 – 127.39 (m), 124.16 (q, $J$ = 273.2 Hz), 120.87 (hept, $J$ = 3.3 Hz), 81.15 (dd, $J$ = 31.4, 13.2 Hz).

$^{19}$F NMR (282 MHz, CDCl3) δ -63.29, -78.16 – -78.42 (m), -79.84 (dd, $J$ = 21.8, 2.9 Hz).

GC-MS (FID) m/z: calculated for $C_{10}H_4F_8$: 276; measured: 276.

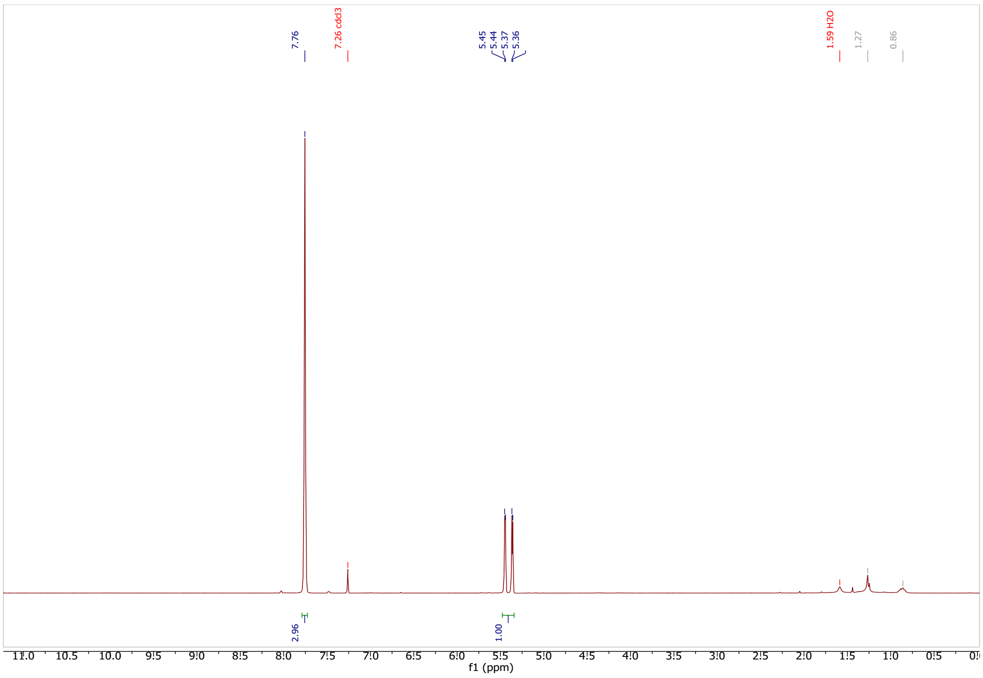

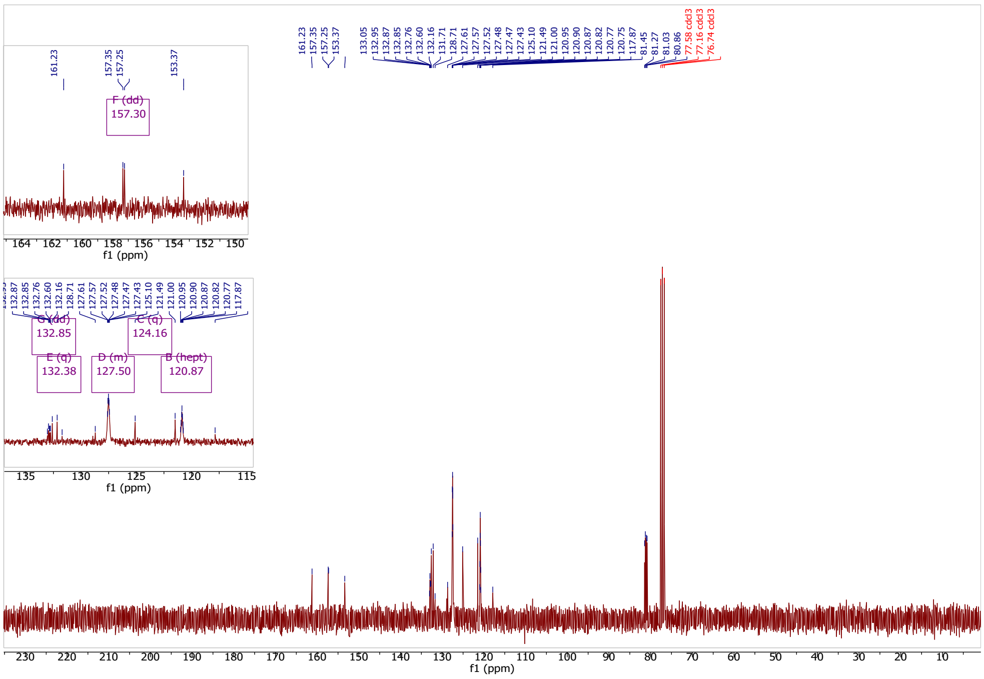

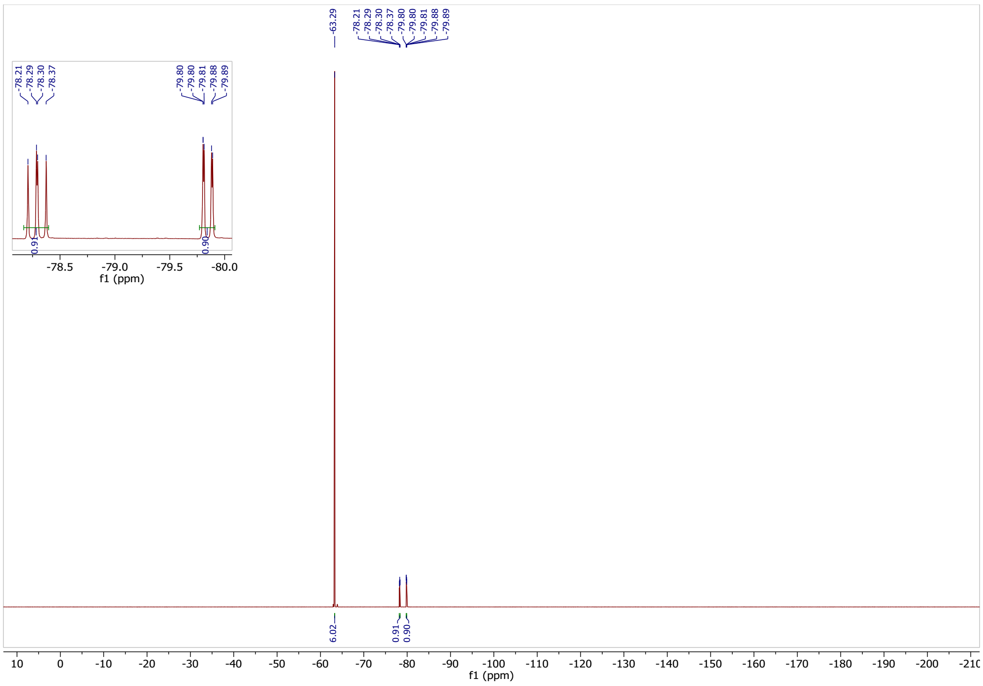

**Synthesis of 2-hydroxy-4,6-bis(trifluoromethyl)benzaldehyde (25)**

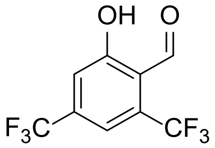

2-Methoxy-4,6-bis(trifluoromethyl)benzaldehyde (100 mg, 0.37 mmol, 1 eq.) was dissolved in dry DCM (5 ml), cooled to -78 °C and treated with $BBr_3$ (1 M solution in DCM, 0.4 ml, 0.40 mmol, 1.1 eq.). The reaction mixture was stirred at -78 °C for 30 min then allowed to reach room temperature and stirred overnight. After completion of the reaction, it was quenched with water (10 ml) and stirred for 30 min. The organic phase was diluted with DCM (5 ml) then separated and washed with brine (5 mL), dried on $Na_2SO_4$. After filtration, the organic phase was evaporated at room temperature resulting in the desired product as a yellow oil (72 mg, 76% yield).

$^1$H NMR (300 MHz, $CDCl_3$) δ 12.26 (s, 1H), 10.34 (q, $J$ = 1.8 Hz, 1H), 7.51 (s, 3H).
LC-MS *m/z*: calculated for $C_9H_4F_6O_2$: 258; measured: $[M-H]^-$ 257.
NMR spectra are in agreement with the literature [14].

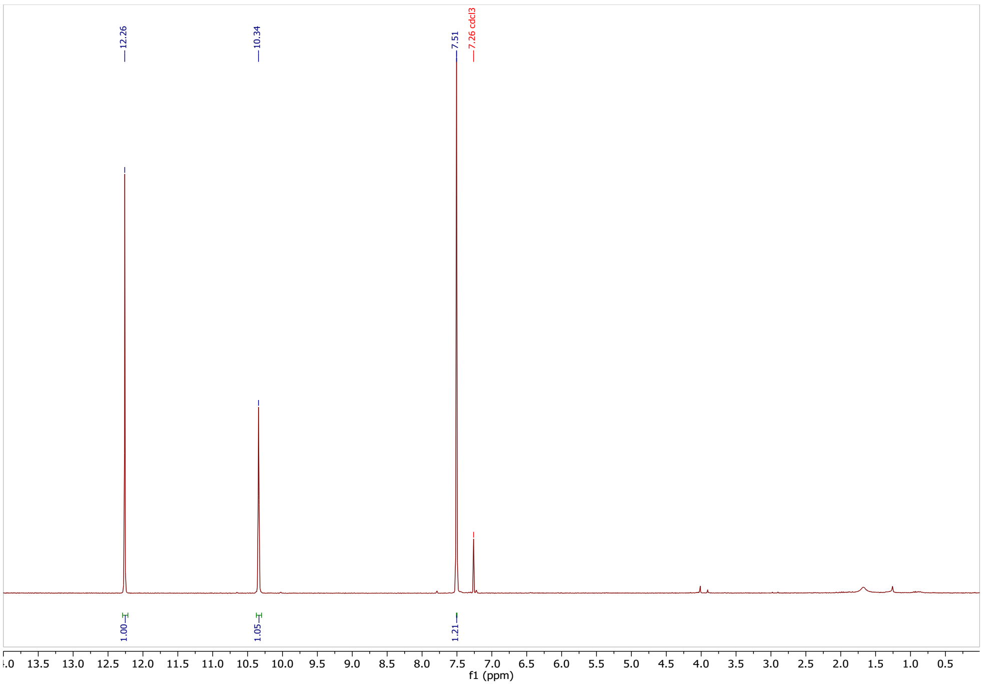

**Synthesis of 2-ethynyl-5-(trifluoromethyl)benzaldehyde (31)**

***5-(trifluoromethyl)-2-((trimethylsilyl)ethynyl)benzaldehyde***

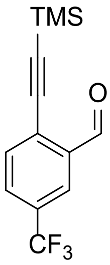

Under argon CuI (0.07 g, 0.27 mmol, 0.05 eq.), $Pd(PPh_3)_4$ (0.086 g, 0.074 mmol, 0.01 eq.) was dissolved in degassed TEA (3 mL) and cooled down to 0 °C. To this mixture a solution of 2-bromo-5-(trifluoromethyl)benzaldehyde (1 mL, 7.4 mmol, 1 eq.) in anhydrous THF (4 mL) was added. Next, ethynyltrimethylsilane (1.6 mL, 11.1 mmol, 1.5 eq.) was added dropwise and stirred at 0 °C for 30 min then the mixture was warmed up to 45 °C and stirred for 2 h. After cooling down to room temperature the mixture was filtered through a celite pad and washed with THF. The filtrate was concentrated in vacuo. The crude product was purified by normal-phase flash chromatography (Hex/EtOAc 0-10%) affording the product, 5-(trifluoromethyl)-2-((trimethylsilyl)ethynyl)benzaldehyde as a white crystal (1.6 g, 80 % yield).

$^{1}H$ NMR (300 MHz, $CDCl_3$) δ 10.56 (s, 1H), 8.16 (s, 1 H), 7.88 – 7.52 (m, 2 H), 0.30 (s, 9H).
$^{19}F$ NMR (282 MHz, $CDCl_3$) δ -63.22.

LC-MS *m/z* calculated for $C_{13}H_{13}F_3OSi$: 270; measured $[M+H]^+$: 271.

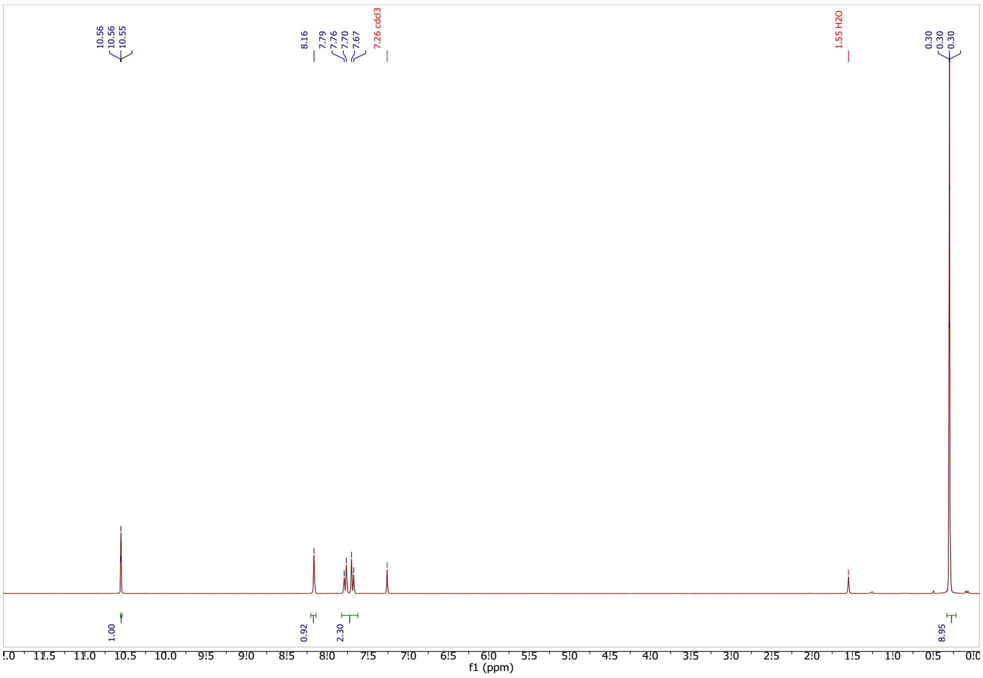

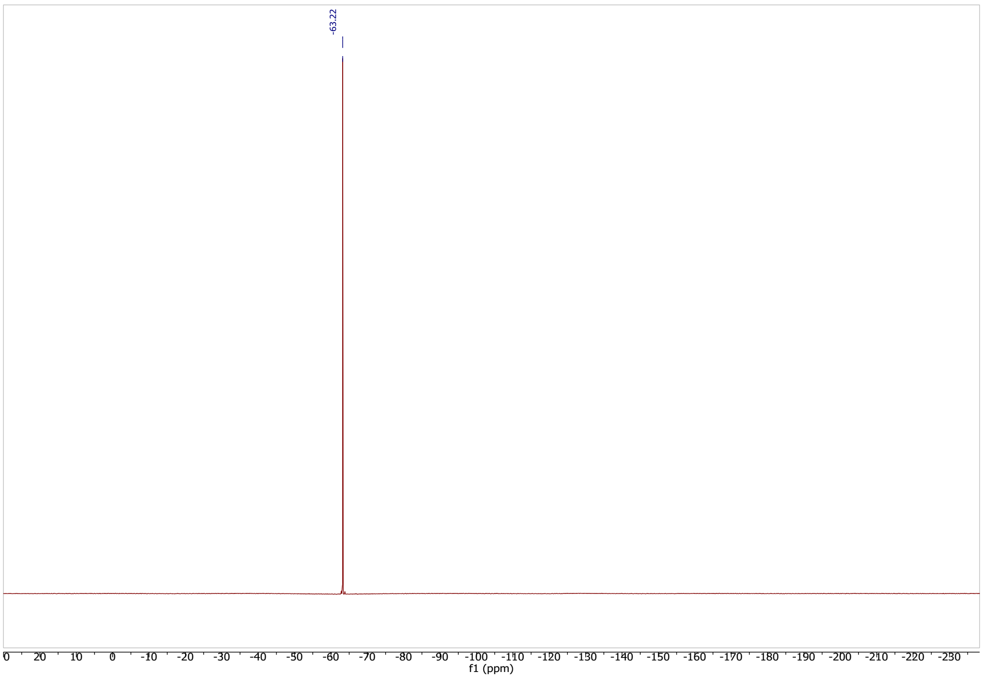

***2-ethynyl-5-(trifluoromethyl)benzaldehyde***

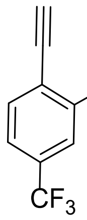

In a flask 5-(trifluoromethyl)-2-((trimethylsilyl)ethynyl)benzaldehyde (51.2 mg, 0.19 mmol, 1 eq.) was dissolved in methanol (5 mL) and $K_2CO_3$ (2.62 mg, 0.02 mmol, 0.01 eq.) was added and stirred at room temperature for 30 min. The mixture was diluted with water (20 mL) and extracted 3 times with DCM (3 x 10 mL). The combined organic phases were washed with brine (10 mL) and dried over $Na_2SO_4$, filtered and concentrated in vacuo affording the product as a white solid (19.9 mg, 53%).

$^{1}$H NMR (300 MHz, $CDCl_3$) δ 10.55 (s, 1H), 8.20 (s, 1H), 7.84 – 7.71 (m, 2H), 3.61 (s, 1H).
$^{13}$C NMR (75 MHz, $CDCl_3$) δ 189.97, 137.01, 134.66, 131.59 (q, $J$ = 33.5 Hz), 130.10 (q, $J$ = 3.6 Hz), 126.93 (q, $J$ = 273.3 Hz), 124.53 (q, $J$ = 4.0 Hz), 86.95, 86.92.
$^{19}$F NMR (282 MHz, $CDCl_3$) δ -63.29.
HRMS (ESI/Q-TOF) $m/z$: $[M+H]^+$ Calcd. for $C_{10}H_4OF_3$, 197.0214, measured: 197.0216.

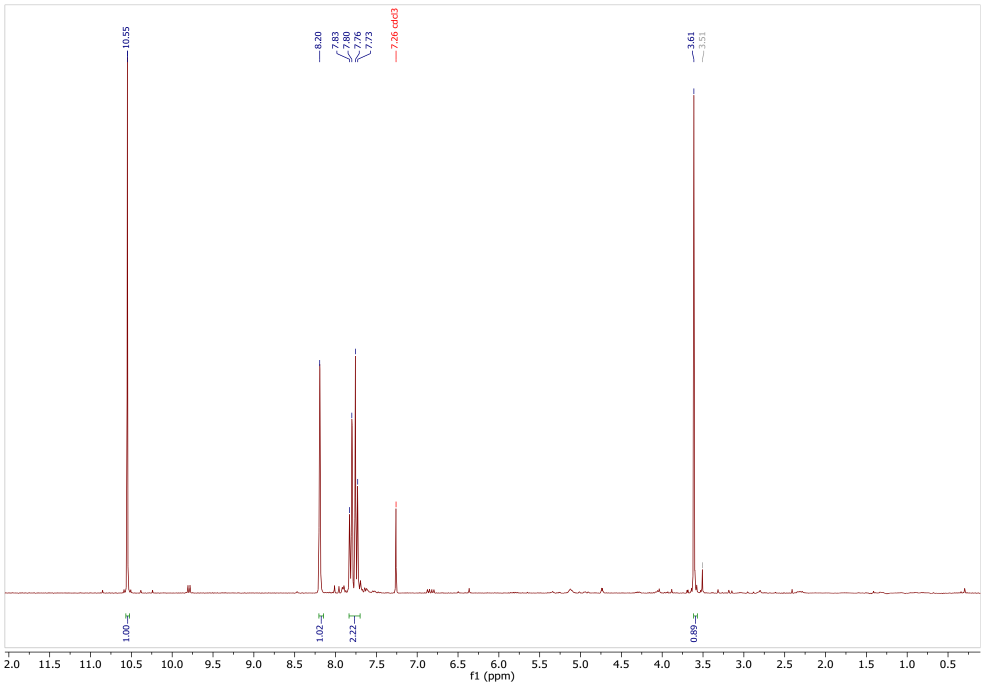

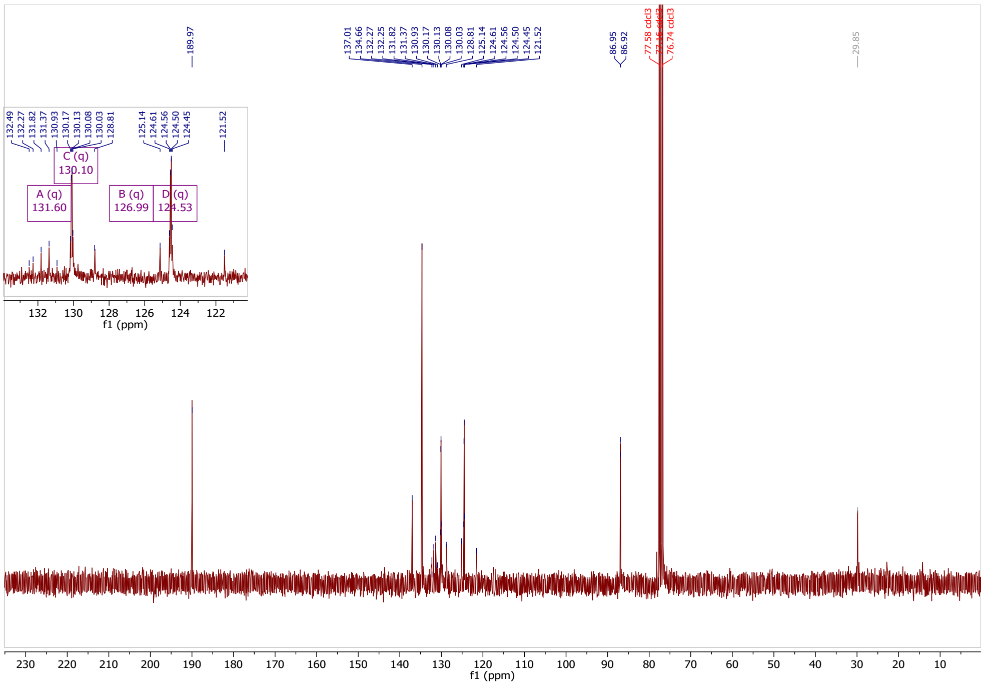

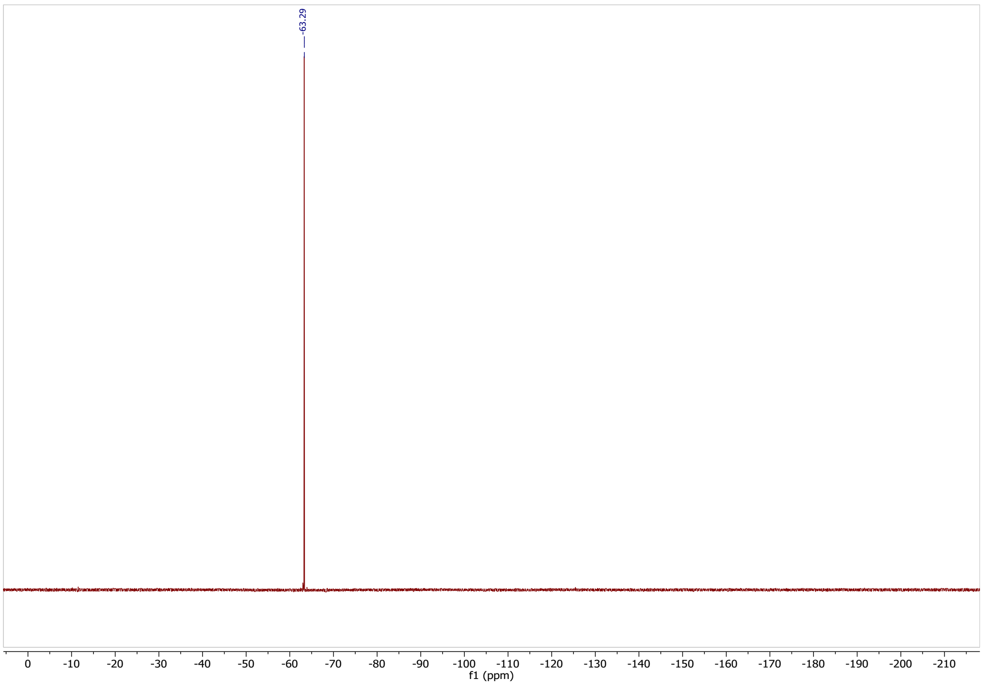

**Synthesis of 3-((3,5-bis(trifluoromethyl)phenyl)amino)-4-methoxycyclobut-3-ene-1,2-dione (33)**

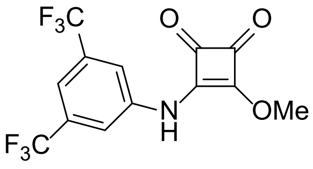

3,4-Dimethoxy-3-cyclobutene-1,2-dione (100 mg, 0.70 mmol, 1 eq.) was dissolved in MeOH (5 mL) and 3,5-bis(trifluoromethyl)aniline (0.12 mL, 0.77 mmol, 1.1 eq.) was added. The mixture was stirred at room temperature for 48 h. The resulting precipitate was filtered and washed with cold MeOH (2 x 2 mL) and dried in vacuo to obtain a bright yellow amorphous solid (42.7 mg, 16%).

$^1$H NMR (300 MHz, DMSO-*d6*) δ 11.18 (s, 1H), 8.04 (s, 2H), 7.78 (s, 1H), 4.41 (s, 3H).
$^{13}$C NMR (75 MHz, DMSO-*d6*) δ 187.39, 184.48, 179.88, 169.12, 140.18, 131.14 (q, *J* = 33.1 Hz), 123.05 (q, *J* = 273.1 Hz), 119.34 (q, *J* = 3.4 Hz), 116.25 (hept, *J* = 3.0 Hz), 60.92.
HRMS (ESI/Q-TOF) *m/z*: $[M+H]^+$ Calcd. for $C_{13}H_8O_3NF_6$, 340.0403, measured: 340.0407.
NMR spectra are in agreement with the literature [15].

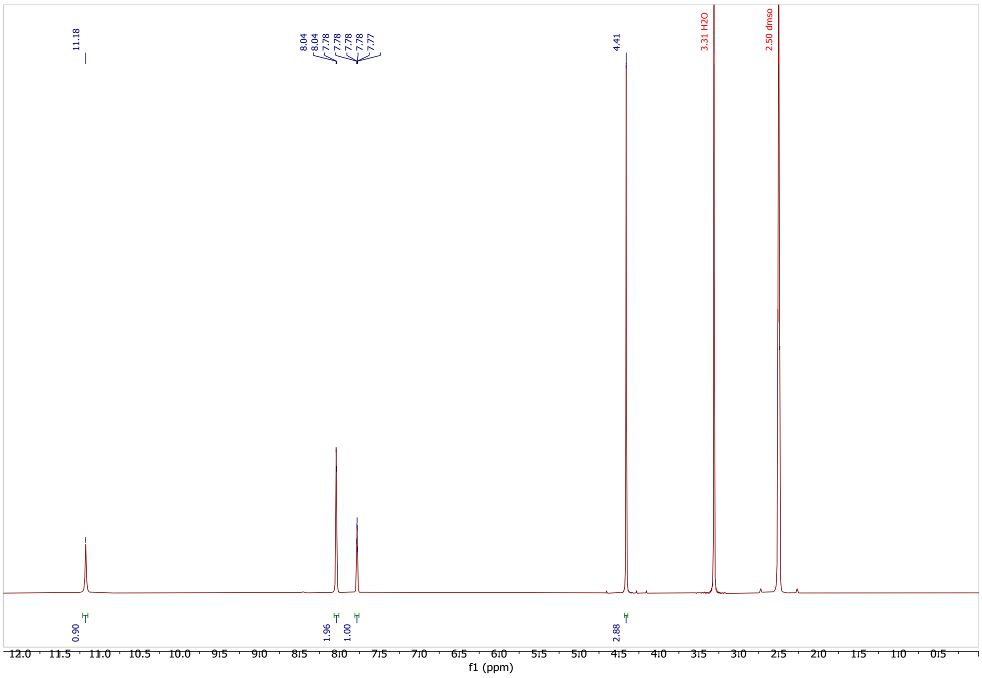

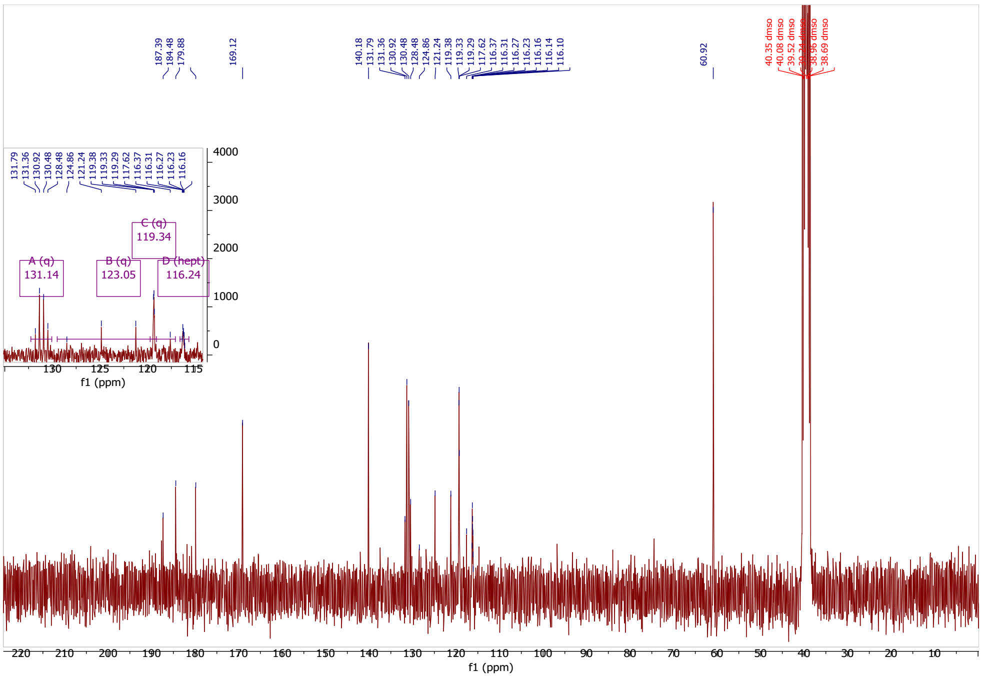

#### Synthesis of Man(α1,2)Man-PEG3-amine (38)

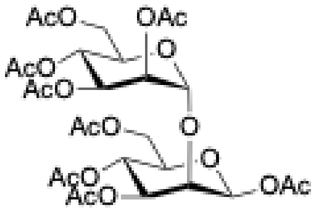

**35**

Commercially available 1,3,4,6-tetra-O-acetyl-β-D-mannose (1.70 g, 4.88 mmol) and trichloroacetimidate 2,3,4,6-tetra-O-acetyl-β-D-mannopyranose (3.12 g, 6.33 mmol) were dissolved in dry DCM (48 mL) and 4 Å molecular sieves were added. The mixture was stirred at 0 °C for 15 min, then TMSOTf (264.5 µL, 1.46 mmol) was added dropwise. The end of the reaction was determined after 30 min by TLC (4:6, Hex/EtOAc). The reaction solution was washed with sat. $NaHCO_3$ and the phases were separated. The aqueous phase was extracted with DCM. The combined organic phases were dried with $MgSO_4$ and the solvent removed under vacuum. The product was purified by column chromatography to yield **35** (3.19 g, 86%) as a white foam.

$^{1}H$ NMR (400 MHz, $CDCl_3$): δ 2.03, 2.04, 2.05, 2.09, 2.11, 2.13, 2.16, 2.17 (s, 3H, 8 acetyl groups), 3.80 (m, J=2.40 Hz, H5s), 4.04-4.36 (m, J=7.11 Hz, 4H6s, H2s), 4.43 (m, J=2.37 Hz, H5s), 5.01 (d, J=1.76 Hz, H1s), 5.12 (q, J=4.19 Hz, H3s), 5.30-5.40 (m, H2s, 2H4s), 5.50 (q, J=4.44 Hz, H3s), 5.80 (d, J=0.92 Hz, H1s).

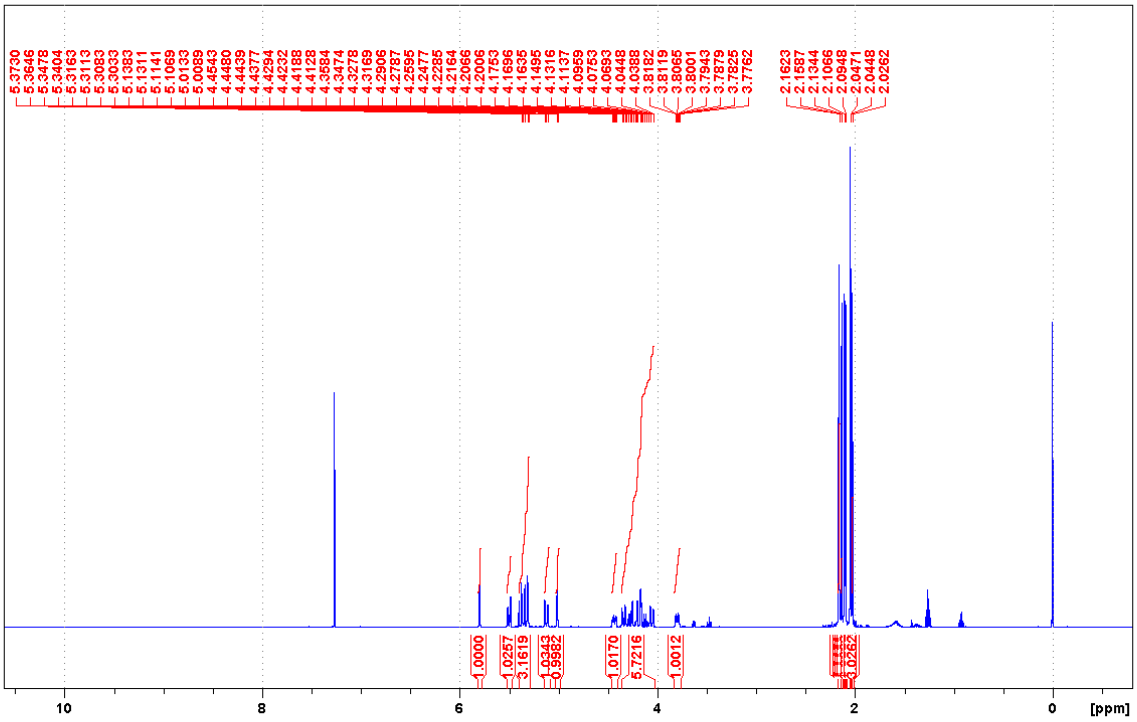

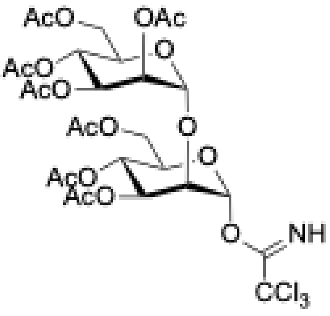

**36**

The acetyl group in the anomeric position was removed as previously described [16]. Briefly, compound **36** (3.19 g, 4.70 mmol) was dissolved in THF (40 mL) to which benzylamine (0.55 g, 5.13 mmol) was added and allowed to react under stirring overnight at r.t. The reaction mixture was then diluted with EtAcO and washed with HCl 1M. Organic phase was separated, dried over anhydrous $MgSO_4$, filtered and concentrated. The resulting product (2.06 g, 69%) was then purified by chromatography column (3:7, Hex/EtAcO) as a white foam. It (2.06 g, 3.24 mmol) was then dissolved in dry DCM (25 mL), to which it was added trichloroacetimidate (2.27 mL, 22.6 mmol) and DBU (338 μL, 2.26 mmol) at 0ºC under $N_2$ atmosphere. After allowing the reaction mixture to warm up till r.t., stirring was maintained for 2 h. The solvent was then removed, and the crude was purified by column chromatography (4:6, Hex/EtAcO) to give compound **36** (1.60 g, 63%) as a white foam.

$^1$H NMR (400 MHz, $CDCl_3$): δ 2.16, 2.14, 2.10, 2.08, 2.06, 2.05, 2.02 (s, 3H, 7 acetyl groups), 4.13-4.30 (m, 7 H, 4H6s, H2s, 2H5s), 5.00 (d, J=1.72 Hz, H1s), 5.28-5.50 (m, 5 H, 2H3s, 2H4s, H2s), 6.43 (d, J=2.04 Hz, H1s), 8.72 (s, 1 H, NH).

$^{13}$C NMR (101 MHz, $CDCl_3$): δ 20.62, 20.64, 20.64, 20.69, 20.69, 20.86, 20.86 ($CH_3$ acetyl groups), 61.70, 62.29 (C6s), 65.30, 66.15 (C4s), 68.37 (C2s), 69.54, 69.67 (C3s), 69.85, 71.29 (C5s). 74.96 (C2s), 90.52 (CCl3), 95.56 (C1s), 99.22 (C1s), 160.12 (CNH), 169.26, 169.48, 169.71, 169.90, 170.32, 170.62, 170.82 (CO).

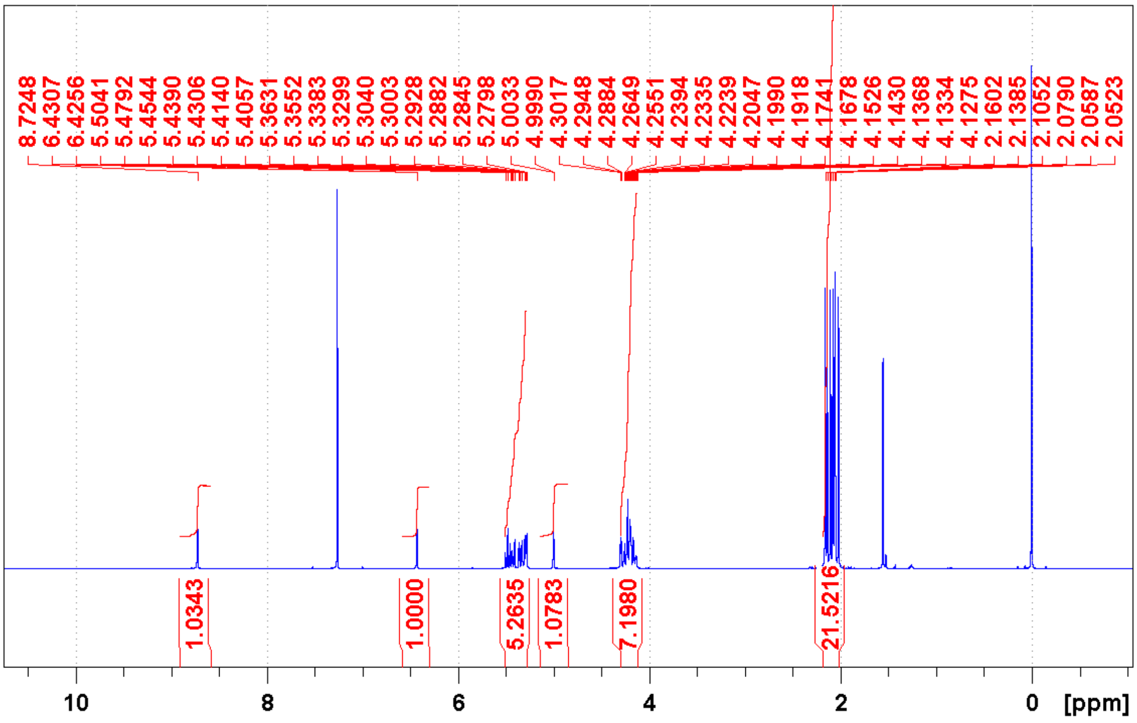

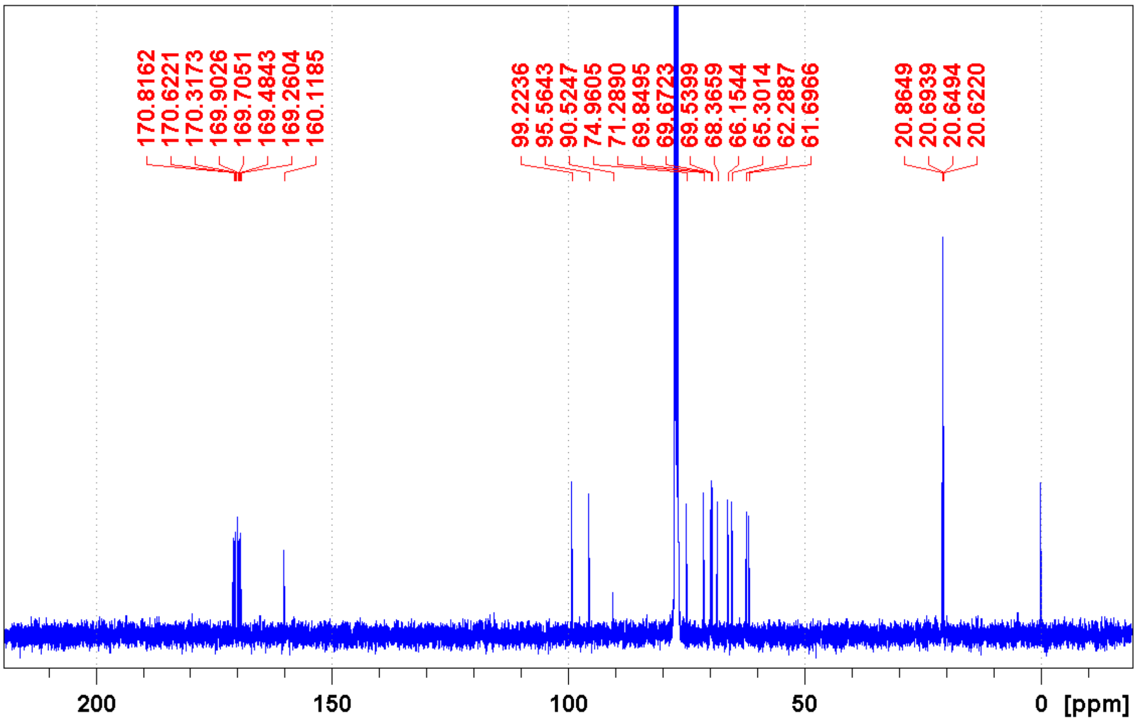

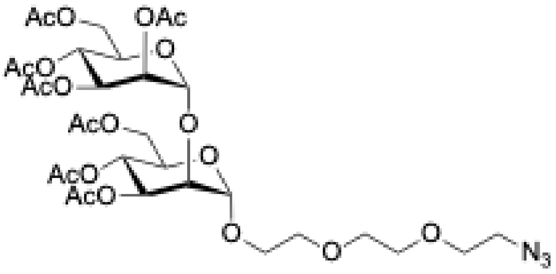

**37**

Derivative **36** (300 mg, 0.38 mmol) and commercially available $N_3$-PEG3-OH (101 mL, 0.08 mmol) were dissolved in dry DCM (4 mL) and 4 Å molecular sieves were added. The mixture was taken from r.t. to 0º C and stirred at 0 °C for 10 min, then TMSOTf (14 mL, 1.46 mmol) was added dropwise. The end of the reaction was determined after 1 h by TLC (4:6, Hex/EtOAc). The reaction solution was concentrated. The product was purified by column chromatography to yield **37** (300 mg, 98%).

$^1$H NMR (400 MHz, $CDCl_3$): δ 2.01, 2.03, 2.04, 2.08, 2.09, 2.15, 2.16 (s, 3H, 7 acetyl groups), 3.63 (t, J=4.52 Hz, $CH_2$-PEG), 3.68-3.75 (m, 2H6s, 4x$CH_2$-PEG, 2H5s, 2H4s), 3.83 (m, 1 H, $CH_2$-PEG), 4.00 (m, 1 H, $CH_2$-PEG), 4.06-4.25 (m, 2H6s, H2s, 2H3s), 4.93 (s, H1s), 4.99 (s, H1s), 5.26-5.43 (m, H2s, 2H6s, 2H3s).

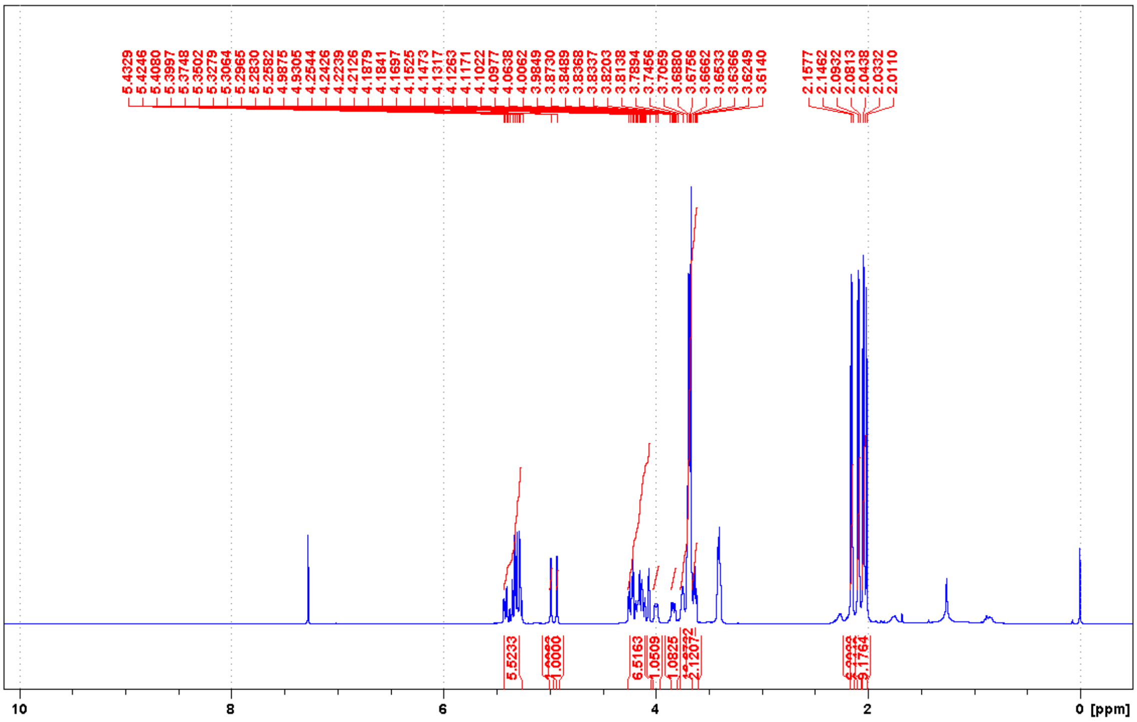

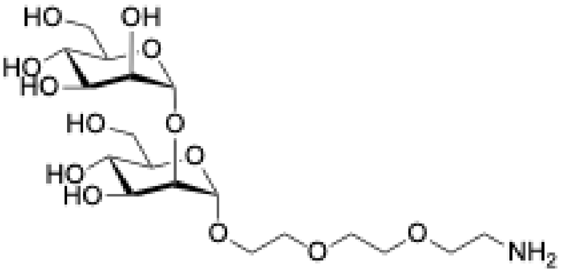

**38**

Acetyl protecting groups were removed by dissolving compound **37** (300 mg, 0.38 mmol) in 10 mL of MeOH. MeONa was added dropwise until pH 9 was reached. After stirring for 2 h at r.t., Amberlite $H^+$ resin was added. Completion of the reaction was checked by TLC (95:5, DCM/MeOH), subsequently the reaction mixture was filtered, concentrated and the deprotected molecule (184 mg, 97%) was used in the next step without further purification. Over 100 mg of preactivated Pd-C in 5 mL under $H_2$ atmosphere the intermediate was added dissolved in 5 mL of MeOH with 3-5 drops of AcOH. The reaction was kept under stirring overnight and then purified through C-18 cartridge to give pure final product **38** (155 mg, 89%) as a white powder.

$^{1}H$ NMR (400 MHz, $D_2O$): δ 3.21 (t, J=5.10 Hz, C$\underline{H}_2$-$NH_2$), 3.62-3.80 (m, 2H6s, 9H-($CH_2$-PEG), 2H5s, 2H4s) 3.84-3.95 (m, 2H6s, H-($CH_2$-PEG), 2H3s), 4.00 (q, J=1.65 Hz, H2s), 4.08 (q, J=1.69 Hz, H2s), 5.04 (d, J=1.48 Hz, 1Hs), 5.14 (d, J=1.40 Hz, H1s).

$^{13}C$ NMR (101 MHz, $D_2O$): δ 39.12 ($\underline{C}H_2$-$NH_2$), 60.26 ($CH_2$-PEG), 60.90, 61.13 (C6s), 66.42 ($CH_2$-PEG), 66.87, 66.90 (C4s), 69.49, 69.64 ($CH_2$-PEG), 69.91 (C2s), 70.14, 70.27 (C3s), 71.67 (C-PEG) 72.77, 73.26 (C5s), 78.64 (C2s), 98.34, 102.32 (C1s).

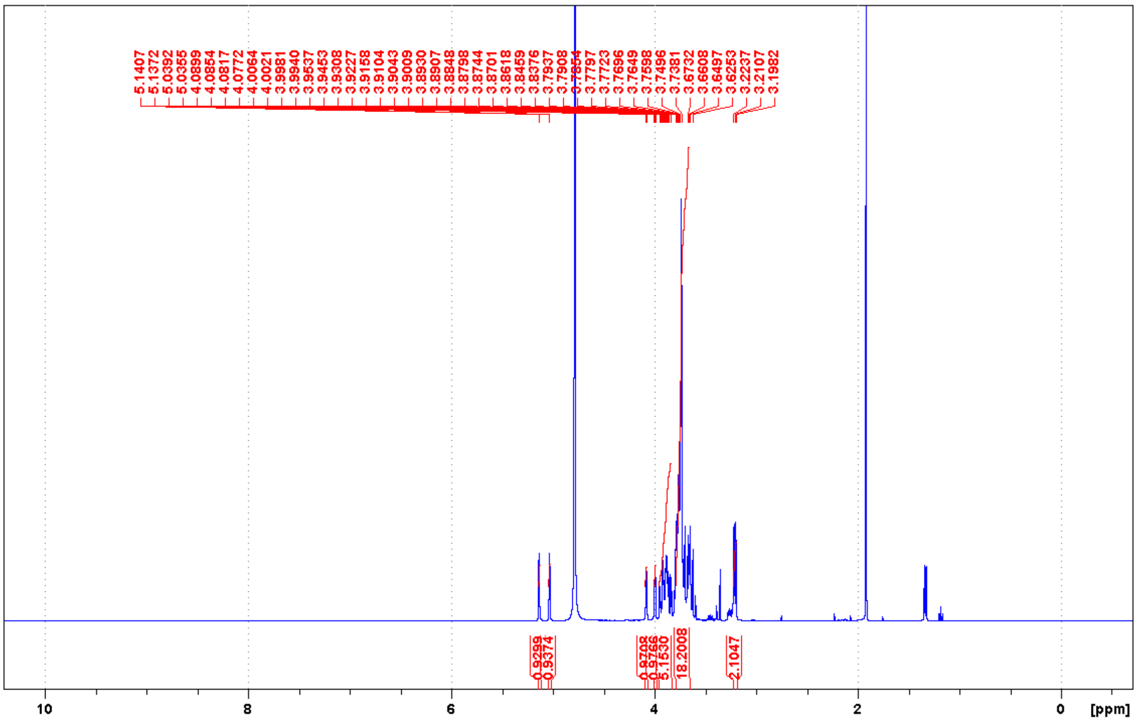

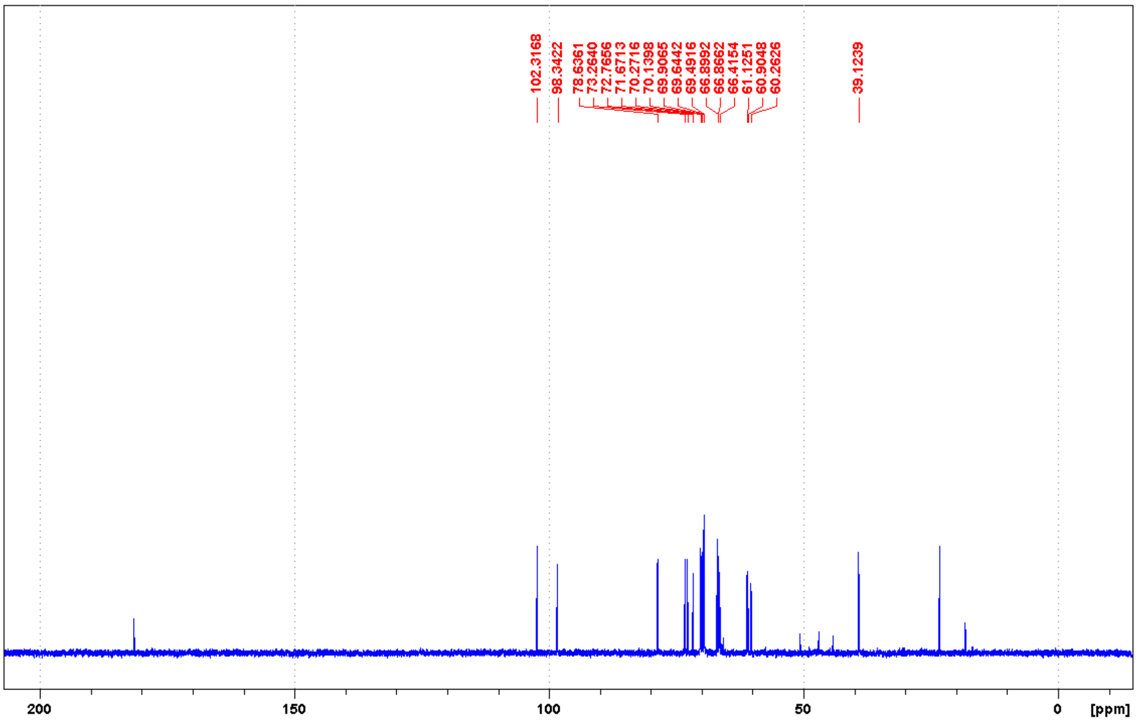

#### Library characterization

##### *N*-Acetyl-lysine reactivity assay based on HPLC-MS

HPLC-MS measurements were performed using a Shimadzu LCMS-2020 device equipped a positive–negative double ion source (DUIS±) and a quadrupole MS analyzer in the range of *m/z* 50–1000. The sample was eluted with gradient elution using eluent A (0.1% FA in $H_2O$) and eluent B (0.1% FA in ACN). The column temperature was always kept at 30 °C; the injection volume was 10 µL, and the flow rate was set to 1.5 mL/min. A Reprospher C18 (5 µm, 100 mm × 3 mm) column was used along with the following gradient. The initial condition was 0% B eluent, followed by a linear gradient to 100% B eluent by 1 min; from 1 to 3.5 min, 100% B eluent was retained. From 3.5 to 4.5 min, the initial condition with 5% B eluent was restored and retained until 5 min.

For the reactivity and stability assay, in a glass LC-MS vial the internal standard (indoprofen or papaverine or 1,4-dicyanobenzene, 50 µl, 2 mM in acetonitrile) and the 100 mM BBS buffer solution (900 µl, pH 10.2) was mixed with or without 50 mM *N*-acetyl-lysine (providing results of reactivity or stability, respectively). Finally, to this mixture the corresponding electrophile was added (50 µl, 20 mM in acetonitrile) right before the measurement. The vial was mixed, then analyzed by HPLC-MS (10 µl injection volume) at intervals of approximately 60 min (0-4h, 6h, 8h, 12h, 24h). Final volume was 1 ml. In the case of highly reactive compounds, the measurement was done within 1 h, monitored every 5 minutes. The AUC (area under the curve) values were determined via integration of HPLC or MS chromatograms and then corrected with the internal standard. The fragments' AUC values were subjected to ordinary least-squares (OLS) linear regression, and to compute the important parameters (kinetic rate constant and half-life time), an Excel sheet was applied. The data are expressed as means of duplicate determinations. The kinetic rate constant for the degradation and corrected *N*-acetyl-lysine reactivity were calculated as follows. The reaction half-life for pseudo-first-order reactions ($t_{1/2}$) is ln 2/k, where k is the reaction rate. In the case of competing reactions (reaction with *N*-acetyl-lysine and degradation), the apparent reaction rate is $k_{Lys} = k_{Blank} + k_{Binding}$. When half-lives are measured experimentally, $t_{1/2}(Lys) = \ln 2/(k_{Lys}) = \ln 2/(k_{Blank} + k_{Binding})$. In our case, the corrected $k_{Blank}$ and $k_{Lys}$ (regarding blank and *N*-acetyl lysine-containing samples, respectively) can be calculated by linear regression of the measured kinetic data points. The corrected $k_{Binding}$ is calculated as $k_{Lys} – k_{Blank}$, and finally, the half-life is determined using the equation $t_{1/2} = \ln 2/k$.

**Assessment of stability at pH 7.4**

Next, we characterized the intrinsic stability of the compounds in HEPES buffer (pH 7.4). A 1 mM solution of the fragment in HEPES buffer (pH 7.4) with 10 % acetonitrile with a 0.1 mM solution of indoprofen or papaverine as the internal standard was incubated (providing results of stability). The reaction mixture was analyzed by HPLC-MS sampling after 0, 1 and 20 h, respectively. The AUC (area under the curve) values were determined via integration of HPLC or MS chromatograms and then corrected with the internal standard.

### Supporting Figures

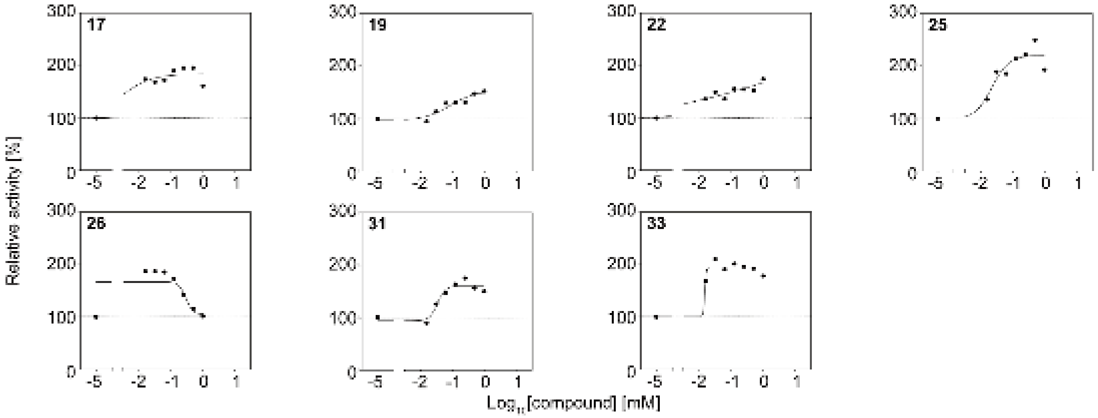

**Figure S1. HRP assay dose-response curves for compounds showing ≥ 50% activation.** Dose-response of compounds showing over 50% activation of DC-SIGN binding to HRP in the primary screening. Dose-response curves of compound **22** could not be fitted. Compound **26** inhibits at higher concentrations and could not be fitted. Other hit compounds showed micromolar $EC_{50}$ values: **17** (2.8 µM), **19** (84 µM), **22** (no fit), **25** (23 µM), **26** (no fit), **31** (35 µM), **33** (15 µM).

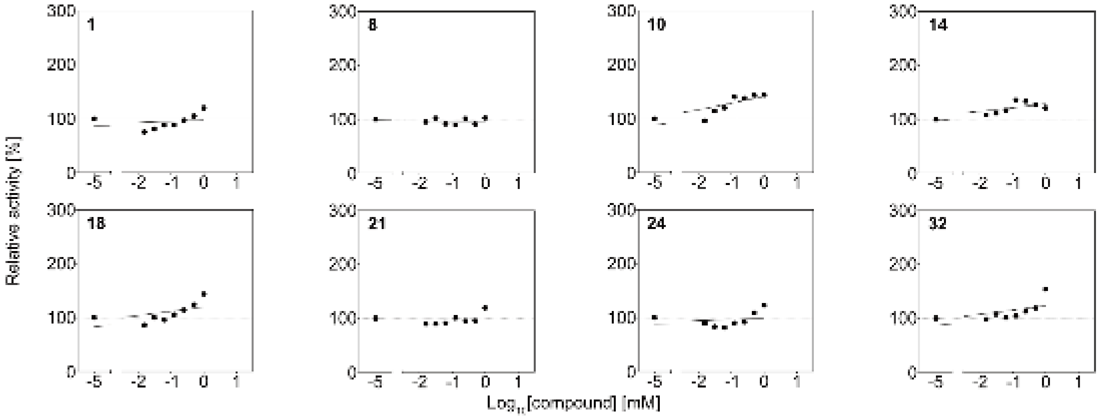

**Figure S2. HRP assay dose-response curves for compounds showing ≥10% activation.** Dose-response of compounds showing over 10% inhibition of DC-SIGN binding to HRP in the primary screening. No compound apart from **30** (Figure 2c) could be validated as inhibitor based on dose-response.

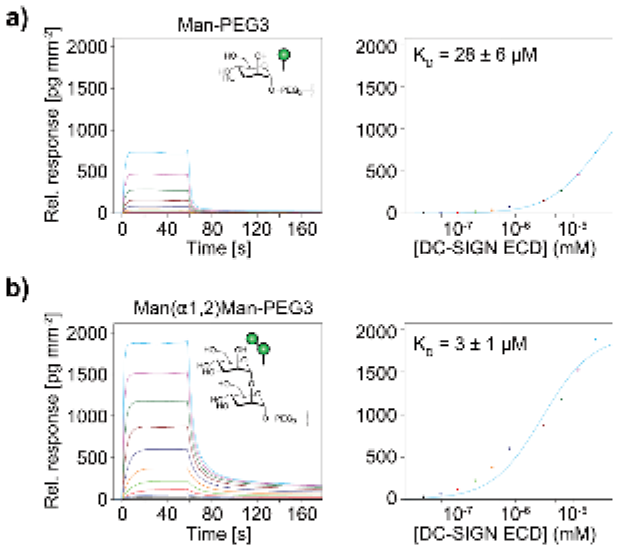

**Figure S3. Interaction of DC-SIGN ECD with the glycosylated GCI chip.** Sensorgrams of the titration of DC-SIGN ECD over the mannose-PEG3 **(a)** and Man(α1,2)Man-PEG3 **(b)** coupled channels of the GCI chip (left). Equilibrium fitting of the dissociation constant revealed micromolar affinities for both channels (right).

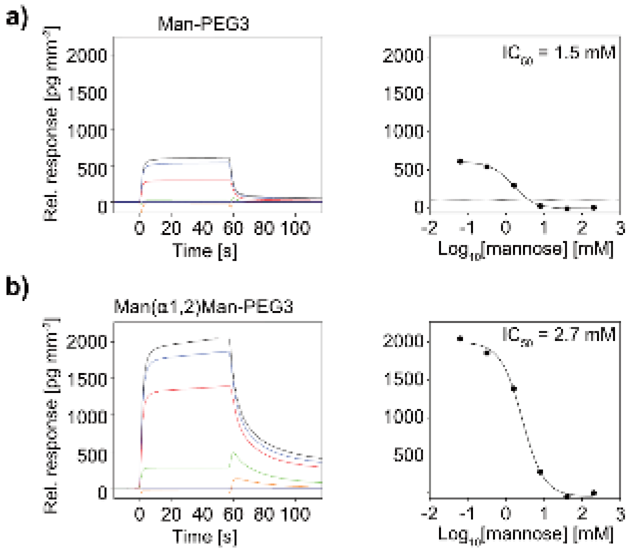

**Figure S4. Inhibition of DC-SIGN ECD interacting with the glycosylated GCI chip.** Sensorgrams of the titration of mannose at a constant concentration of DC-SIGN ECD over the mannose-PEG3 **(a)** and Man(α1,2)Man-PEG3 **(b)** modified channels of the GCI chip (left). Millimolar $IC_{50}$ values confirm carbohydrate-specificity of the interaction with the chip (right).

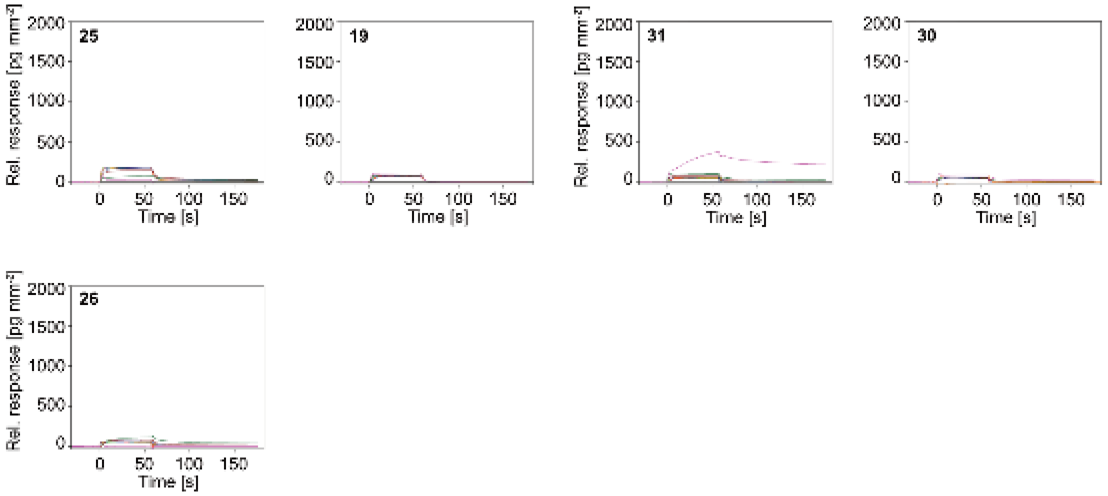

**Figure S5. Sensorgrams of excluded compounds.** Sensorgrams of DC-SIGN ECD interacting with the mannose-PEG3-modified channel after modification with compounds excluded from further experiments.

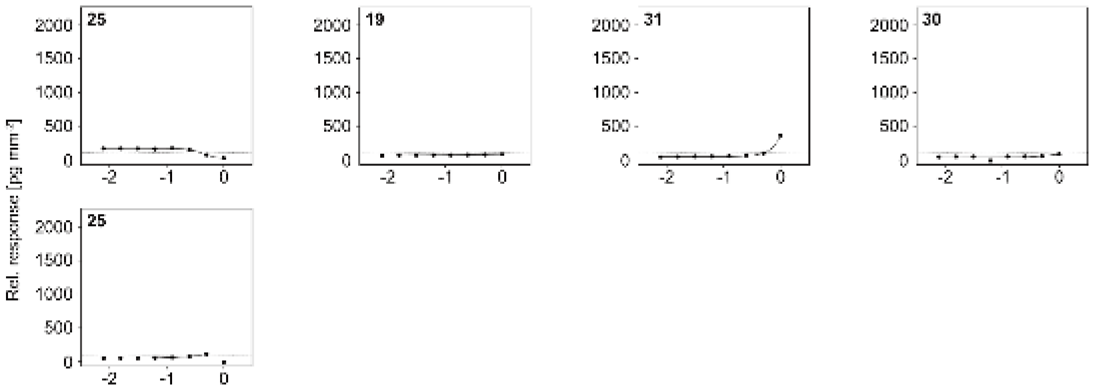

**Figure S6. Dose-response curves of excluded compounds.** Dose-response curves for sensorgrams shown in figure S5 of compounds without observable dose-response in the GCI assay with DC-SIGN ECD.

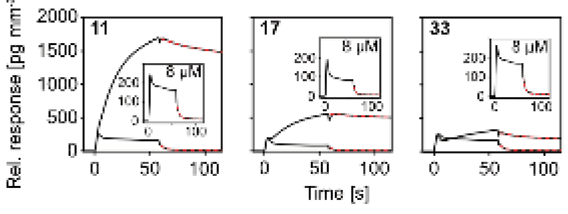

**Figure S7. Covalent hits reduce the off-rate of the DC-SIGN ECD-carbohydrate interaction.** Off-rates ($k_{off}$) of the ECD-mannose-PEG3 interaction are drastically reduced for **11**, **17** and **33**-modified DC-SIGN ECD, when comparing low (0.008 mM; $k_{off,\mathbf{11}} \approx 0.20\ s^{-1}$, $k_{off,\mathbf{17}} \approx 0.27\ s^{-1}$ $k_{off,\mathbf{33}} \approx 0.20\ s^{-1}$) and high (0.25 mM; $k_{off,\mathbf{11}} \approx 0.02\ s^{-1}$, $k_{off,\mathbf{17}} \approx 0.02\ s^{-1}$ $k_{off,\mathbf{33}} \approx 0.03\ s^{-1}$) concentrations of the compounds.

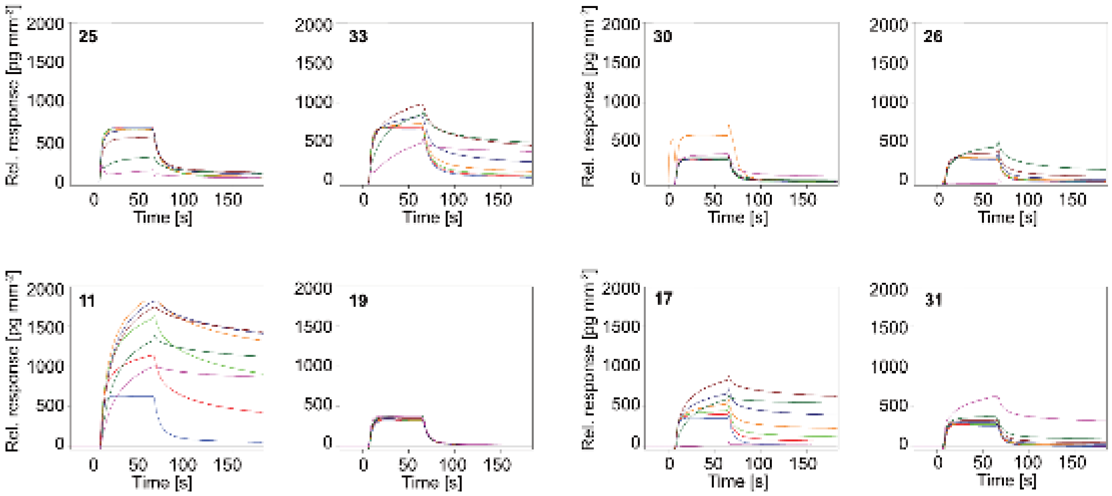

**Figure S8. Sensorgrams of compounds tested on the Man(α1,2)Man-PEG3-modified chip.** Sensorgrams of DC-SIGN ECD interacting with the Man(α1,2)Man-PEG3-modified channel after modification with compounds all hit compounds. Overall, similar results as for the mannose-PEG3-modified channel were obtained (Figures 2e, f, S5 and S6).

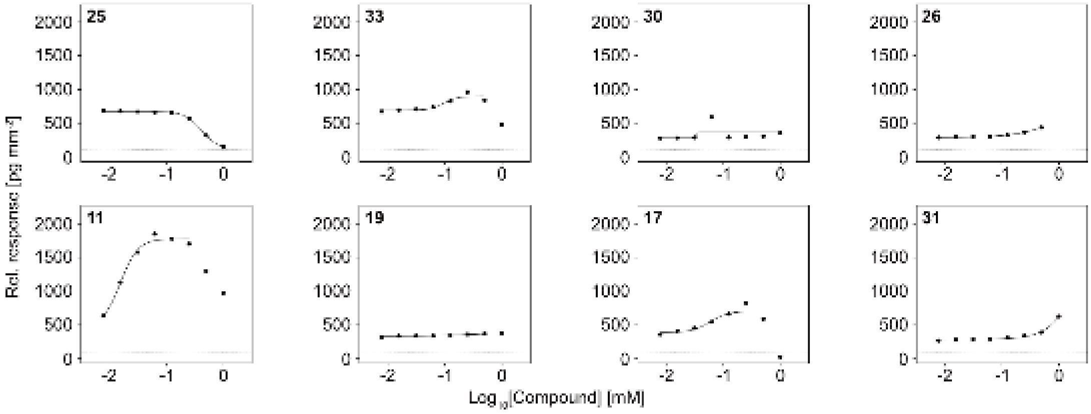

**Figure S9. Dose-response curves of compounds tested on the Man(α1,2)Man-PEG3-modified chip.** Dose-response curves for sensorgrams shown in figure S8. Similar to the results obtained from mannose-PEG3-modified channel, only compounds **33** ($EC_{50}$ = 96 µM, hill slope = 2.8), **11** ($EC_{50}$ = 16 µM, hill slope = 3.7) and **17** ($EC_{50}$ = 58 µM, hill slope = 2.7). For other hit compounds showed dose-response curves could not be fitted.

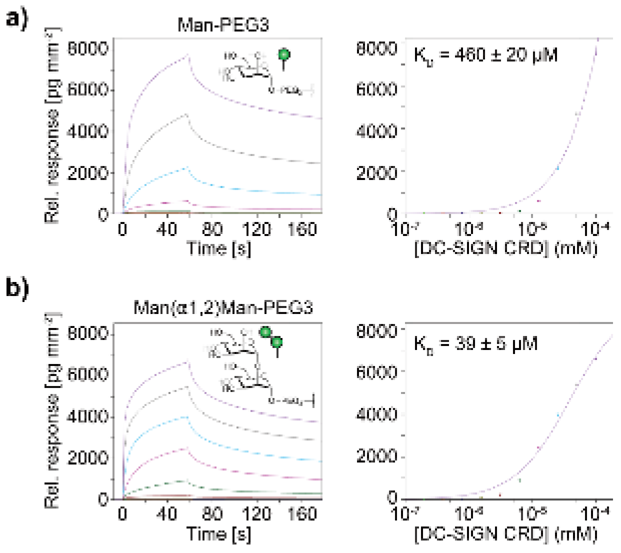

**Figure S10. Interaction of DC-SIGN CRD with the glycosylated GCI chip.** Sensorgrams of the titration of DC-SIGN CRD over the mannose-PEG3 **(a)** and Man(α1,2)Man-PEG3 **(b)** coupled channels of the GCI chip (left). Equilibrium fitting of the dissociation constant reveals micromolar affinities for both channels (right).

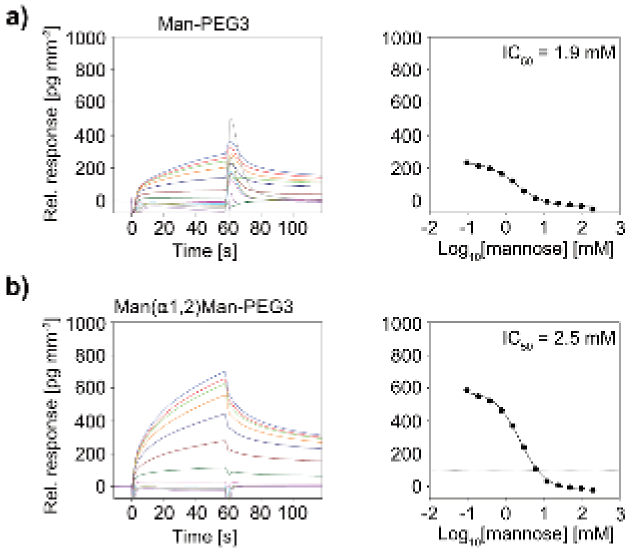

**Figure S11. Inhibition of DC-SIGN CRD interacting with the glycosylated GCI chip.** Sensorgrams of the titration of mannose at a constant concentration of DC-SIGN CRD over the mannose-PEG3 **(a)** and Man(α1,2)Man-PEG3 **(b)** modified channels of the GCI chip (left). Millimolar $IC_{50}$ values confirm carbohydrate-specificity of the interaction with the chip (right).

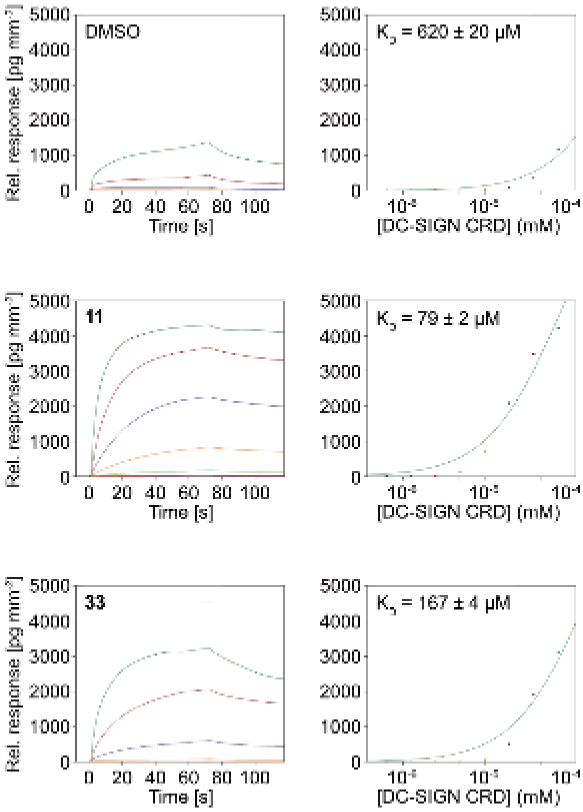

**Figure S12. Interaction of 11 and 33-modified DC-SIGN CRD with the mannose-PEG3-modified GCI chip.** Sensorgrams of the titration of unmodified (DMSO), **11** or **33**-modified DC-SIGN CRD over the mannose-PEG3 coupled channel of the GCI chip (left). Equilibrium fitting of the dissociation constant reveals an 8-fold increase in affinity for **11**-modified CRD and a 4-fold increase for **33**-modified CRD compared to the DMSO control.

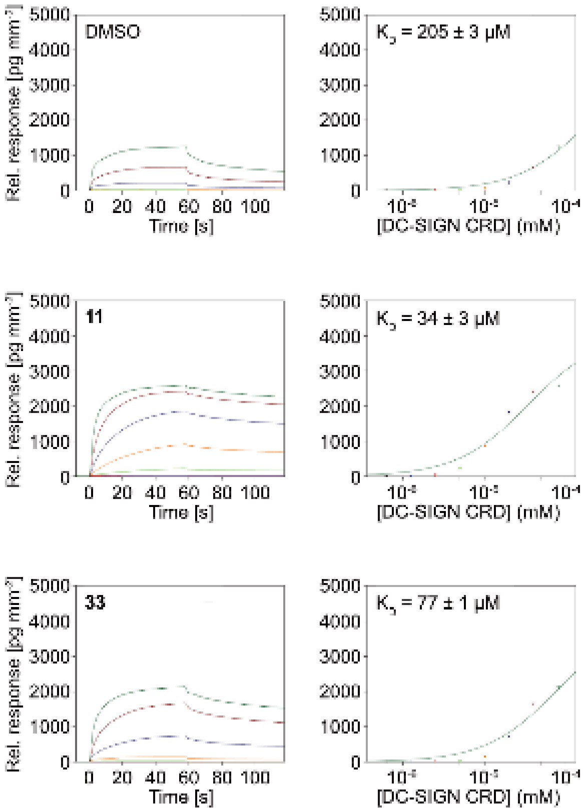

**Figure S13. Interaction of 11 and 33-modified DC-SIGN CRD with the Man(α1,2)Man-PEG3 GCI chip.** Sensorgrams of the titration of unmodified (DMSO), **11** or **33**-modified DC-SIGN CRD over the mannose-PEG3 coupled channel of the GCI chip (left). Equilibrium fitting of the dissociation constant revealed an 8-fold increase in affinity for **11**-modified CRD and a 4-fold increase for **33**-modified CRD compared to the DMSO control (right).

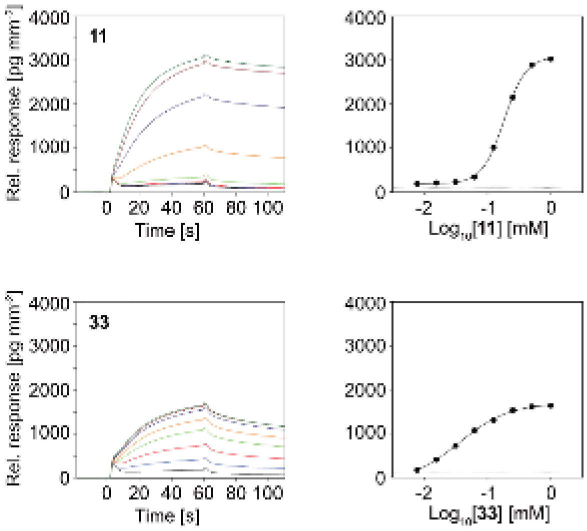

**Figure S14. Dose-response for compounds 11 and 33 obtained from the Man(α1,2)Man-PEG3-modified channel.** Sensorgrams of the titration **11** and **33** to DC-SIGN CRD on the Man(α1,2)Man-PEG3-modified channel of the GCI chip (left). Dose-response curves for sensorgrams reveal activities ($EC_{50,\mathbf{33}}$ = 183 µM, hill slope = 2.5; $EC_{50,\mathbf{33}}$ = 36 µM, hill slope = 1.1).

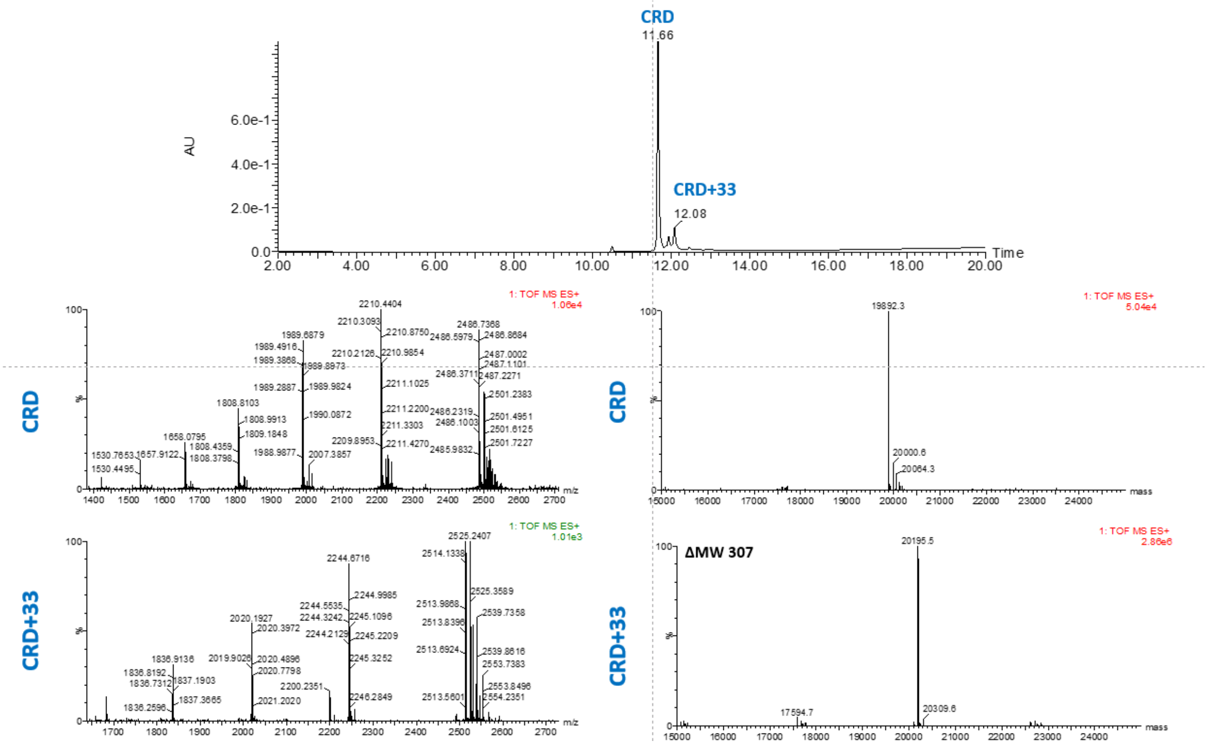

**Figure S15. PDA chromatogram and LC-MS spectrum for 33-modified DC-SIGN CRD.**

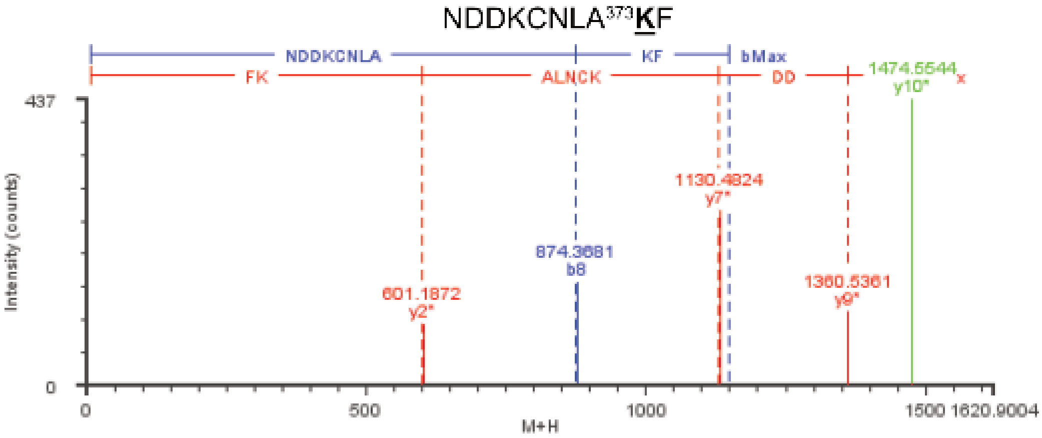

**Figure S16. LC-MS/MS analysis of a 33-modified DC-SIGN CRD peptide.** Annotated MS/MS spectrum of a **33**-modified DC-SIGN CRD peptide after digestion with chymotrypsin suggest K373 to be modified. Fragment ions with asterisk denote modified MS/MS fragment ions. Analytical data is shown in table S4.

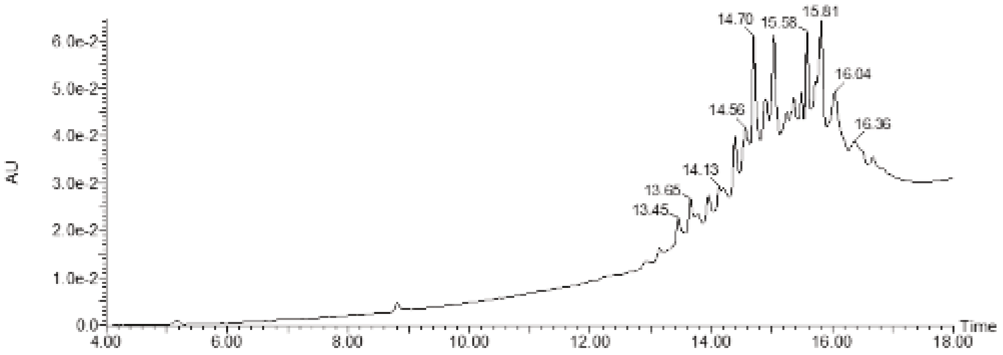

**Figure S17. PDA chromatogram for 11-modified DC-SIGN CRD.**

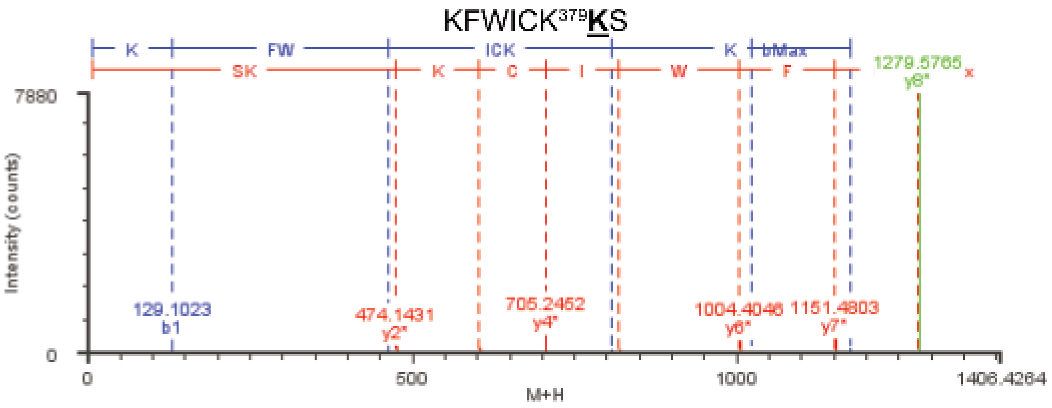

**Figure S18. LC MS/MS spectrum for 11-modified DC-SIGN CRD.** Annotated MS/MS spectrum of a **11**-modified DC-SIGN CRD peptide after digestion with ProAlanase suggest K379 to be modified. Fragment ions with asterisk denote modified MS/MS fragment ions. Analytical data is shown in table S5.

### Supporting Tables

**Table S1.** Structures and warhead chemistries of the electrophilic fragment library.

| ID | Structure | Warhead | Reactivity modes |
|---|---|---|---|
| **1** | 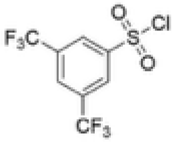 | Sulfonyl chloride | substitution at S |
| **2** | 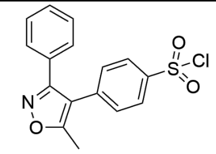 | Sulfonyl chloride | substitution at S |
| **3** | 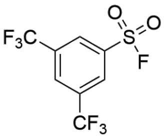 | Sulfonyl fluoride | substitution at S |
| **4** | 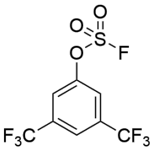 | Fluorosulfate | substitution at S |
| **5** | 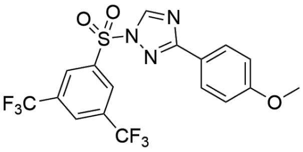 | Sulfonyl triazole | substitution at S |
| **6** | 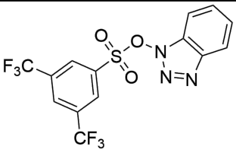 | *N*,*O*-Sulfonate | substitution at S |
| **7** | 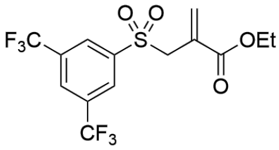 | Sulfonyl acrylate | substitution at C |
| **8** | 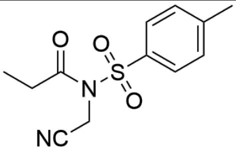 | *N*-Acyl-*N*-alkyl sulfonamide (NASA) | substitution at C |
| **9** | 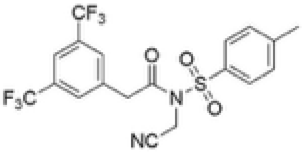 | *N*-Acyl-*N*-alkyl sulfonamide (NASA) | substitution at C |
| **10** | 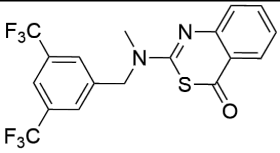 | Benzothiazinone | substitution at C |

| ID | Structure | Warhead | Reactivity modes |
|---|---|---|---|
| **11** | 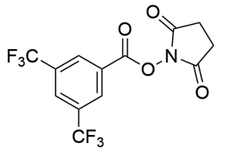 | *N*-Hydroxysuccinimidyl ester | acylation (substitution at C) |
| **12** | 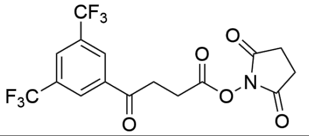 | *N*- Hydroxysuccinimidyl ester | acylation (substitution at C) |
| **13** | 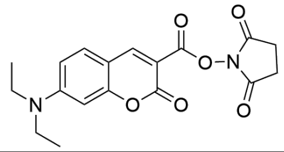 | *N*- Hydroxysuccinimidyl ester | acylation (substitution at C) |
| **14** | 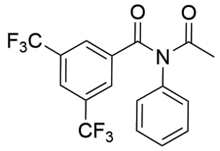 | Bisacetanilide | substitution at C |
| **15** | 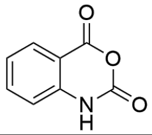 | Isatoic anhydride | substitution at C |
| **16** | 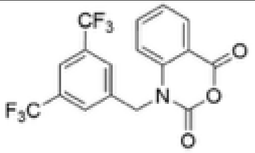 | Isatoic anhydride | substitution at C |
| **17** | 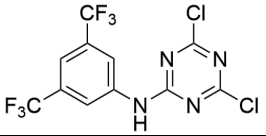 | Dichlorotriazine | substitution at C |
| **18** | 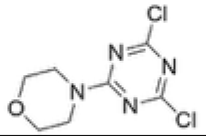 | Dichlorotriazine | substitution at C |
| **19** | 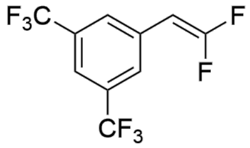 | Difluorostyrene | substitution at C |
| **20** | 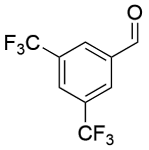 | Aldehyde | Direct addition |
| **21** | 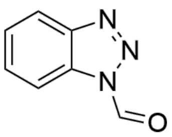 | *N*-formyl heterocycle | Direct addition |
| **22** | 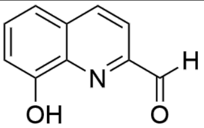 | Heterocyclic aldehyde | Direct addition |

| ID | Structure | Warhead | Reactivity modes |
|---|---|---|---|
| **23** | 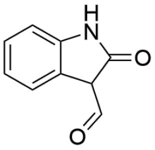 | Aldehyde | Direct addition |
| **24** | 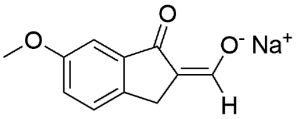 | Ketoenolate | Direct addition |
| **25** | 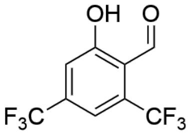 | Hydroxy aldehyde (salicylaldehyde) | Direct addition |
| **26** | 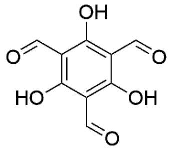 | Acylphloroglucinol (salicylaldehyde) | Direct addition |
| **27** | 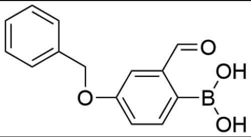 | Formylphenyl boronic acid | Direct addition |
| **28** | 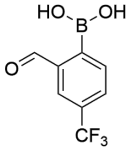 | Formylphenyl boronic acid | Direct addition |
| **29** | 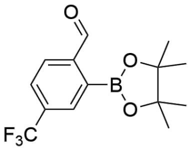 | Formylphenyl boronic ester | Direct addition |
| **30** | 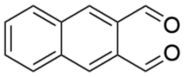 | *ortho*-Phthalaldehyde (OPA) | Direct addition |
| **31** | 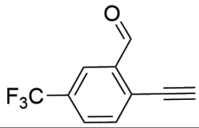 | Ethynyl aldehyde/formylacetylene | Direct addition |
| **32** | 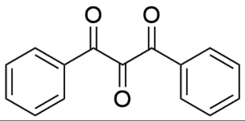 | Polycarbonyl | Direct addition |
| **33** | 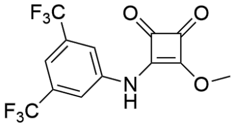 | Squarate | conjugate addition-elimination (substitution at C) |
| **34** | 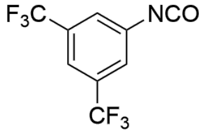 | Isocyanate | Direct addition |

**Table S2.** Pseudo first-order kinetic constants and half-life of compounds at pH 10.2 and stability at pH 7.4.

| ID | $k_{Lys}$ pH 10.2 [$h^{-1}$] | $k_{blank}$ pH 10.2 [$h^{-1}$] | Binding pH 10.2 [$h^{-1}$] | Reactivity $t_{1/2}$ pH 10.2 [h] | Stability $t_{1/2}$ pH 10.2 [h] | Stability 1h pH 7.4 [%] | Stability 20h pH 7.4 [%] |
|---|---|---|---|---|---|---|---|
| **1** | - | - | - | <1min | <1min | 8% | 0% |
| **2** | 3.6 | 1.9 | 1.7 | 0.4 | 0.4 | 66% | 1% |
| **3** | - | 27.2 | - | <1min | 0 | 98% | 17% |
| **4** | 1.4 | 0.1 | 1.2 | 0.6 | 4.9 | 65% | 9% |
| **5** | 1.3 | 0.6 | 0.6 | 1.1 | 1.1 | 93% | 38% |
| **6** | - | 2 | - | <1min | 0.3 | 10% | 0% |
| **7** | 29.9 | 0.5 | 29.4 | 0 | 1.5 | 87% | 44% |
| **8** | 50 | 14.7 | 35.3 | 0 | 0 | 80% | 68% |
| **9** | 6.6 | 4 | 2.6 | 0.3 | 0.2 | 60% | 1% |
| **10** | 0 | 0 | 0 | 78 | 23.6 | 98% | 1% |
| **11** | - | - | - | <1min | <1min | 52% | 0% |
| **12** | - | - | - | <1min | <1min | 35% | 0% |
| **13** | - | 8.4 | - | <1min | 0.1 | 1% | 0% |
| **14** | 3.1 | 0.5 | 2.6 | 0.3 | 1.4 | 45% | 3% |
| **15** | - | 2.1 | - | <1min | 0.3 | 41% | 0% |
| **16** | - | 26.5 | - | <1min | 0 | 51% | 1% |
| **17** | 8.9 | 0.7 | 8.1 | 0.1 | 1 | 95% | 5% |
| **18** | 41.1 | 0.1 | 41 | 0 | 6.1 | 86% | 73% |
| **19** | 0.7 | 0.2 | 0.5 | 1.3 | 3.3 | 96% | 2% |
| **20** | 0 | 0.1 | - | 24h + | 5.7 | 83% | 20% |
| **21** | 0 | 0 | 0 | 27.7 | 150.4 | 82% | 81% |
| **22** | 0 | 0 | - | 24h + | 113.7 | 94% | 35% |
| **23** | 0 | 0 | 0 | 65.8 | 154.2 | 97% | 94% |
| **24** | 0 | 0 | - | 24h + | 35.4 | 100% | 97% |
| **25** | 0 | 0 | 0 | 40.9 | 246.8 | 96% | 30% |
| **26** | 15.8 | 0 | 15.8 | 0 | 20.5 | 86% | 74% |
| **27** | 0.1 | 0.1 | 0 | 35.4 | 10.8 | 97% | 90% |
| **28** | 0 | 0 | 0 | 110.5 | 447.2 | 97% | 95% |
| **29** | - | - | - | - | <1min | 19% | 0% |
| **30** | - | 0.1 | - | <1min | 11.7 | 82% | 60% |
| **31** | 3 | 0 | 3 | 0.2 | 15.4 | 98% | 42% |
| **32** | - | - | - | - | <1min | 66% | 6% |
| **33** | 6.1 | 0.1 | 6.1 | 0.1 | 12.2 | 34% | 13% |
| **34** | - | - | - | <1min | <1min | 73% | 39% |

**Table S3.** Surface area and $pK_a$ values of lysines in each ECD monomer.

| Lysine # | Type[a] | SASA (monomer) Å | SASA (tetramer) Å | $pK_a$ (monomer) | $pK_a$ (tetramer) |
|---|---|---|---|---|---|
| K285 | surface* | 145.9 | 63.01 | 8.0 | 8.0 |
| K295 | surface | 115.1 | 115.05 | 10.1 | 10.1 |
| K340 | surface | 108.0 | 107.98 | 10.5 | 10.5 |
| K368 | surface | 131.1 | 131.12 | 10.4 | 10.4 |
| K373 | buried** | 28.9 | 28.90 | 11.8 | 10.6 |
| K378 | buried* | 18.6 | 17.45 | 11.6 | 11.5 |
| K379 | surface* | 96.1 | 38.88 | 10.2 | 10.2 |

[a] *lysine at dimerization interface, ** lysine at carbohydrate binding site

**Table S4.** Analytical data of a modified peptide detected in the peptide mapping analysis of 33-modified DC-SIGN CRD using chymotrypsin digestion

| Peptide | Retention time (Min) | Calculated Peptide Mass (Da) | Measured *m/z* | Mass Error (ppm) | Measured Mass (Da) |
|---|---|---|---|---|---|
| NDDKCNLAKF | 14.5 | 1473.5458 | 737.7805 | -0.5 | 1473.545 |

**Table S5.** Analytical data of a modified peptide detected in the peptide mapping analysis of 11-modified DC-SIGN CRD using ProAlanase digestion

| Peptide | Retention time (Min) | Calculated Peptide Mass (Da) | Measured *m/z* | Mass Error (ppm) | Measured Mass (Da) |
|---|---|---|---|---|---|
| KFWICKKS | 13.8 | 1278.5695 | 640.2934 | 1.1 | 1278.5709 |
